# Global analysis of biosynthetic gene clusters reveals conserved and unique natural products in entomopathogenic nematode-symbiotic bacteria

**DOI:** 10.1101/2022.01.21.477171

**Authors:** Yi-Ming Shi, Merle Hirschmann, Yan-Ni Shi, Shabbir Ahmed, Desalegne Abebew, Nicholas J. Tobias, Peter Grün, Jan J. Crames, Laura Pöschel, Wolfgang Kuttenlochner, Christian Richter, Jennifer Herrmann, Rolf Müller, Aunchalee Thanwisai, Sacha J. Pidot, Timothy P. Stinear, Michael Groll, Yonggyun Kim, Helge B. Bode

## Abstract

Microorganisms contribute to the biology and physiology of eukaryotic hosts and affect other organisms through natural products. *Xenorhabdus* and *Photorhabdus* (*XP*) living in mutualistic symbiosis with entomopathogenic nematodes produce a myriad of natural products to mediate bacteria–nematode–insect interactions. However, a lack of systematic analysis of the biosynthetic gene clusters (BGCs) has limited the understanding of how natural products justify the bacterial niche specificity. Here we combine pangenome and sequence similarity networks to analyze BGCs from 45 *XP* species. The identified 1,000 BGCs belong to 176 families, over half of which are unknown. Eleven BGCs represent the most conserved families. We then homologously express the ubiquitous and unique BGCs and identify compounds featuring unusual architectures. The bioactivity evaluation demonstrates that the prevalent compounds are eukaryotic proteasome inhibitors, insect virulence factors, or insect immune suppressors. These findings account for the functional basis of bacterial natural products in this tripartite relationship.

---

Interactions between microorganisms (e.g. bacteria) and higher eukaryotes are ubiquitous and have essential medical, environmental, and evolutionary significance^1^. Microorganisms supply nutrients^2^, shape immune systems^3^, maintain diverse and productive communities^4^, drive evolution^5^, and enhance ecological niches^6^ for higher eukaryotic hosts. Such microbe-host interactions can be relationships ranging from mutualistic/parasitic to pathogenic symbiosis^7^, in which microorganisms sense and respond to environmental changes with diffusible small molecules. These small molecules are also known as natural products or specialized metabolites, which affect not only the microbial host but also neighboring microbes and other organisms^8^. However, due to limitations in genetic tractability of microbial species, as well as formidable obstacles to imitating microbial natural habitats^9^, only a few correlations between microbial natural products (e.g. colibactin^10,11^ and tilivalline^12^ produced by bacteria in the human gut microbiota) and the function that microbial natural products endow the producers with have been characterized.

Entomopathogenic *Xenorhabdus* and *Photorhabdus* (*XP*) bacteria live in mutualistic symbiosis with nematodes of the genera *Steinernema* and *Heterorhabditis*, respectively. The dauer-stage nematodes carrying the symbiotic bacteria within their intestines actively search for insect larvae in the soil^13,14^, additionally sensing signals from plant roots infected by insects^15^. When nematodes invade insect prey through natural openings and cuticles, the bacterial symbionts are released into the insect hemolymph, where the bacteria begin to propagate and produce proteins (e.g. toxins and lytic enzymes) and natural products that help with killing the insect prey, degrading the insect cadaver, and protecting it against other soil-living organisms. The nematodes then feed on the predigested insect tissues, as well as *XP*, and reproduce within the cadaver. Upon food depletion, a new generation of dauer-stage nematodes re-associate with the symbiotic bacteria, exit the carcass, and seek new prey. Notably, although *XP* strains have yet to be found independently from environmental sources, they can be cultivated and genetically manipulated under standard laboratory conditions^13^. Also, the other two organisms, nematodes and insects, can be established readily in laboratory environments. Therefore, the contribution of individual bacterial factors to the mutualism, as well as to the predator-prey relation with the participation of one or multiple players can be easily delineated. These aspects render the system as a model promising to address questions concerning assignments of ecological functions for microbial natural products.

Over the past decade, it has been shown that some *XP* natural products involved in bacterial cell-cell communication, nematode development, insect pathogenicity, insect immune suppression, cytotoxicity, and antimicrobial activity, are instrumental in maintaining the complex life cycle^8^. Our previous metabolic analysis on 30 *XP* strains preliminarily revealed their biosynthetic capacity of natural products, via linking the metabolic profile of wild-type strains to known natural product biosynthetic gene clusters (BGCs)^16^. Intense research efforts have also been devoted to the characterization of unknown BGCs for natural product discovery, as well as functional assignments of natural products in the context of bacteria-nematode-insect interactions^8,17–22^. However, these studies either mostly revolved around the identification and characterization of individual BGCs on a single-genome basis, or lacked a comprehensive comparison of intra/interspecies BGCs. This did not reveal to what extent BGCs that might be linked to the special ecological niche are either conserved or unique within *XP* genomes. Therefore, a more systematic approach is needed to create a global BGC map that could be utilized for identifying BGCs of ecological importance across *Xenorhabdus* and/or *Photorhabdus*, as well as for exploring the full biosynthetic capacity of *XP* strains for accelerating genome mining.

Here, to provide insights into natural products that may account for the niche specificity of *XP*, we apply genome analysis of 45 *XP* strains that cover almost all *XP* taxonomy via combining pangenomic and domain sequence similarity network approaches, homologous expression of desired BGCs, chemical structure elucidation, and biological assays.

## RESULTS

### An overview of *XP* BGCs

We began by using the antibiotics & Secondary Metabolite Analysis Shell (antiSMASH) 5.0 (ref ^23^) to predict and annotate the natural product BGCs in 29 *Xenorhabdus* and 16 *Photorhabdus* strains (**Supplementary Table 1**). A total of 1,000 BGCs were detected and categorized into eight classes (**Fig. 1a**, **Supplementary Fig. 1**, and **Supplementary Table 2**), corresponding to an average of 22 BGCs per species which is 2-to 10-fold higher than the average BGC levels of any other Enterobacteria^24^. Most species show a linear relationship between the number of BGCs and the size of their genome (**Fig. 1b**). Compared to *Xenorhabdus*, *Photorhabdus* tends to harbor a larger genome size with more BGCs. Non-ribosomal peptide synthetases (NRPSs) are the most abundant BGC class in *XP*, accounting for 59% of the total BGCs, with approximately 13 BGCs per species. Due to the abundance of NRPS BGCs, it seems likely that their products play essential ecological roles. The “Others” group of BGCs comprised of various unclassified and unannotated BGCs is the second-largest class, whose products might facilitate bacteria to fulfill specific ecological functions. The polyketide synthase (PKS)/NRPS hybrid class is modestly enriched and broadly distributed. PKS (type I and other PKSs), ribosomally synthesized and post-translationally modified peptide (RiPP), terpene, and saccharide BGCs are significantly depleted in *XP*. Two earlier extensive BGC analyses of prokaryotes^25^ and human microbiota^26^ reveal that these sources are rich in saccharide BGCs and scant NRPS BGCs. Interestingly, these are contrary to the BGC composition of the nematode-symbiotic *XP*, suggesting *XP* BGCs as a distinct source for experimental natural product explorations.

**Fig. 1.**
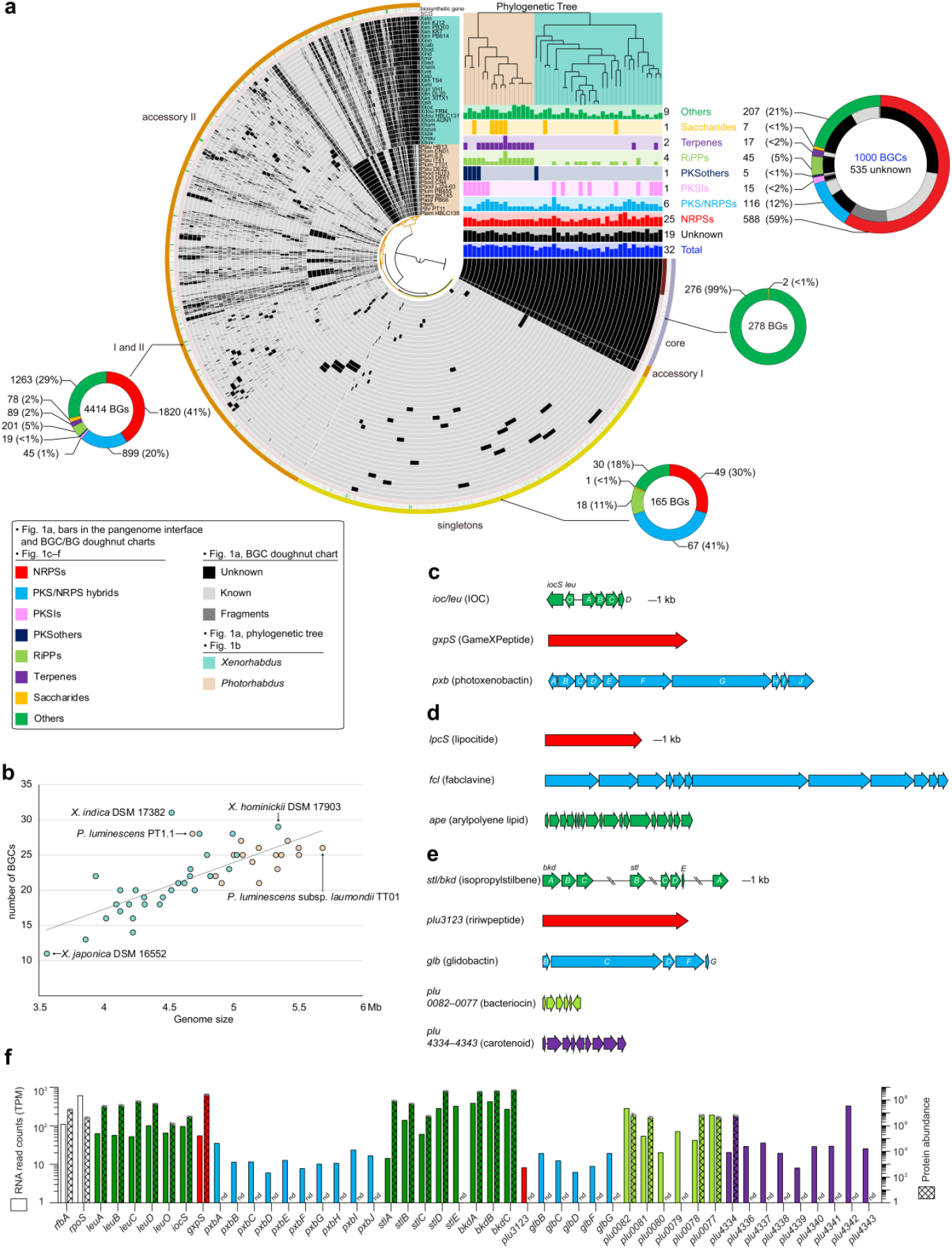
Pangenomic analysis and BGC overview of 45 *XP* genomes. **a,** Overview of classifications and distribution of BGs/BGCs in *XP*. In the circle interface, each layer (grey) represents all genes (black) in a single genome; single-copy-core-gene (scg, dark red); biosynthetic gene (green); bin names (core, grey-purple; accessories I and II, orange; singletons, yellow). The number and classification of BGCs in a strain is represented by the bar charts under the phylogenetic tree. The maximum number of each BGC class is indicated on the right side of the bar charts. The total number of BGC in each class and its percentage is indicated on the left side of the doughnut charts. The numbers of identified BGCs in total, unknown BGCs, and BGs in different pangenomic regions are indicated in the doughnut charts. **b,** Relationship between predicted BGCs and genome size for each strain. Strains that harbor the most BGCs or the largest genome size within the genus are denoted. **c,** The most widely distributed BGCs in *XP*, including the previously unidentified *ioc/leu* and *pxb*. **d,** The most widely distributed *X*-specific BGCs, including the previously unidentified *lpc*. **e,** The most widely distributed *P*-specific BGCs, including the previously unidentified *plu00822–0077*. **f,** Comparison of the transcriptional and translational levels of genes in the conserved BGCs (*ioc/leu*, *gxp*, *pxb*, *stl/bkd*, *plu3123*, *glb*, *plu00822–0077*, and *plu4334–4343*) in *P. luminescens* subsp. *laumondii* TT01 with the housekeeping genes (*rfbA* and *rpoS*). See Supplementary Tables 6 and 7 for actual values. Proteomic data represent mean ± s.d. from four independent experiments.

### Conserved *XP* BGCs

In the context of prokaryotic genome evolution driven by gene gain and loss over long periods of time, the gene content of a pangenome that comprises phylogenetically-related bacterial species reflects a record of responses to forces of natural selection that bacterial species confront^27^. Core genes shared by all species in a pangenome are essential for basic biological aspects, while accessory and singleton genes presented in some and one species, respectively, are regarded as “dispensable”. These “dispensable” genes are still allied to complementary biochemical pathways and functions that might endow unique advantages for ecological adaptation of bacteria^28^. Therefore, we asked a question here, among all predicted BGCs, if there exist highly conserved BGC(s) across *XP* genomes. To delve deeply into the BGCs, we monitored biosynthetic genes (BGs) by the Anvi’o^29^ pangenomic approach to characterize their distribution in core, accessory, and singleton regions^27^ (**Fig. 1a** and **Supplementary Table 3**). Surprisingly, although NRPS BGCs are prolific in *XP*, all of their BGs scatter in the accessory and singleton regions. The core region mostly encompasses BGs belonging to an unknown *ioc/leu* BGC, which is proposed to be related to β-lactone biosynthesis (**Fig. 1c**). The *gxpS* (**Fig. 1c**) responsible for GameXPeptide biosynthesis^30^ located in the accessory region is the most broadly distributed NRPS gene cluster family (GCF) across *Xenorhabdus* (72%) and *Photorhabdus* (93%), followed by the antiprotozoal rhabdopeptide/xenortide-like peptides^31^ (**Supplementary Fig. 2**) that are found in 51% of *Xenorhabdus* and 87% of *Photorhabdus*. A set of five consecutive BGs (*pxbF–J*) in the accessory region composes an unknown cluster (*pxb*; **Fig. 1c**) representing the most prevalent PKS/NRPS hybrid GCF across *Xenorhabdus* (58%) and *Photorhabdus* (81%).

To scrutinize the prevalence of genus-specific BGCs, we analyzed the pangenome of *Xenorhabdus* and *Photorhabdus* separately (**Supplementary Fig. 3** and **Supplementary Tables 4** and **5**). NRPS BGs related to the xenoamicin (*xab*) BGC^32^ and eight unknown BGCs are located in the core region of the *Xenorhabdus* pangenome. Among them, an unknown NRPS (*lpcS*; **Fig. 1d**) stands out since it exists in 96% (28 out of 29) of strains as the most widespread *Xenorhabdus-specific* (*X*-specific) GCF. In the accessory region of the *Xenorhabdus* pangenome, multiple consecutive BGs making up the broad-spectrum antimicrobial fabclavine^33^ BGC (*fcl*; **Fig. 1d** and **Supplementary Fig. 2**), are found in 44% of *Xenorhabdus* strains as the most prevalent genus-specific PKS/NRPS hybrid GCF. Sequential consecutive BGs that compose the *ape* BGC (**Fig. 1d**) are found exclusively in 76% of *Xenorhabdus* strains. The *ape* BGC synthesizing arylpolyene lipids^34^ (**Supplementary Fig. 2**) that protect the bacteria from oxidative stress and promote biofilm formation^35^ is the most prominent GCF among Gram-negative bacteria^25,34^. Isopropylstilbene (**Fig. 1e** and **Supplementary Fig. 2**) is a multipotent compound and an essential growth factor of dauer-stage nematodes^36^, whose BGs (*stl/bkd*) located in the core region are highly conserved across all *Photorhabdus* strains. In the accessory region of the *Photorhabdus* pangenome, BGs of glidobactin (*glb*) that is a potent eukaryotic proteasome inhibitor^37^, ririwpeptide^38^ (*plu3123*; **Supplementary Fig. 2**), and carotenoid (*plu4334–4343*), as well as an unknown bacteriocin (*plu0082–0077*) make up BGCs that represent the most widespread *Photorhabdus*-specific (*P*-specific) PKS/NRPS hybrid (93%, 15 out of 16), NRPS (87%), terpene (81%), and RiPP (93%) GCFs, respectively (**Fig. 1e**).

Although these BGCs (**Fig. 1c–e**) are widespread in *XP*, some of the chemical structures accounting for the biosynthetic pathways remain cryptic. Two major reasons for this might be the BGC being silent in wild-type strains under laboratory conditions, and/or product(s) being undetectable or difficult to isolate. We therefore leveraged our previous transcriptomic (**Supplementary Table 6**) and proteomic (**Supplementary Table 7**) datasets of *Photorhabdus luminescens* subsp. *laumondii* TT01 wild-type strain cultivated in LB medium^39^ to obtain information about the transcription and translation of the conserved BGCs. The transcriptomic data showed that in the exponential growth phase, all conserved BGCs are actively transcribed with different levels (**Fig. 1f**). However, proteins encoded by the *pxb*, *plu3123*, *glb*, *plu0082–0077*, and *plu4334–4343* BGCs are partly or completely untranslated, while almost all genes belonging to the putative β-lactone (*ioc/leu*), GameXPeptide (*gxp*), and isopropylstilbene (*stl/bkd*) BGCs are expressed with high protein abundance, comparable to the levels of housekeeping genes (**Fig. 1f**). The proteomic data, except for the case of the putative β-lactone BGC (*ioc/leu*), are in line with the previous metabolic analysis^16^, in which GameXPeptides and isopropylstilbene are the chemotypes in *Photorhabdus* wild-type strains. These findings hint that among the conserved BGCs yielding previously unidentified natural products, the *pxb*, *plu0082–0077*, and *plu4334–4343* are silent clusters due to translational regulation mechanisms, while the product(s) of β-lactone BGC (*ioc/leu*) should be present in the wild-type strain but has yet to be detected and characterized by means of standard spectroscopic methods.

### Unique *XP* BGCs

With the unidentified, conserved BGCs in hand, we set out to assess their biosynthetic novelty as well as the thorough biosynthetic capacity of *XP*. We subsequently compared *XP* BGCs against the MIBiG^40^ reference BGCs by the biosynthetic gene similarity clustering and prospecting engine (BiG-SCAPE) based on distance metrics^41^. The BiG-SCAPE analysis suggested biosynthetic uniqueness of 535 BGCs (53%) that found to be unrelated to the MIBiG BGCs and our in-house BGC data (**Supplementary Table 8**). 46% of NRPS, 61% of PKS/NRPS hybrid, 73% of PKSI, 97% of RiPP, 100% of saccharide, and 58% of other BGCs have yet to be identified. The previously unidentified *X*-specific *lpc* BGC, as well as most of the known *XP* BGCs (312 entries, 87%), including the aforementioned prevalent NRPSs (encoding the biosyntheses of GameXPeptide^30^, rhabdopeptide/xenortide-like peptides^31^, and ririwpeptide^38^) and PKS/NRPS hybrids (encoding the biosyntheses of fabclavine^33^ and glidobactin^37^), are concentrated in the main network (**Fig. 2**). This indicates they are very similar in terms of domain sequences. Most of the unknown BGCs (378 entries, 70%) distantly related to the known BGCs are mostly on the periphery of the main network (**Fig. 2**), exemplified by the previously unidentified PKS/NRPS hybrid BGC (*pxb*) that is prevalent across *XP*. The remaining 157 (30%) unknown BGCs, including the *XP* highly conserved β-lactone (*ioc/leu*) and *P*-specific bacteriocin (*plu0082–0077*) BGCs, are classified into 55 GCFs (as 26 isolated clades and 29 singletons) without connections with MIBiG references or the main network, suggesting their underlying biosynthetic novelty. Thus, the network might assist us in mining new natural products based on the distance of their encoded BGCs relative to that of MIBiG and known nodes.

**Fig. 2.**
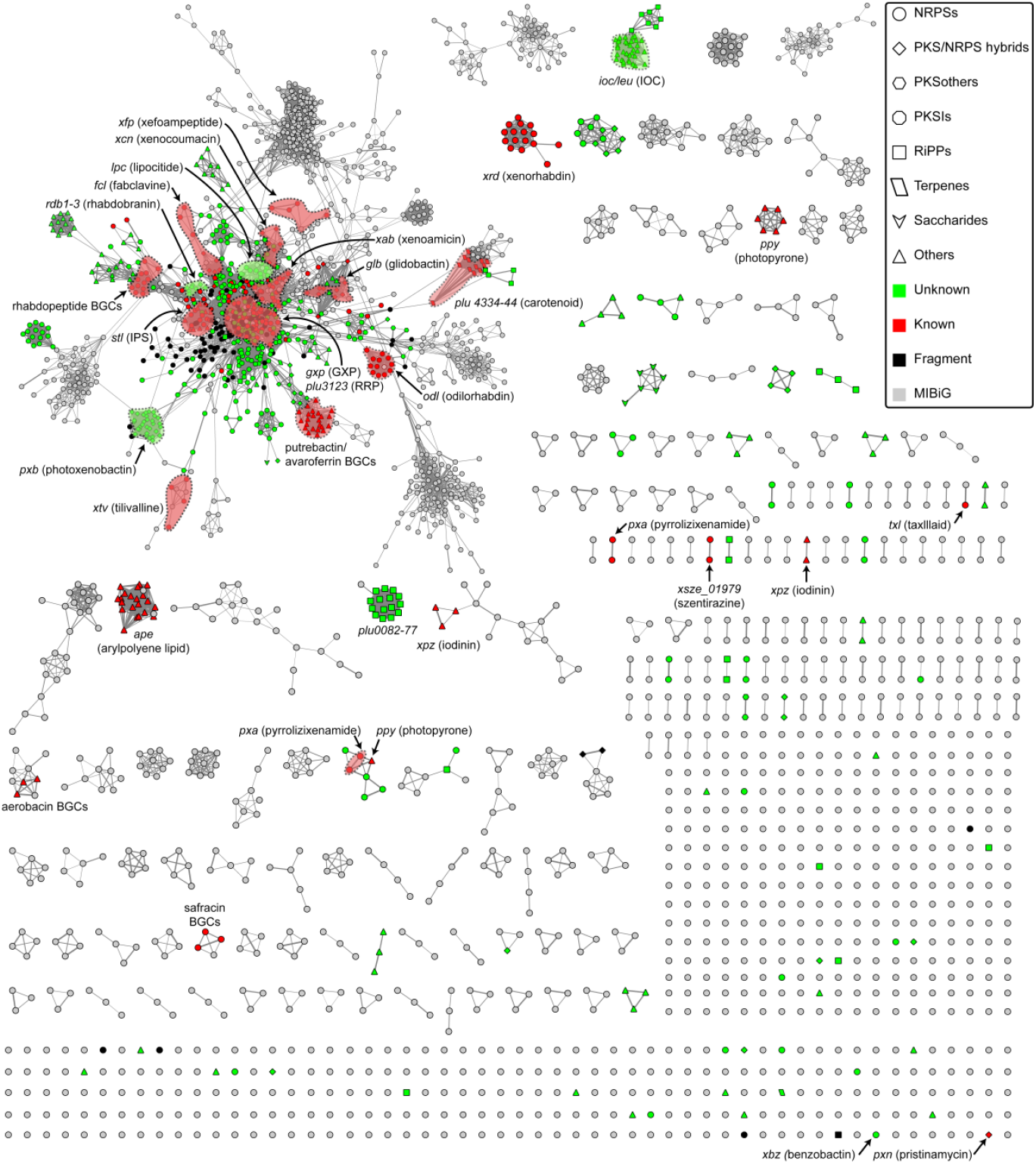
Sequence similarity network of BGCs identified in 45 *XP* genomes. Previously unidentified BGCs involved in this study and selected known BGCs are annotated and highlighted. BGCs in the main network belonging to a given GFC are not exhaustively highlighted due to nodes scattered. IPS, isopropylstilbene; GXP, GameXPeptide; RRP, ririwpeptide.

### 3-Isopropyl-4-oxo-2-oxetanecarboxylic acid (IOC) existing in all *XP* strains is a eukaryotic proteasome inhibitor

Recognizing that the putative β-lactone producing BGC is highly expressed under normal laboratory conditions (**Fig. 1f**) prompts us to predict a possible chemical structure based on the functions of biosynthetic genes, which might facilitate identification of the authentic product by re-examining the metabolic profile of wild-type strains. The putative biosynthetic cluster features six genes (**Fig. 3a** and **Supplementary Table 8**). *leuABCD* are involved in the l-leucine biosynthesis. *leuO* is positioned next to *leuA* and encodes a global transcription factor involved in regulating natural product biosynthesis^42^ and other physiological traits^43^. *iocS* encodes an enzyme belonging to the ANL (acyl-CoA synthetases, NRPS adenylation domains, and luciferase enzymes) superfamily. Such a gene architecture is reminiscent of the biosynthesis of cystargolides^44^, during which 3-isopropylmalate as an intermediate in the leucine pathway is the precursor for one-step lactonization to afford 3-isopropyl-4-oxo-2-oxetanecarboxylic acid (IOC, **1**) with a β-lactone moiety (**Supplementary Fig. 4**). Although the enzyme responsible for β-lactonization remains uncharacterized in the cystargolide biosynthesis^44^, a recent report demonstrates the acyl-AMP ligase, OleC, to be a β-lactone synthetase during the biosynthesis of long-chain olefinic hydrocarbons^45^. Therefore, we speculated that IocS might be responsible for adenylating the 4-carboxyl group and then triggering lactonization to give IOC (**1**).

**Fig. 3.**
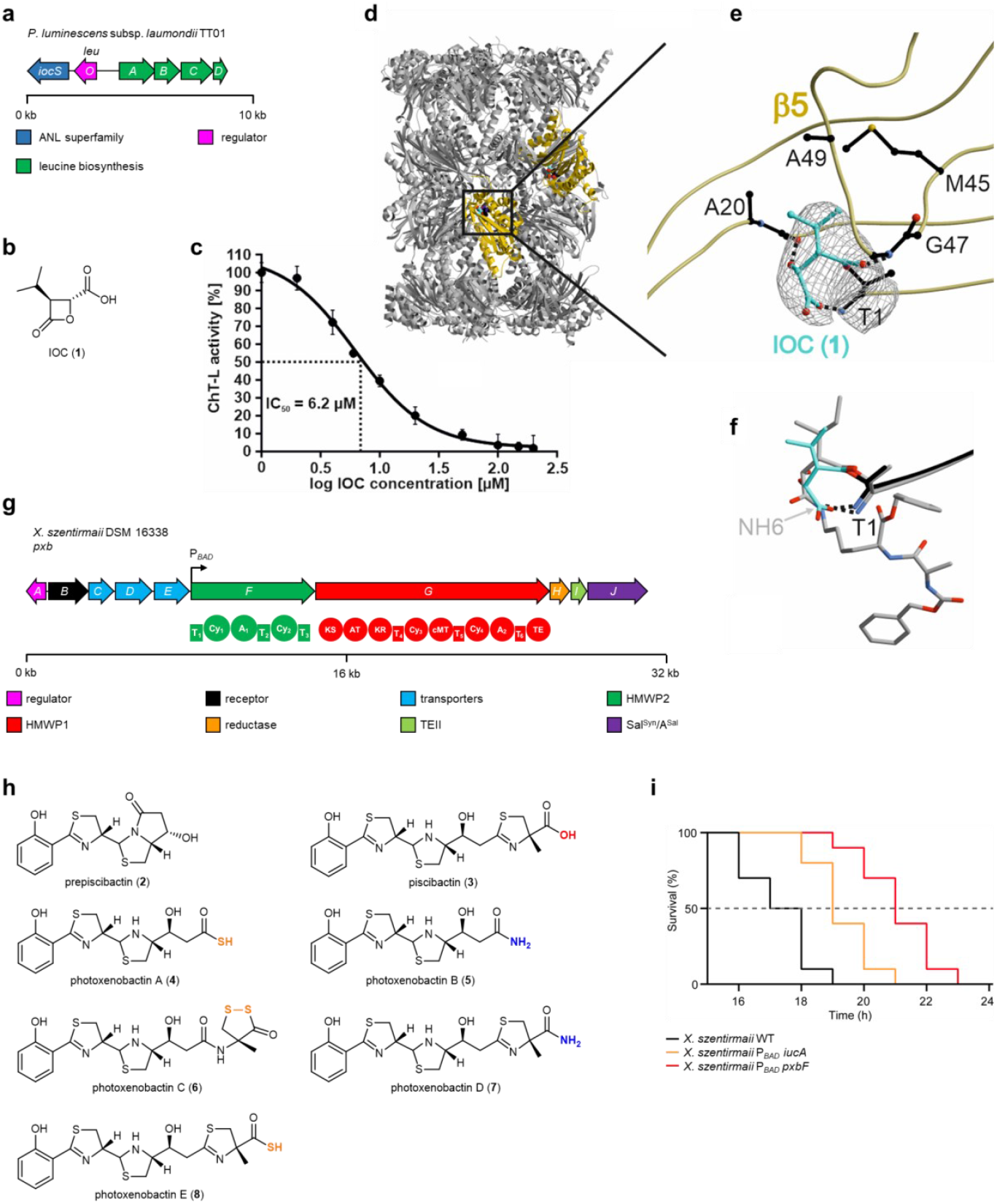
BGCs, chemical structures, and bioactivities of IOC and piscibactins/photoxenobactins. **a,** Genetic architecture of the *ioc/leu* BGC. **b,** Chemical structure of IOC (**1**). **c,** Dose–response curve for IOC-mediated inhibition of the β5 subunit of the yeast 20S proteasome. The IC_50_ value of **1** was determined to 6.2 ± 1.2 μM. Data represent mean ± s.d. from three independent experiments. **d,** Crystal structure of the yeast 20S proteasome in complex with **1** (spherical model, cyan carbon atoms) bound to chymotrypsin-like active sites (β5 subunits, gold, PDB ID 7O2L). **e,** Illustration of the 2*F*_O_–*F*_C_ electron density map (grey mesh, contoured to 1σ) of **1** covalently linked through an ester bond (magenta) to Thr1O^γ^ of subunit β5. Protein residues interacting with **1** are highlighted in black. Dots illustrate hydrogen bonds between **1** and protein residues. **f,** Superposition of **1** (cyan) and homobelactosin C (grey, PDB ID 3E47)^50^ complex structures with the yeast 20S proteasome highlights similar conformations at the chymotrypsin-like active site. **g,** Genetic architecture of the *pxb* BGC and domain organization. A black arrow shows the position where an l-arabinose-inducible promoter P_*BAD*_ is inserted. T, thiolation; A, adenylation; Cy, heterocyclization; KS, ketosynthase; AT, acyltransferase; KR, ketoreductase; cMT, carbon methyltransferase; TE, thioesterase. **h,** Known chemical structures of prepiscibactin (**2**) and piscibactin (**3**) from *Photobacterium damselae* subsp. *piscida^55^*, as well as previously unidentified photoxenobactins A–E (**4–8**) from *X. szentirmaii* DSM 16338. The terminal heteroatoms are highlighted. **i**, Survival curve of *G. mellonella* larvae (ten insects per strain) infected with *X. szentirmaii* WT (ø 79 cells), *X. szentirmaii* P*_BAD_ iucA* mutant (ø 81 cells), and *X. szentirmaii* P*_BAD_ pxbF* mutant (ø 90 cells). LT_50_ values: WT, 16.9 h; P*_BAD_ iucA*, 18.6 h; P*_BAD_ pxbF*, 20.3 h. The LT_50_ time point is indicated with a dot line. *Δ*LT_50_ = LT_50_^mutant^ – LT_50_^WT^. The *iuc* BGC encodes the biosynthesis of aerobactin in *X. szentirmaii*^60^. Under a non-induced condition during insect injection assays, *X. szentirmaii* P*_BAD_ iucA* and *X. szentirmaii* P*_BAD_ pxbF* mutants are equivalent to corresponding BGC knock-out mutants.

To detect the putative β-lactone, we cultured *P. luminescens* subsp. *laumondii* TT01, *Xenorhabdus nematophila* ATCC 19061, and *Xenorhabdus szentirmaii* DSM 16338 wild-type strains in various media. By HPLC–HRMS analysis of the culture supernatant from Sf-900 (a serum-free insect cell medium) with a negative ion mode, we did detect a peak with an *m/z* 157.0508 [M – H]^−^ whose deduced sum formula, C_7_H_9_O_4_, coincides with that of **1** (**Supplementary Table 10**). Finally, (2*R*,3*S*)-**1** was synthesized and showed identical retention time and MS/MS fragmentation patterns with **1** in HPLC–HRMS (**Supplementary Figure 5**), which confirmed the planar structure and stereochemistry of **1** (**Fig. 3b**).

The ubiquitin-proteasome system responsible for degrading misfolded and malfunctioning proteins in eukaryotes plays an essential role in cell-cycle regulation and apoptosis^46^. The system is also involved in degrading repressors of the insect immune response cascade^47^. The proteasome 20S core particle, the catalytic core of the system, is assembled from four stacked heptameric rings adopting an α_1-7_β_1-7_β_1-7_α_1-7_ stoichiometry^48^. The active site nucleophile of each proteolytic center is an N-terminal threonine (Thr1) located at subunits β1 (caspase-like activity), β2 (trypsin-like activity), and β5 (chymotrypsin-like activity)^49^. Natural products featuring a β-lactone moiety, such as omuralide, belactosins, and cystargolides, have been proven to suppress the proteolytic activity of the core particle^45^,^50^. Their uniform mode of proteasome inhibition relies on opening of the β-lactone and transesterification upon nucleophilic attack by the catalytic N-terminal threonine (Thr1O^γ^)^51^. Nevertheless, β-lactone natural products significantly differ in their chemical structures and thus, in their mode of binding. Inspired by cystargolides and belactosins containing an IOC moiety as reactive head group^50^,^52^, we assumed that IOC (**1**) might represent the smallest β-lactone that still blocks the activity of the proteasome. Indeed, **1** inhibits the yeast 20S proteasome with an IC_50_ value of 6.2 μM for the β5 subunit (**Fig. 3c**), whereas it has low binding affinities for β1 (625 μM) and β2 (60 μM). Therefore, we solved the crystal structure of **1** in complex with the yeast 20S proteasome at 3.0 Å (PDB ID 7O2L). The electron density map displayed **1** covalently bound to Thr1O^γ^ of all active sites due to the high ligand concentrations used for crystal soaking (**Fig. 3d**). However, since **1** lacks strong interactions with protein residues in the caspase- and trypsin-like binding channels, the 2*F*_O_-*F*_C_-map for the ligand is diffuse at β1 and β2. In contrast, **1** is well defined in the β5 subunit (**Fig. 3e**) and adopts a similar conformation as observed for homobelactosin C^50^ (**Fig. 3f**). Hereby, the acyl-oxygen atom of **1** derived from β-lactone ring-opening is stabilized by the oxyanion hole (Gly47NH), whereas the generated hydroxyl group is hydrogen-bonded to the carbonyl oxygen of residue 19. In similarity to NH6 in belactosin products^51^, the carboxylate group of **1** interacts with the threonine N-terminus and displaces the nucleophilic water molecule (**Fig. 3e**), thereby preventing hydrolysis of the acyl enzyme complex and explaining its inhibitory effect. Furthermore, the isopropyl moiety of **1** at the P1 site is stabilized by Ala20, Met45, and Ala49 in the chymotrypsin-like channel. Although these interactions are present in other β-lactone containing compounds, they adopt a diverse and unpredictable mode of binding. Without nitrogen atoms and extension units, **1** might feature the minimal scaffold for proteasome inhibition. Therefore, **1** could be an *XP* universal virulence factor against insects, as well as soil-living food competitors like protozoa, through disturbing the ubiquitin-proteasome system and thereby causes cell-cycle disturbance and immunodeficiency.

### Photoxenobactins as the most prevalent polyketide/non-ribosomal peptide hybrids across *XP* featuring unusual C-termini are virulence factors against insects

The prevalent PKS/NRPS hybrid GCF containing 32 *pxb* (<u>p</u>hoto<u>x</u>eno<u>b</u>actin) BGCs is shown to have weak similarity to micacocidin^53^ and yersiniabactin^54^ BGCs in the BiG-SCAPE network (**Fig. 2** and **Supplementary Table 8**). Notably, compared with HMWP1 encoded by the yersiniabactin BGC in *Yersinia pestis*^54^, its homolog (PxbG) lacks one carbon-methyltransferase domain (cMT_1_) involved in the bismethylation of a C2 polyketide moiety in yersiniabactin. Moreover, PxbG embeds an additional module comprising a heterocyclization domain, an A domain, and a T domain (Cy_4_–A_2_–T_6_; **Fig. 3g** and **Supplementary Fig. 6**).

To unveil the underlying biosynthetic theme of *pxb* BGC, we overexpressed the cluster in *X. szentirmaii* DSM16338 by promoter exchange strategy^19^ to insert a P_*BAD*_ promoter in front of *pxbF*. Besides prepiscibactin (**2**) and piscibactin (**3**)^55^, the *X. szentirmaii* P*_BAD_ pxbF* mutant yielded four additional compounds, all of which were also present in the wild-type strain but with low production titers in Sf-900 medium (**Supplementary Fig. 7**). These four compounds share similar MS/MS fragmentation patterns to **3** and yersiniabactin in the low mass region including a diagnostic *m/z* 190.031 [M + H]^+^ fragment ion of hydroxyphenylthiazoline^56^ (**Supplementary Fig. 8**) which indicates these compounds are new piscibactin derivatives, termed photoxenobactins A–D (**4**–**7**; **Fig. 3h**). From a 40-L fermentation broth of the *X. szentirmaii* P*_BAD_ pxbF* Δ*hfq* mutant that produced desired compounds with a reduced background of other natural products for facile purification^19^, we obtained **4**–**6**, as well as photoxenobactin E (**8**; **Supplementary Fig. 7**). The chemical structures of **4**, **5**, and **8** were readily elucidated by HRMS and NMR spectroscopic methods and that of **7** was confirmed by tandem MS (**Supplementary Figs. 8** and **9**), revealing that, unexpectedly, **4**, **5**, **7**, and **8** have various chain lengths and termini such as thiocarboxylic acid (**4** and **8**) and carboxamide (**5** and **7**). Although the production titer of photoxenobactin C (**6**) in the *X. szentirmaii* P*_BAD_ pxbF* mutant appeared to be sufficient for isolation (**Supplementary Fig. 7**), we only obtained a trace amount of the pure compound. Photophobia and thermo-instability in any kind of organic solvents have been found to be the culprit, leading to conversion into an array of rearranged products, such as methyl ester piscibactin (**9**) in methanol (**Supplementary Fig. 10**). Finally, combining extensive labeling experiments (**Supplementary Figs. 11 and 12**), and 2D NMR data (**Supplementary Fig. 9**), we proved that **6** bears a unique dithioperoxoate moiety.

Inspired by piscibactin capable to chelate gallium and ferric ions^55^, we set out to explore whether photoxenobactins are metallophores since this compound class is essential for bacteria to acquire trace elements from environments with additional function (e.g. toxicity, signal, protection, antibiotics)^57^. A fraction mainly containing *pxb* BGC products were incubated with different inorganic metal salts (e.g. Ga^III^, Fe^III^, Cu^II^, Zn^II^, Mo^VI^, and V^V^), and only piscibactin-Ga^III^/-Fe^III^/Cu^II^ (**10**–**12**) and photoxenobactin D-Ga^III^/-Fe^III^/Cu^II^ (**13**–**15**) were detected (**Supplementary Fig. 13**). It has been shown recently that in addition to iron scavenging, yersiniabactin forms complex with Cu^II^, which serves as a superoxide dismutase mimic involved in the protection against the oxidative burst in phagocytes^58^. Due to the high structural similarity between **3**, **7**, and yersiniabactin, as well as the cupric chelating property of **3** and **7**, it is tempting to speculate that **3** and **7** scavenge environmental copper, forming cupric complexes to protect the bacteria from reactive oxygen species generated by the insect immune system. An earlier report describes killings of *Galleria mellonella* upon injection of *Escherichia coli* carrying a *pxb* BGC from *Photorhabdus asymbiotica*. Ulbactin E and a compound with a sum formula C_20_H_25_O_4_N_3_S_3_ which was predicted to be a desmethyl yersiniabactin were found in the methanol extract of insect carcasses, suggesting both compounds as virulence factors against insects^59^.

Indeed, C_20_H_25_O_4_N_3_S_3_ coincides with methyl ester piscibactin (**9**), a rearranged product of **6** that occurs in methanol as observed herein (**Supplementary Fig. 13**). Therefore, we reasoned that **6** should be one of the authentic insecticidal compounds. Next, we attempted to re-examine the toxicity of *pxb* BGC products during the insect infection process by comparing it with that of aerobactin, an identified virulence-related siderophore in *X. szentirmaii*^60^. We then injected *X. szentirmaii* wild-type strain, which was shown to produce aerobactin^60^ and **2**–**7** (**Supplementary Fig. 7**), as well as the *pxb* (*X. szentirmaii* P*_BAD_ pxbF*) and *iuc* (*X. szentirmaii* P*_BAD_ iucA*; aerobactin) knock-out mutants into *G. mellonella* larvae. The *pxb* BGC knock-out mutant killed insects 3.4 h [*Δ*median lethal time (LT_50_)] more slowly than the wild-type strain (**Fig. 3i**). Furthermore, the *pxb* BGC products exerted a greater impact on insect virulence than aerobactin, whose knock-out mutant showed *Δ*LT_50_ = 1.7 h.

### GameXPeptides as the most widespread non-ribosomal peptides across *XP* inhibit insect immune responses

GxpS, an NRPS with five modules (**Fig. 4a**), is responsible for the biosynthesis of GameXPeptides, which are a class of cyclic pentapeptides composed of valine, leucine, and phenylalanine (**Fig. 4b**). Although GameXPeptides are one of the diagnostic chemotypes with high production titers in almost all *XP*^16^, their function has remained cryptic over the past decade. Our recent bioactivity screening for crude extracts produced by specifically overexpressed mutants^19^ indicated that GameXPeptides might be one of the bioactive contributors of wild-type strains inhibiting in vitro production of prostaglandin E2 without cytotoxicity and antimicrobial activity. With the synthetic GameXPeptide A (**16**) in hand, we therefore pursued its possible suppression of insect immune responses. Insects rely on innate immunity consisting of cellular and humoral immune responses to overcome infections^61^. Cellular immune responses mediated by eicosanoids involve encapsulation that is performed by immune hemocytes along with morphological changes, melanization activated by phenoloxidase, nodulation, and phagocytosis^62^. The cytoplasmic extension observed in the hemocytes of lepidopteran insect, *Spodoptera exigua*, as an immune response to the *E. coli* challenge was significantly inhibited by **16** (**Fig. 4c**) in a dose-dependent manner (**Fig. 4d**) with an IC_50_ value of 17.2 ng/larva (**Supplementary Table. 11**). Although **16** exerted no suppression against the phenoloxidase activation (**Fig. 4e**), it remarkably decreased the number of nodules formed (**Fig. 4f**) in a dose-dependent manner (**Fig. 4g**) with an IC_50_ value of 25.8 ng/larva (**Supplementary Table 12**). These results suggest that **16** specifically suppresses insect hemocyte-spreading and nodule formation upon insects being challenged by *E. coli*, and thereby defeats the insect cellular immune response. It is worth mentioning that the inhibitions of phenoloxidase activity and the proteolytic cascade leading to active phenoloxidase are accomplished by two known widespread compound classes, rhabduscin^63^ and rhabdopeptide/xenortide-like peptides^8^, respectively. Consequently, the functional characterization of ubiquitous GameXPeptides is one of the most important jigsaws for deconstructing *XP* to suppress insect immune systems during symbiotic nematode invasion.

**Fig. 4.**
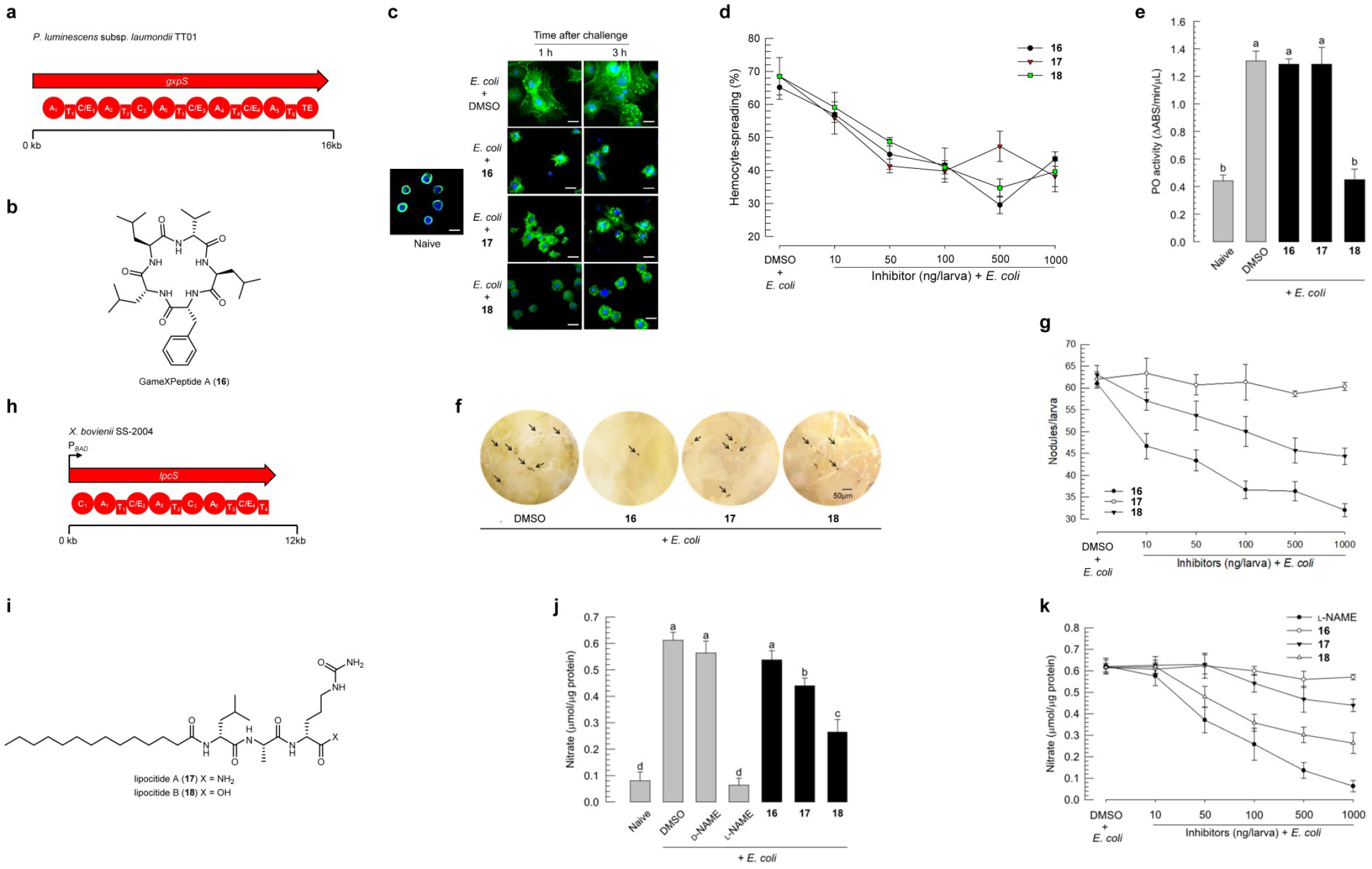
BGCs, chemical structures, and bioactivities of GameXPeptide A and lipocitides. **a**, Domain organization of GxpS. **b**, Chemical structures of GameXPeptide A (**16**). **c**, In vivo observation of hemocytes-spreading behavior in different time intervals upon injection of **16**–**18** (1000 ng/larva) in *S. exigua* larvae. Blue, nucleus; green, actin cytoskeleton. **d**, In vitro analysis of hemocyte-spreading behavior. **16**–**18** suppressed hemocyte-spreading in a dose-dependent manner with IC_50_ values of 17.2, 10.0, and 26.2 ng/larva, respectively. **e,** Suppression of PO activity in *S. exigua* larvae by **18** (1000 ng/larva). PO, phenoloxidase. **f**, Suppression of hemocyte nodule formation in *S. exigua* larvae by **16** and **18** (1000 ng/larva). Nodules were counted at eight-hour post-infection (black spots indicated with black arrows). **g**, Dose-dependent suppression of nodule formation by **16** and **18** with IC_50_ values of 25.8 and 86.1 ng/larva, respectively. **h**, Domain organization of the LpcS. A black arrow shows the position where an l-arabinose-inducible promoter P_*BAD*_ is inserted. **i,** Previously unidentified chemical structures of lipocitides A (**17**) and B (**18**) from *X. bovienii* SS-2004. **j**, Suppression of NO production in the hemolymph of *S. exigua* larvae injected with **17** and **18** (1000 ng/larva). **k**, Dose-dependent suppression of NO production in the hemolymph of *S. exigua* larvae by **17** and **18**. l-NAME (N_ω_-nitro-l-arginine methyl ester hydrochloride) and d-NAME (N_ω_-nitro-d-arginine methyl ester hydrochloride) as controls. kb, kilobase. C, condensation; A, adenylation; T, thiolation; E, epimerization. Each treatment consists of three independent replications on five larvae. Data represent mean ± s.d. Letters above standard error bars indicate significant differences among means at Type I error = 0.05 (LSD test).

### Lipocitides as the most widespread *Xenorhabdus-specific* natural products inhibit insect immune responses via the NO signal transduction pathway

The most broadly distributed *X*-specific GCF existing in all but *Xenorhabdus cabanillasii* JM26 is centralized in the main BiG-SCAPE network and displays a degree of relatedness with the *xcn* (xenocoumacin)^64^ and *fcl* (fabclavine)^33^ GCFs (**Fig. 2** and **Supplementary Fig. 8**). We designated this *X*-specific cluster as *lpc* which encodes a tetramodular NRPS with an unusual terminal T–C/E–T domain architecture (**Fig. 4h**). This BGC is silent under laboratory conditions, consistent with the transcriptional level of *lpcS* in *X. szentirmaii* US wild-type strain^39^ being about 16-fold lower than those of the housekeeping genes (**Supplementary Table 13** and **Supplementary Fig. 14**). We were able to activate the BGC in *Xenorhabdus bovienii* SS-2004 by the promoter-exchange strategy. The *X. bovienii* P*_BAD_ lpcS* mutant produced an array of N-terminal acylated linear tripeptides (**Supplementary Fig. 15**). Two major products, lipocitides A and B (**17** and **18**; **Fig. 4i**), were purified and their structures were identified by NMR spectroscopy, Marfey’s method, and chemical synthesis (**Supplementary Fig. 9** and **Supplementary Note**), revealing that **17** and **18** bear a myristoyl and a consecutive amino acid sequence of d-leucine/l-alanine/d-citrulline, as well as a carboxamide and carboxylic acid in their respective C-termini. Comparison of the tandem MS of the other lipocitides in *X. bovienii* SS-2004 with **17** and **18** revealed that lipocitides feature either d-leucine/l-alanine/d-citrulline-OH or d-leucine/l-alanine/d-citrulline-NH_2_ as a backbone and differ in the N-acyl substitutions (**Supplementary Fig. 16**).

Nitric oxide (NO) converted from l-arginine by NO synthases is an upstream component of the eicosanoid signaling pathway to trigger insect innate immune responses against exogenous challenges^62^. Inspired by l-citrulline and arginine-derived compounds being inhibitors of NO synthesis^65^, we examined whether the major lipocitides, **17** and **18**, could inhibit NO production to defeat insect immune responses. The elevated NO concentration in the hemolymph of *S. exigua* larvae caused by *E. coli* infection was suppressed by both compounds (**Fig. 4j**) in a dose-dependent manner (**Fig. 4k**) with IC_50_ values of 2.37 and 0.42 μg/larva, respectively (**Supplementary Table 14**). Earlier reports showed that NO activates phospholipases A_2_ for producing downstream eicosanoid signaling molecules^66^, thereby mediating cellular immune responses. Both **17** and **18** suppressed the cytoplasmic extension in the hemocytes of *S. exigua* upon *E. coli* challenge (**Fig. 4c**, **4d** and **Supplementary Table 11**). In addition, **18** significantly suppressed the phenoloxidase activation (**Fig. 4e**) and decreased the number of nodules formed (**Fig. 4f**, **4g**, and **Supplementary Table 12**). These results indicate that lipocitides suppress insect NO production, which leads to sequential inhibitions on cellular immune responses, and thus might cause fatal immunosuppressive conditions of the insects under infection of *Xenorhabdus* symbiotic nematode. In contrast, GameXPeptide A (**16**) displayed no suppression on NO production (**Fig. 4j**, **4k**), which indicates GameXPeptides have a different upstream target from lipocitides or mediate other signaling transduction pathways.

### Rhabdobranin with a prodrug activation mechanism during biosynthesis is a T-shape polyketide/non-ribosomal peptide hybrid

The above survey of previously unidentified conserved BGCs has showcased the abilities of *XP* to produce pervasive and structurally unique natural products. We then set out to examine the uncharacterized BGCs that only exist in specific species to assess the biosynthetic potential of *XP*. In the BiG-SCAPE main network, eight unknown PKS/NRPS hybrid BGCs from seven *Xenorhabdus* and one *Photorhabdus* compose a GCF, termed *rdb* (<u>r</u>hab<u>d</u>o<u>b</u>ranin). The *rdb* BGCs feature a peptidase encoded gene, suggesting a prodrug activation mechanism similar to that observed during the biosyntheses of xenocoumacin/amicoumacin that are potent antibiotics inhibiting mRNA translation^8^ and colibactin that is a genotoxin alkylating DNA^11^. Although the *rdb* GCF is adjacent to rhabdopeptide/xenortide-like BGCs, it connects neither to amicoumacin and xenocoumacin^64^ BGCs nor to any MIBiG entries. We classified these eight highly similar BGCs into three types, *rdb1–3*, based on the presence/absence of the first A domain in RdbH and the TE domain in RdbI (**Fig. 5a**, **Supplementary Fig. 17**, and **Supplementary Table 9**), which might lead to products with distinct numbers of amino acid residues and non-linear biosynthetic assembly line logic, respectively. *rdb1* encodes a weakly predicted TE domain in Rdb1I, while that encoded in *rdb2* and *rdb3* is clearly annotated. Additionally, an extra type II TE, Rdb3V, is encoded in *rdb3*. Compared with Rdb1H and Rdb2H, the Rdb3H lacks an A domain.

**Fig. 5.**
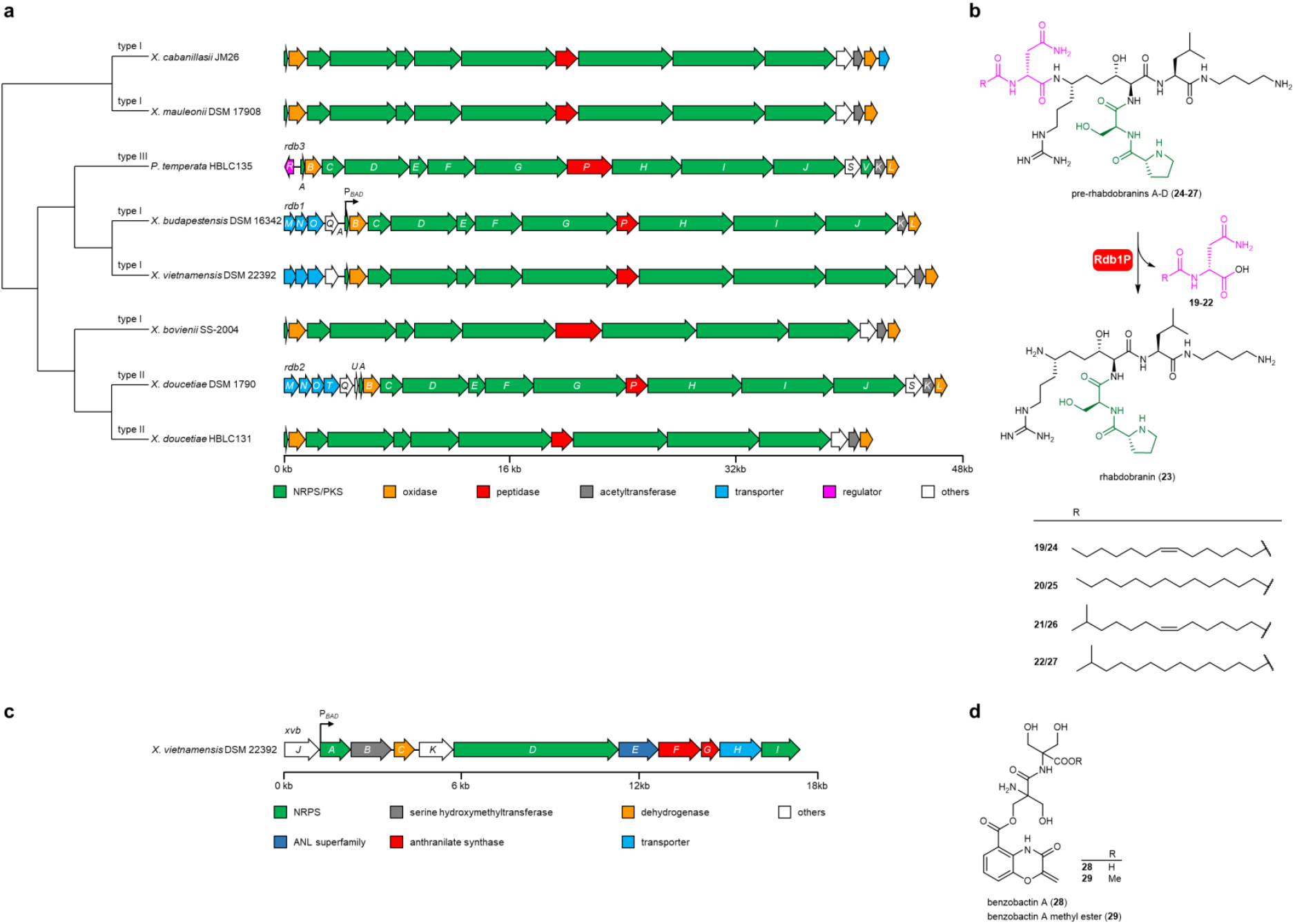
Representative BGCs of uniqueness and chemical structures thereof in *XP*. **a,** Phylogenetic tree and gene organization of the *rdb* BGCs. Phylogenetic tree is based on protein sequences of BGCs. BGC subclassification is indicated next to the branch. **b,** Chemical structures of previously unidentified pre-rhabdobranins A–D (**24**–**27**) and rhabdobranin (**23**) from *X. budapestensis* DSM 16342, as well as the proposed late-stage biosynthesis involved in a prodrug activation mechanism, similar to xenocoumacin and colibactin. The N-terminus capped acylated d-asparaginyl moiety (**19**–**22**) and the dipeptidyl branch is highlighted in pink and green, respectively. The stereocenters were predicted by analyzing the conserved motif in C domain and KR domain that are responsible for stereocontrol. **c,** Genetic architecture of the *xvb* BGC. **d**, Previously unidentified benzobactins A (**28**) and a methyl ester thereof (**29**) from *X. vietnamensis* DSM 22392. A black arrow shows the position where an l-arabinose-inducible promoter P_*BAD*_ is inserted. kb, kilobase.

To identify products derived from this GCF, we focused on *rdb1* that contains five out of eight BGCs in this GCF, and attempted to activate the *rdb1* in *Xenorhabdus budapestensis* DSM 16342 by inserting a P_*BAD*_ promoter in front of *rdb1A*. The *X. budapestensis* P*_BAD_ rdb1A* mutant yielded four N-myristoyl-D-asparagine congeners (**19**–**22**), as well as a non-XAD-resin-bound hydrophilic compound with a low production level (**23**; **Supplementary Fig. 18**). Since an acylated D-asparaginyl capping the N-terminus of xenocoumacin, zwittermicin, and colibactin has been found to be a self-resistance mechanism^67^, the detection of N-myristoyl-d-asparagine analogs was consistent with our hypothesis that a prodrug strategy was involved in the *rdb* biosynthesis. To accumulate the inactive prodrugs for structural identification, we deleted the peptidase encoded gene, *rdb1*P, and the resultant *X. budapestensis* P*_BAD_ rdb1A* Δ*rdb1P* mutant led to loss of **19**–**22** and high production of four new peaks with larger masses, designated as pre-rhabdobranins A–D (**24**–**27**; **Fig. 5b**) with differences in the N-acylated moiety. A 12-L culture of the *X. budapestensis* P*_BAD_ rdb1A* Δ*rdb1P* Δ*hfq* mutant was performed, allowing for the isolation of pre-rhabdobranin D (**27**), whose structure was determined by HRMS and NMR spectroscopy (**Supplementary Fig. 9** and **Supplementary Table 10**). Intriguingly, pre-rhabdobranins are characterized by a proline-serine dipeptidyl side chain that branches off at the N atom of an aminomalonyl building block. To the best of our knowledge, this represents the first T-shape peptide in contrast to the canonical linear-chain-elongation on thiotemplated assembly lines. Moreover, the second module of Rdb1I (A–T–TE; **Supplementary Fig. 17**) appears to be inactive, and presumably, the C domain of Rdb1I recruits a putrescin for peptidyl chain off-loading.

### Benzobactins with cytotoxicity are synthesized by an orphan thiotemplated-based assembly line by recruiting noncanonical building blocks

BGCs as singletons in the BiG-SCAPE network could be ideal test cases for genome mining for the purpose of novel natural product discovery. We selected an NRPS BGC termed *xvb* (*<u>X</u>. <u>v</u>ietnamensis* DSM 22392 <u>b</u>enzobactins) for characterization (**Fig. 5c**). A more detailed analysis of the BGC composition shows that the substrate specificity of A domains was unpredictable by antiSMASH and there exist genes encoding specialized tailoring enzymes for substrate modification (e.g. a putative serine hydroxymethyltransferase encoded by *xvbB*), as well as genes encoding synthases for non-amino acid substrates (two putative anthranilate synthases encoded by *xvbF* and *xvbG*). These indicate the *xbv* product(s) might contain noncanonical building blocks. To determine the product(s) derived from this orphan BGC, we inserted a P_*BAD*_ promoter to express the *xvb* BGC that yielded benzobactin A (**28**) and its methyl ester (**29**; **Supplementary Fig. 19**). Their structures were confirmed by HRMS and NMR spectroscopy methods (**Supplementary Fig. 9** and **Supplementary Table 10**), revealing that **28** and **29** feature a rare benzoxazolinate moiety that was only found in C-1027^68^ and ashimides^69^ produced by *Streptomyces*, as well as a non-proteinogenic amino acid residue 2-hydroxymethylserine that is an unprecedented building block in natural products (**Fig. 5d**). **28** showed cytotoxic activity against the HepG2 cell line with an IC_50_ value of 19.0 μg/mL.

## Discussion

On the journey to understand the roles of *XP* natural products on mediating bacteria-nematode-insect interactions in the ecological niche, we previously carried out a large-scale metabolic exploration of 30 *XP* strains by rapid MS-based network analysis, as well as a survey of the correlation between the occurrence of known natural products and the BGCs encoding their biosyntheses^16^. This reveals that under standard laboratory conditions, the wild-type strains produce a plethora of natural products, most of which belong to the compound class of non-ribosomal peptides. However, in general, the MS-based network approach is constrained by 1) BGCs that are transcriptionally and/or translationally silent under standard laboratory conditions (e.g. BGC expressions need to be in an insect mimicking medium^39,70^ or under iron-limited conditions^60^); 2) compounds that are membrane-bound (e.g. arylpolyene lipids^34^) and that are difficult to be detected by standard LC/MS methods (e.g. compounds that are extremely hydrophilic/hydrophobic, too small/large, or poorly ionized/fragmented).

Here, to overcome the limitations of the metabolic analysis, we take the “BGCs first” strategy, since BGCs account for the genomic capacity of a strain for producing natural products. To date, the entomopathogenic bacteria *XP* have not been systematically analyzed in this regard. A promoter exchange strategy in the wild-type strain or Δ*hfq* mutant for homologously overexpressing BGCs of interest is then applied to efficiently translate BGCs into truly natural products for isolation, structural elucidation, and functional characterization. We examined the genomes of 45 *XP* strains covering almost all currently found *XP* taxonomy that are accessible in our strain collection, via pangenomic analysis with BGC annotations by antiSMASH. This allows us to visualize the distribution of BGs in the core, accessory, and singleton regions of the pangenomes. By tracking the frequencies of occurrence for consecutive BGs in the core and accessory regions, we were able to rapidly refine the most highly conserved BGCs among 1,000 BGC entries. We discovered that 11 BGCs, belonging to the classes of NRPSs, PKS/NRPS hybrids, RiPPs, terpenes, and others, represent the most ubiquitous GCFs across *XP* or in one of the two genera (**Fig. 1c–e**). Among them, five GCFs, namely *ioc/leu*, *pxb*, *lpc*, *plu0082–0077*, and *plu4334–4343* were previously unidentified.

All the *XP* species live under nearly the same ecological niche but harbor distinctive BGCs in terms of numbers and classes. For example, the number of BGCs in *Xenorhabdus indica* DSM 17382 is three times as high as that in *Xenorhabdus japonica* DSM 16552. *Photorhabdus temperata* subsp. *thracensis* DSM 15199 features seven BGC classes while *X. japonica* DSM 16552 only has three classes (**Supplementary Fig. 1**). Therefore, we assume that such deviations among *XP* species are possibly indicative of a minimum number of required BGCs—the highly conserved ones—for *XP* to maintain their lifestyle adaptation. The *ioc/leu* BGC responsible for IOC (**1**) biosynthesis was found to be present across all *XP* genomes, while there is not a specific NRPS GCF that universally exists in every *XP* species though the NRPSs are the most abundant class. Indeed, the *ioc/leu* BGC is also widely distributed in other γ-Proteobacteria like free-living pathogens *Vibrio cholerae* and *Y. pestis* (GCF_00781 in **Supplementary Table 15**). Although this BGC has yet to be studied in other microorganisms and the degree of structural conservation of IOC (**1**) among γ-Proteobacteria is unknown, it is conceivable that the conservation of structural genes, *leuA–D* for l-leucine biosynthesis and *iocS* for putative lactonization, can serve as an indicator that IOC (**1**) is highly conserved among γ-Proteobacteria targeting the eukaryotic proteasome. The *pxb* BGC as the most widespread PKS/NRPS hybrid GCF across *XP* produces piscibactins (**2** and **3**) and photoxenobactins (**4**–**8**), both of which are structurally related to yersiniabactin but with different chain-lengths and C-termini. In contrast to the precise target-oriented biosynthesis of the yersiniabactin BGC, it appears that the *pxb* BGC is more diversity-oriented though the biosynthetic machinery remains cryptic. Yersiniabactin with high affinities for ferric iron contributes to the virulence of human pathogens like *Y. pestis and E. coli*, which has been described a long history in the literature^71^. Our study showed that the *pxb* products are associated with the insecticidal activity of *X. szentirmaii*, but only piscibactin (**3**) and photoxenobactin D (**7**) retain metal-chelating abilities. This suggests that the other *pxb* products might be non-metal-chelation virulence factors against insects. In particular, photoxenobactin C (**6**) with a dithioperoxoate moiety is highly reactive and thus might account for the overall insecticidal activity of strains expressing the *pxb* BGC. As GameXPeptides and lipocitides are insect immune inhibitors targeting different transduction pathways as demonstrated here, both compound classes could synergistically contribute to a potent overall effect that producer strains can benefit from. The discovery of the widespread genus-specific GCFs might serve as a support to a point previously hypothesized regarding functional complementation achieved by structurally related natural products^16^. The pigmented arylpolyene lipids with two large conjugated systems protect its producing strain against reactive oxygen species^35^. While the *ape* BGCs encoding their biosynthesis are supposed to be the most ubiquitous GCF in Gram-negative bacteria^25,34^, the *ape* GCF are exclusively present in *Xenorhabdus* but absent in *Photorhabdus*. The *plu4334–4343* BGC as the most widely distributed *P*-specific terpene GCF is supposed to produce carotenoids that could conceivably fulfill the role of antioxidative protection^72^ for *Photorhabdus*. The chemical structure identification and functional characterization of the most ubiquitous *Xenorhabdus* and/or *Photorhabdus* natural products have made substantial progress towards deconstructing the niche specificity of *XP*.

Domain sequence similarity network was used to assess the full biosynthetic capacity of *XP*, which enables us to extensively map relationships between the *XP* BGCs and MIBiG references and thereby pinpoints BGCs that have great potentialities for producing novel natural products. This leads to the discovery of 535 unknown BGCs, 30% of which are thought to be unique. A recent survey of a correlation between taxonomic distance in myxobacteria and natural product novelty points out that the probability of novel natural product discovery could be increased by exploring genera located in taxonomically distant clades^73^. This strategy could be extensively applied in free-living microorganisms. However, it is less likely to fulfill in *XP* strains, because of their narrow ecological niche, in which bacteria are obligate mutualistic associations with specific soil nematodes, as well as limited taxonomic strains—only 26 *Xenorhabdus* and 19 *Photorhabdus* have been identified^74^. Nonetheless, inspired by the discovery of structurally unique photoxenobactins (**4**–**8**), pre-rhabdobranins (**24**–**27**), and benzobactins (**28** and **29**), genome mining for novel natural products from *XP* might yet be possible. A rarefaction analysis showed that the number of GCFs would increase upon the addition of every new genome, suggesting that sequencing of more *XP* isolates will probably reveal further previously unseen GCFs (**Supplementary Fig. 20**). This implies that the acquisition event of BGCs by horizontal gene transfer constantly occurs in *XP*, despite their relatively narrow niche. Distributions of GCFs at the genus level, as well as a matrix comparing GCFs to species, were then created to infer the differences within *XP* (**Supplementary Fig. 21**). While *Xenorhabdus* and *Photorhabdus* genus have 33 out of 176 common GCFs, a striking GCF diversity exists in *Xenorhabdus* even though *Xenorhabdus* has a tendency for fewer BGCs per genome than *Photorhabdus* (**Fig. 1b**). *X*-specific GCFs are 1.5 times as many as *P*-specific GCFs per species. Together with the higher number of unique *Xenorhabdus*-specific GCFs on average, which is in line with the more discrete GCF distribution pattern at the species level of *Xenorhabdus*, these could point towards *Xenorhabdus* being more likely to encode novel BGCs compared to *Photorhabdus*. Additionally, despite there being limited taxonomic species, BGC exploration within-species variation still holds great promise. For example, *Xenorhabdus stockiae* taxonomically related strains, *Xenorhabdus* sp. KK7.4, *Xenorhabdus* sp. KJ12.1, *Xenorhabdus* sp. PB30.3, and *Xenorhabdus* sp. PB61.4, over-representing across Thailand^75^ harbor seven unique genus-specific GCFs, including four isolated clade GCFs and three singletons (**Supplementary Fig. 22**).

*XP* adaptation to the harsh environment and competition against other soil microorganisms might be a driving force for the selection of valuable BGCs that produce highly efficacious natural products. Combining the pangenomic and sequence similarity network approaches provides deeper insights into BGCs responsible for natural product formation, and thereby allows more systematic inference of associations as to the underlying roles of widespread or unique natural products in the ecological niche. Such a combined approach can also be applied to microbiomes from other niches to narrow down the list of candidate BGCs that probably encode ecologically important natural products. With the functional characterization of the most conserved *XP* natural products, future detailed analysis of their targets, as well as potential synergistic/antagonistic between different compound classes (e.g., the synergistic immune suppression of GameXPeptides and lipocitides), might lead to a more comprehensive understanding of how *XP* orchestrate the interplay of natural products to maintain the symbiotic lifestyle.

## Acknowledgments

We are grateful to Prof. Jeroen S. Dickschat (University of Bonn) for providing l-[U-^34^S]-cysteine, Dr. Anja Schüffler (Institut für Biotechnologie und Wirkstoff-Forschung gGmbH) for assisting with fermentation, Stefanie Schmidt and Alexandra Amann from HIPS for testing the bioactivity of benzobactin A, Prof. Ralf-Udo Ehlers and Dr. Carlos Molina from e-nema GmbH for providing six *Photorhabdus* wild-type strains, Dr. Frank Wesche (Goethe University Frankfurt) for synthesizing 4-fluorosalicylate-SNAC, and Dr. Nick Neubacher (Goethe University Frankfurt) for constructive suggestions. Yi-Ming Shi was supported by the Alexander von Humboldt Foundation. This work is supported by the LOEWE Center for Translational Biodiversity Genomics (LOEWE TBG) to Nicholas J. Tobias and Helge B. Bode and the ERC advanced grant (835108) to Helge B. Bode.

## Author contributions

Y.-M.S. and H.B.B. designed the project. Y.-M.S. performed genome and biosynthetic gene cluster studies. Y.-M.S., M.H., D.A., J.J.C., and L.P. generated mutants. M.H., Y.-N.S, D.A., P.G., and J.J.C. performed compound isolation and structural elucidation under the guidance of Y.-M.S,. Y.-M.S. and N.J.T. conducted statistical analyses. Y.-M.S. performed the chemical synthesis of lipocitides. M.H., A.S., J.H., R.M., and Y.K. conducted bioassays. W.K. and M.G. conducted proteasome inhibition assays and complex crystallization. C.R. conducted NMR spectra measurements. A.T. conducted strain isolation. N.J.T., S.J.P., and T.P.S. performed genome sequencing, assembly, and annotation. Y.-M.S. and H.B.B. wrote the manuscript with input from all co-authors.

