## Supplementary Information for "Global analysis of biosynthetic gene clusters reveals conserved and unique natural products in entomopathogenic nematode-symbiotic bacteria"

### Methods

#### General experimental procedures

All chemicals were purchased from Sigma-Aldrich, Acros Organics, or Iris BIOTECH. Isotope-labeled chemicals were purchased from Cambridge Isotope Laboratories, Inc. Genomic DNA of selected *Xenorhabdus* and *Photorhabdus* strains were isolated using the Qiagen Gentra Puregene Yeast/Bact Kit. DNA polymerases (Taq, Phusion, and Q5) and restriction enzymes were purchased from New England Biolabs or Thermo Fisher Scientific. DNA primers were purchased from Eurofins MWG Operon. PCR amplifications were carried out on thermocyclers (SensoQuest). Polymerases were used according to the manufacturers' instructions. DNA purification was performed from 1% TAE agarose gel using Invisorb® Spin DNA Extraction Kit (STRATEC Biomedical AG). Plasmids in *E. coli* were isolated by alkaline lysis. HPLC–UV–MS analysis was conducted on an UltiMate 3000 system (Thermo Fisher) coupled to an AmaZonX mass spectrometer (Bruker) with an ACQUITY UPLC BEH C18 column (130 Å, 2.1 mm × 100 mm, 1.7 µm particle size, Waters) at a flow of 0.6 mL/min (5–95% acetonitrile/water with 0.1% formic acid, v/v, 16 min, UV detection wavelength 190–800 nm). HPLC–UV–HRMS analysis was conducted on an UltiMate 3000 system (Thermo Fisher) coupled to an Impact II qTof mass spectrometer (Bruker) with an ACQUITY UPLC BEH C18 column (130 Å, 2.1 mm × 100 mm, 1.7 µm particle size, Waters) at a flow of 0.4 mL/min (5–95% acetonitrile/water with 0.1% formic acid, v/v, 16 min, UV detection wavelength 190–800 nm). Flash purification was performed on a Biotage SP1™ flash purification system (Biotage, Uppsala, Sweden) by a $C_{18}$ main column (Interchim, PF50C18HP-F0080, 120 g) with a self-packed pre-column (Interchim, PF-DLE-F0012, Puriflash dry-load empty F0012 Flash column) coupling with a UV detector. HPLC purification was performed on preparative and semipreparative Agilent 1260 systems coupled to a DAD and a single quadrupole detector with a C18 ZORBAX Eclipse XDB column (9.4 mm × 250 mm, 5 µm, 3 mL/min; 21.2 mm × 250 mm, 5 µm, 20 mL/min; 50 mm x 250 mm, 10 µm, 40 mL/min). Freeze drying was performed by BUCHI Lyovapor™ L-300 Continuous. NMR experiments were acquired on a Bruker AVANCE 500, 600, or 700 MHz spectrometer equipped with a 5 mm cryoprobe. 2*R*,3*S*-IOC (**1**) and GameXPeptide A (**16**) were synthesized by the WuXi App Tec.

#### Genome sequencing, assembly, and annotation

Isolated DNA was sequenced on the Illumina NextSeq 500 platform. DNA libraries were constructed using the Nextera XT DNA preparation kit (Illumina) and whole-genome sequencing was performed using 2 × 150 bp paired-end chemistry. A sequencing depth of >50× was targeted for each sample. Genomes were assembled using SPAdes v. 3.10.1 [1] and annotated using Prokka v. 1.12 [2].

### antiSMASH annotations and BiG-FAM preliminary classification

antiSMASH 5.0 [3] was employed to mine all the genome sequences for the presence of putative natural product BGCs. The identified BGCs summarized for each strain (Supplementary Fig. 1) and visualized in the Anvi'o [4,5] layers (Fig. 1a and Supplementary Fig. 3). We then submitted the antiSMASH job IDs to the biosynthetic gene cluster families database (BiG-FAM) [6] for preliminary GCFs explorations and classifions of annotated BGCs (Supplementary Fig. 15), followed by BiG-SCAPE [7] refinement with a cut-off of 0.65 (Supplementary Fig. 8). The GFCs were double-checked manually via the interactive network (Fig. 2) and made corrections if necessary. A putative thiopeptide BGC (Xszus_1.region006, Xsze_2.region003, Xsto_4.region001, Xpb_30.3_21.region001, Xmir_10.region001, Xmau_6.region001, Xkoz_3.region001, Xjap_NZ_FOVO01000011.region001, Xish_1.region003, Xhom_ANU1.region005, Xhom_2.region003, Xets_11.region001, XenKK7.region002, XenDL20_c00108_NODE_12.region001, Xekj_19.region001, Xehl_28.region001, Xe30TX1_c0031_NODE_38.region001, Xdo_HBLC131_1.region001, Xdo_FRM16.1.region005, Xbov_NC_013892.1.region004, Ptem_HBLC135_17.region001, Ppb6_4.region001, Plum_TT01_1.region008, Pthr_PT1.1_23.region001, Plau_IT4.1_12.region001, Plum_IL9_35_scf0001.region001, Pbod_HU2.3_20.region001, Plau_HB1.3_105.region001, Plum_EN01_24_scf0009.region001, Pbod_DE6.1_24.region001, Plau_DE2.2_108.region001, Phpb_1.region001, Pbod_LJ_007.region001, Pbod_CN4_25_scf0020.region001, Paeg_BKT4.5_19.region001, P_tem_1.region017 et al.) that exists throughout 45 *XP* genomes was excluded in the analysis, since it turned out that its annotation by antiSMASH 5.0 is a false positive and early reports suggest that this cluster is responsible for ribosomal methylthiolation [8,9]. Two BGCs, Xdo_HBLC131_4.region001 encoding the biosynthesis of glidobactins in *X. doucetiae* HBLC131 and Ptem_HBLC135_2.region002 encoding the biosynthesis of ririwpeptides in *P. temperata* HBLC135 were artificially integrated into their respective genome by CRAGE[10] previously, and thus the two BGCs were also excluded in our analysis.

### Pangenome analysis

All genomes were obtained from the National Center for Biotechnology Information (NCBI). Supplementary Table S1 reports their accession numbers. The pangenome analysis herein mainly followed the Anvi'o v6.1 pangenomic workflow [4,5]. After simplifying the header lines of 45 FASTA files for genomes using "anvi-script-reformat-fasta", we converted FASTA files into Anvi'o contigs databases by "anvi-gen-contigs-database" and then decorated the contigs database with hits from HMM models by "anvi-run-hmms". The program "anvi-run-ncbi-cogs" was run to annotate genes in the contigs databases with functions from the NCBI's Clusters of Orthologous Groups (COGs). Tables of gene callers IDs with start and stop nucleotide positions were exported by "anvi-exporttable", by which a table with BGC classification (with a possible compound name) and boundary defined by antiSMASH were generated. The obtained table with COGs, as well as BGC and compound annotations were imported back to contigs databases by "anvi-import-functions". External genome storage was created by "anvi-gen-genomes-storage" to store DNA and amino acid sequences, as well as functional annotations of each gene. With the genome storage in hand, we used the program "anvi-pan-genome" with the genomes storage database, the flag "--use-ncbi-blast", and the parameter "--mcl-inflation 8". The results were displayed in an interface by "anvi-display-pan". The organization of gene clusters was represented by "presence/absence" patterns, as shown in the dendrogram in the center of the interface. The core gene bin was characterized by searching gene clusters using filters with "Min number of genomes gene clusters occurs, value = 45". The single-copy-core-gene (scg) bin was found by "Min number of genomes gene clusters occurs, value = 45" and "Max number of genes from each genome value = 1". The singleton bin was identified by "Max number of genomes gene clusters occurs, value = 1". The rest of the gene clusters that were neither sorted into the core gene bin and the singleton bin were appended to the accessory bins I and II. The bin summary was exported by "anvi-summarize" for further analysis (Supplementary Tables 3–5). The SCG bin was refined by "Max functional homogeneity index 0.9" and "Min geometric homogeneity index 1". The resulted protein sequences were exported by "anvi-get-sequences-for-gene-clusters" and aligned by using ClustalW which is incorporated in Geneious v6.1.8. Phylogenetic trees were generated using the Geneious tree builder utilizing the Jukes-Cantor distance model and UPGMA, and subsequently were imported back to Anvi'o by "anvi-import-misc-data" and visualized by the interface. The statistical data of BGCs obtained from antiSMASH 5.0 [3] and BiG-SCAPE [7] were import to the layers of the interface by "anvi-import-misc-data" for visualization.

### BiG-SCAPE analysis

BGCs in all genome sequences obtained from antiSMASH 5.0 [3] analyses were compared to reference BGCs from the MIBiG repository v2.0 [11,12] using BiG-SCAPE [7] with PFAM database 32.0 [13]. The analysis was conducted using default settings with mode 'auto', mixing all classes, and retaining singletons. Networks were computed for raw distance cut-offs of 0.30–0.95 in increments of 0.05. Results were visualized as a network using Cytoscape 3.7.2 [14] for a cut-off of 0.65 (Fig. 2 and Supplementary Table 8).

### Strain and culture conditions

Wild-type strains and the mutants thereof and *E. coli* (Supplementary Table 16) were cultivated on lysogeny broth (LB) agar plates at 30 $^{o}$C overnight and were subsequently inoculated into liquid LB culture at 30 $^{o}$C with shaking at 200 rpm. For compound production, the overnight LB culture was transferred into 5 mL LB, XPP[15], or Sf-900$^{TM}$ II SFM medium (1:100, v/v) with 2% (v/v) of

Amberlite™ XAD-16 resins, 0.1 % of L-arabinose as the inducer (for mutants with a $P_{BAD}$ promoter), and selective antibiotics such as ampicillin (Am, 100 µg/mL), kanamycin (Km, 50 µg/mL), or chloramphenicol (Cm, 34 µg/mL) at 30 °C with shaking at 200 rpm.

### Culture extraction and HPLC-UV-MS analysis

The XAD-16 resins were collected after 72 h and extracted with 5 mL methanol or ethyl acetate. The solvent was dried under rotary evaporators, and the dried extract was resuspended in 500 µL methanol or acetonitrile/water (1:1, v/v, for photoxenobactins), of which 5 µL was injected and analyzed by HPLC-UV-MS or HPLC-UV-HRMS. Unless otherwise specified, HPLC-UV-MS and HPLC-UV-HRMS chromatograms in the figures were shown on the same scale.

### Construction of insertion mutants

A 500–800-bp upstream of the target gene (*lpcS*, *pxbF*, *rdb1A*, and *xvbA*) was amplified with a corresponding primer pair listed in Supplementary Table 18. The resulting fragments were cloned using Hot Fusion [16] into pCEP_kan or pCEP_cm backbone that was amplified by pCEP_Fw and pCEP_Rv. After the transformation of a constructed plasmid into *E. coli* S17-1 *λ* pir, clones were verified by PCR with primers pCEP-Ve-Fw and pDS132-Ve-Rv. A wild-type strain (recipient) was mated with *E. coli* S17-1 *λ* pir (donor) carrying a constructed plasmid. Both strains were grown in the LB medium to an $OD_{600}$ of 0.6 to 0.7, and the cells were washed once with the fresh LB medium. Subsequently, the donor and recipient strains were mixed on an LB agar plate in ratios of 1:3 and 3:1, and incubated at 37°C for 3 h followed by incubation at 30°C for 21 h. After that, the bacterial cell layer was harvested with an inoculating loop and resuspended in 2 mL fresh LB medium. 200 µL of the resuspended culture was spread out on an LB agar plate with Am/Km or Am/Cm incubated at 30°C for 2 days. Individual insertion clones were cultivated and analyzed by HPLC-UV-HRMS, and the genotype of all mutants was verified by plasmid- and genome-specific primers.

### Construction of deletion mutants

A ~1,000-bp upstream and a ~1,000-bp downstream fragments of a target gene or region (*rdb1P*) were amplified using primer pairs listed in Supplementary Table 18. The amplified fragments were fused using the complementary overhangs introduced by primers and cloned into the pCKcipB vector that was linearized with PstI and BglII) by Hot Fusion [16]. Transformation of *E. coli* S17-1 *λ* pir with the resulting plasmid and conjugation with a wild-type strain, as well as the generation of double crossover mutants via counterselection on LB plates containing 6% sucrose, were done as previously described [17]. The deletion mutant was verified via PCR using primer pairs listed in Supplementary Table 18, which yielded a ~2,000-bp fragment for mutants genetically equal to the WT strain and a ~1,000-bp fragment for the desired deletion mutant.

### Labeling experiments for structural elucidation of photoxenobactin C by MS

The cultivation of strains for labeling experiments was carried out as described above. The overnight culture was transferred into LB medium additionally fed with 4-fluorosalicylate-SNAC, L-methionine-(methyl-$d_3$), L-[U-$^{13}$C,$^{15}$N]cysteine, and L-[U-$^{34}$S]cysteine at a final concentration of 1 mM. In terms of inverse feeding experiments, the cell pellets of the 100 µL overnight culture were washed once with ISOGRO $^{13}$C or $^{13}$C,$^{15}$N medium (100 µL), and resuspended in corresponding medium (100 µL). The feeding culture in isotope labeling medium (5 mL) was inoculated with a washed overnight culture (50 µL) and additional L-cysteine was added at a final concentration of 1 mM.

### Isolation and purification

In terms of photoxenobactin isolation, 10 mL LB medium was inoculated with a colony of the *X. szentirmaii* $P_{BAD}$ *pxbF* Δ*hfq* from an LB agar plate and cultivated overnight. The 10 mL culture was taken to inoculate 2 x 100 mL LB medium with an $OD_{600}$ ≈ 0.1. The 2 x 100 mL cultures were incubated overnight and the whole culture volume (200 mL) was used to inoculate a 20 L LB fermenter (Braun, Melsungen) supplemented with 2% XAD-16 and 0.2% arabinose (antifoam was added when required). Fermenter settings were as follows: 30°C without pH control, three six-blade impellers 150 rpm. After 24 h, 10 L the culture was collected from the fermenter, and the XAD resins were separated from the cells by filtration. 1) The XAD resins were extracted with 2 x 1 L of ethyl acetate with 1% formic acid; the combined organic phase was dried under reduced pressure. 2) The culture without XAD was centrifuged and the supernatant was extracted 3 x 5 L ethyl acetate with 1% formic acid; the combined organic layers were dried under reduced pressure. 3) The cell pellet was extracted with 2 x 1 L ethyl acetate with 1% formic acid; the organic supernatant was dried under reduced pressure. After 48 h the remaining 10 L of bacterial culture were extracted as described in 1)-3). The combined extracts from 20 L culture were fractionated by a flash purification system with a $C_{18}$ column with a gradient elution of acetonitrile/water 20-100% at 20 mL/min (every ten percent gradient step was performed with 5 column volume, except the 60-70% step with 10 column volume). Fractions containing photoxenobactins were combined and dried under reduced pressure. Final purification was achieved via preparative and semipreparative HPLCs with a gradient of 30% acetonitrile/water (0-30 min) and 30-100% acetonitrile/water (30-40 min). The fractions were combined in brown flasks and were immediately freeze-dried to afford photoxenobactin A (**4**, 0.8 mg), photoxenobactin B (**5**, 0.6 mg), photoxenobactin C (**6**, 1.2 mg), and photoxenobactin E (**8**, 2.2 mg).

For the isolation and purification of lipocitides A and B, 2% of XAD-16 resins from a 6 L LB culture of the *X. bovienii* $P_{BAD}$ *lpcS* mutant induced by L-arabinose were harvested after 72 h of incubation at 30 °C with shaking at 120 rpm, and were washed with water and extracted with methanol (3 × 1 L) to yield a crude extract 5.3 g after evaporation. The extract was dissolved in methanol and was subjected to preparative HPLC with a C18 column using an acetonitrile/water gradient (0.1% formic acid) 0–32 min, 55–80%, 40 mL/min to afford lipocitides A (**17**, 4.8 mg) and B (**18**, 9.0 mg).

2% of XAD-16 resins from a 12 L LB culture of the *X. budapestensis* $P_{BAD}$ *rdb1A* Δ*rdb1P* Δ*hfq* mutant induced by L-arabinose were harvested after 72 h of incubation at 30 °C with shaking at 120 rpm, and were washed with water and extracted with methanol (3 × 2 L) to yield a crude extract 15.3 g after evaporation. The extract was subject to a Sephadex LH-20 column eluted with methanol. The fraction (2.8 g) containing pre-rhabdobranins was subjected to preparative HPLC with a C18 column using an acetonitrile/water gradient (0.1% formic acid) 0–20 min, 15–35%, 40 mL/min to afford a fraction (206 mg) mainly containing pre-rhabdobranin D, which was further purified by semipreparative HPLC with a C18 column using an acetonitrile/water gradient (0.1% formic acid) 0–24 min, 5–53%, 3 mL/min to afford pre-rhabdobranin D (**27**, 59.1 mg)

Benzobactin A (**28**) and its methyl ester (**29**) that were detected in *X. vietnamensis* $P_{BAD}$ *xvbA* also were produced by *Pseudomonas chlororaphis* subsp. *piscium* DSM 21509 (unpublished). Due to the high production level in *Pseudomonas chlororaphis* subsp. *piscium* DSM 21509, **28** and **29** were isolated from the *Pseudomonas* strain. 4% of XAD-16 resins from a 12 L XPP culture of *Pseudomonas chlororaphis* subsp. *piscium* DSM 21509 $P_{BAD}$ *pbzA* mutant induced by L-arabinose were harvested after 72 h of incubation at 30 °C with shaking at 120 rpm, and were washed with water and extracted with methanol (3 × 2 L) to yield a crude extract 95.4 g after evaporation. The extract was dissolved in methanol and was subjected to preparative HPLC with a C18 column using an acetonitrile/water gradient (0.1% formic acid) 0–18 min, 5–59%, 20 mL/min to afford ten fractions. Fractions 2 (95.6 mg) and 3 (50.7 mg) were further purified by semipreparative HPLC with a C18 column using an acetonitrile/water gradient (0.1% formic acid) 0–35 min, 5–95%, 3 mL/min to afford benzobactin A (**28**, 3.2 mg) and its methyl ester (**29**, 0.9 mg), respectively.

### NMR spectroscopy

$^{1}H$ and $^{13}C$ NMR, $^{1}H$-$^{13}C$ heteronuclear single quantum coherence (HSQC), $^{1}H$-$^{13}C$ heteronuclear multiple bond correlation (HMBC), $^{1}H$-$^{1}H$ correlation spectroscopy (COSY), $^{1}H$-$^{13}C$ heteronuclear multiple quantum correlation/$^{1}H$-$^{1}H$ correlation spectroscopy (HMQC-COSY), and $^{1}H$-$^{13}C$ heteronuclear single quantum coherence/$^{1}H$-$^{1}H$ total correlation spectroscopy (HSQC-TOCSY) were measured. Chemical shifts ($\delta$) were reported in parts per million (ppm) and referenced to the solvent signals. Data are reported as follows: chemical shift, multiplicity (br = broad, s = singlet, d = doublet, t = triplet, dd = doublet of doublet, m = multiplet, and ov = overlapped), and coupling constants in Hertz (Hz).

### $IC_{50}$ value determination with the purified yeast 20S proteasome core particle (yCP)

The concentration of purified yCP was determined spectrophotometrically at 280 nm. yCP (final concentration: 0.05 mg/mL in 100 mM Tris-HCl, pH 7.5) was mixed with DMSO as a control or serial dilutions of IOC (**1**) in DMSO, thereby not surpassing a final concentration of 10% (v/v) DMSO. After an incubation time of 45 min at RT, fluorogenic substrates Boc-Leu-Arg-Arg-AMC, Z-Leu-Leu-Glu-AMC, and Suc-Leu-Leu-Val-Tyr-AMC (final concentration of 200 µM) were added to measure the residual activity of caspase-like (C-L, β1 subunit), trypsin-like (T-L, β2 subunit), and chymotrypsin-like (ChT-L, β5 subunit), respectively. The assay mixture was incubated for another 60 min at RT and afterward diluted 1:10 in 20 mM Tris-HCl, pH 7.5. The AMC-molecules released by hydrolysis were measured in triplicate with a Varian Cary Eclipse Fluorescence Spectrophotometer (Agilent Technologies) at $\lambda_{exc}$ = 360 nm and $\lambda_{em}$ = 460 nm. Relative fluorescence units were normalized to the DMSO treated control. The calculated residual activities were plotted against the logarithm of the applied inhibitor concentration and fitted with GraphPad Prism 9. Half maximum inhibitory concentration ($IC_{50}$) values were deduced from the fitted data. They depend on enzyme concentration and are comparable within the same experimental settings.

#### Crystallization and structure determination of the yCP in complex with IOC (1).

Crystals of the yCP were grown in hanging drops at 20 °C as previously described[18,19].The protein concentration used for crystallization was 40 mg/mL in Tris / HCl (20 mM, pH 7.5) and EDTA (1 mM). The drops contained 1 µL of protein and 1 µL of the reservoir solution [30 mM magnesium acetate, 100 mM 2-(N-morpholino)ethanesulfonic acid (pH 6.7) and 10% (wt/vol) 2-methyl-2,4-pentanediol]. Crystals appeared after two days and were incubated with **1** at final concentrations of 10 mM for at least 24 h. Droplets were then complemented with a cryoprotecting buffer [30% (wt/vol) 2-methyl-2,4-pentanediol, 15 mM magnesium acetate, 100 mM 2-(N-morpholino)ethanesulfonic acid, pH 6.9] and vitrified in liquid nitrogen. The dataset from the yCP:IOC complex was collected using synchrotron radiation (λ =1.0 Å) at the X06SA-beamline (Swiss Light Source, Villingen, Switzerland). X-ray intensities and data reduction were evaluated using the XDS program package (as the table shown below) [20]. Conventional crystallographic rigid body, positional, and temperature factor refinements were carried out with REFMAC5 [21] using coordinates of the yCP structure as starting model (PDB ID 5CZ4) [22]. For model building, the programs SYBYL and COOT [23] were used. The final coordinates yielded excellent R factors, as well as geometric bond and angle values. Coordinates were confirmed to fulfill the Ramachandran plot and have been deposited in the RCSB (PDB ID 7O2L)

Crystallographic data collection and refinement statistics.

| | ***yCP:IOC*** |
|---|---|
| **Crystal parameters** | |
| Space group | $P2_1$ |
| Cell constants | a = 135.1 Å |
| | b = 301.5 Å |
| | c = 144.3 Å |
| | β = 112.9 ° |
| CPs / AU[a] | 1 |

| | |
|---|---|
| **Data collection** | |
| Beam line | X06SA, SLS |
| Wavelength (Å) | 1.0 |
| Resolution range (Å)[b] | 30–3.0 (3-1–3.0) |
| No. observations | 635463 |
| No. unique reflections[c] | 203696 |
| Completeness (%)[b] | 96.2 (98.6) |
| $R_{merge}$ (%)[b, d] | 8.5 (57.7) |
| I/ σ (I)[b] | 9.1 (2.6) |
| **Refinement (REFMAC5)** | |
| Resolution range (Å) | 30-3.0 |
| No. refl. working set | 193349 |
| No. refl. test set | 10176 |
| No. non hydrogen | 49509 |
| No. of ligand atoms | 66 |
| Solvent ($H_2O$, ions, MES) | 138 |
| $R_{work}/R_{free}$ (%)[e] | 17.8 / 21.5 |
| r.m.s.d. bond (Å) / angle (°)[f] | 0.002 / 1.2 |
| Average B-factor ($Å^2$) | 90.5 |
| Ramachandran Plot (%)[g] | 97.4 / 2.3 / 0.3 |
| PDB accession code | 7O2L |

[a] Asymmetric unit
[b] The values in parentheses for resolution range, completeness, $R_{merge}$ and I/σ (I) correspond to the highest resolution shell
[c] Data reduction was carried out from a single crystal. Friedel pairs were treated as identical reflections
[d] $R_{merge}(I) = \Sigma_{hkl}\Sigma_j \mid I(hkl)_j - \langle I(hkl)\rangle \mid / \Sigma_{hkl} \Sigma_j I(hkl)_j$, where $I(hkl)_j$ is the $j^{th}$ measurement of the intensity of reflection hkl and <I(hkl)> is the average intensity
[e] $R = \Sigma_{hkl} \mid |F_{obs}| - |F_{calc}| \mid / \Sigma_{hkl} |F_{obs}|$, where $R_{free}$ is calculated without a sigma cut off for a randomly chosen 5% of reflections, which were not used for structure refinement, and $R_{work}$ is calculated for the remaining reflections
[f] Deviations from ideal bond lengths/angles
[g] Percentage of residues in favored / allowed / outlier region

### Hemocyte-spreading assay

*Spodoptera exigua* larvae were collected from Welsh onion (*Allium fistulsum* L.) fields in Andong, Korea. Insects were reared in the laboratory under conditions of 25 ± 2°C constant temperature, 16:8 h (L: D) photoperiod, and 60 ± 5 % relative humidity. Larvae were reared on an artificial diet [24] and 10% sucrose solutions were fed to adult insects. Fifth instar larvae were used to conduct all experiments. For analyzing hemocyte behaviors in vivo, fifth instar larvae of *S. exigua* were co-injected with 1 µL of heat-killed (95 $^{o}$C for 10 min) *E. coli* TOP10 ($2.4 \times 10^4$ cells/larva) with the test compound (0–1,000 ng/larva) by using a Hamilton microsyringe (Reno, NV, USA). After 1 h post-injection, 10 µL of hemolymph from each larva was collected on the glass slide and incubated for 5 min inside a dark wet chamber at room temperature (RT). The medium was replaced with 3.7% of formaldehyde which was dissolved in PBS and was incubated for 10 min. After washing three times with PBS, cells were permeabilized with 0.2% Triton X-100 in PBS for 2 min at RT. After incubation, slides were washed with PBS three times. Blocking was performed by using 5% skimmed milk (Invitrogen, Carlsbad, CA, USA) which was dissolved in PBS and incubated for 10 min. After washing once with PBS, cells were incubated with fluorescein isothiocyanate (FITC)-tagged phalloidin in PBS for 1 h at RT. After washing three times, cells were incubated with 4',6-diamidino-2-phenylindole (DAPI, 1 mg/mL, Thermo Scientific, Rockford, IL, USA) in PBS for nucleus staining. Finally, after washing twice in PBS, cells were observed under a fluorescence microscope (DM2500, Leica, Wetzlar, Germany) at 400 × magnification. Hemocyte-spreading was determined by the extension of F-actin out of the original cell boundary. For in vitro assay, ~100 µL of hemolymph were collected into 400 µL of anticoagulation buffer (ACB: 186 mM NaCl, 17 mM $Na_2EDTA$, 41 mM citric acid, pH 4.5). After adding ACB, the medium was incubated for 30 min on ice. After centrifugation at 300 × *g* for 5 min, 400 µL of supernatant was discarded. The rest of the suspension was gently mixed with 200 µL of TC100 insect tissue culture medium (Welgene, Gyeongsan, Korea). From this suspension, 10 µL of hemolymph was collected on the glass slide. The slides were co-injected with 1 µL of *E. coli* TOP10 ($2.4 \times 10^4$ cells/larva) with the test compound (0–1,000 ng/larva), followed by the procedure described above.

### Nodulation assay

*E. coli* TOP10 was heat-killed by incubating at 95°C for 10 min. Fifth instar larvae of *S. exigua* were injected with 1 µL of bacteria ($2.4 \times 10^4$ cells/larva) using a Hamilton microsyringe along with 1 µL of different concentrations (10, 50, 100, 500, and 1,000 ppm) of inhibitors. Control larvae were injected with bacteria and DMSO. At 8 h after bacterial injection, nodules were counted by dissecting larvae under a stereomicroscope (Stemi SV 11, Zeiss, Jena, Germany) at 50 × magnifications.

#### Phenoloxidase (PO) activity assay

PO activity from plasma was estimated as previously described [25]. Briefly, DOPA (L-3,4-dihydroxyphenylalanine) was used as a substrate for determining PO activity from treated larvae plasm. For PO activation, each fifth instar larvae of *S. exigua* was challenged with $2.4 \times 10^4$ cells of heat-killed *E. coli* TOP10. Different inhibitors were co-injected (1 µg/larvae) along with *E. coli* TOP10. After 8 h of bacterial challenge, hemolymph was collected from treated larvae in 1.5 mL tube containing few granules of phenylthiocarbamide (Sigma-Aldrich Korea, Seoul, Korea) to prevent melanization. Hemocytes were separated from plasma by centrifuging at 4°C for 5 min at 300 × g. A reaction volume of 200 µL consisted of 180 µL of 10 mM DOPA in PBS (pH 7.4) and 20 µL of plasma. Absorbance was taken using VICTOR multi-label Plate reader (PerkinElmer, Waltham, MA, USA) at 490 nm. PO activity was expressed as ΔABS/min/µL of plasma. Each treatment was replicated three times with independent samples.

#### Measurement of nitric oxide (NO)

NO was indirectly quantified by measuring its oxidized form, nitrate ($NO_2^-$), using the Griess reagent of the Nitrate/Nitrite Colorimetric Assay Kit (Cayman Chemical, Ann Arbor, MI, USA). Fifth instar larvae were injected with 1 µL of heat-killed *E. coli* TOP10 ($2.4 \times 10^4$ cells/larva) using a Hamilton microsyringe along with 1 µL of the test compound. Hemolymph was collected from each sample after 1 h post-infection. 150 µL of hemolymph from three L5 larvae was collected and homogenized in 350 µL of 100 mM phosphate-buffered saline (PBS, pH 7.4) with a homogenizer (Ultra-Turrax T8, Ika Laboratory, Funkentstort, Germany). After centrifugation at 14,000 × *g* for 20 min at 4°C, the supernatant was used to measure the nitrate amounts, and the total protein was measured in each sample by Bradford assay. The samples were analyzed in a 200 µL final reaction volume. Briefly, 80 µL of samples was added to the wells, and then 10 µL of enzyme cofactor mixture and 10 µL of nitrate reductase mixture were added. After incubation at RT for 1 h, 50 µL of Griess reagent R1 and immediately 50 µL of Griess reagent R2 were added to each well. The plate was left at RT for 10 min for color development. For a standard curve to quantify nitrate concentrations of the samples, nitrates with final concentrations of 0, 5, 10, 15, 20, 25, 30, and 35 µM in 200 µL reaction volume were used. The absorbance was recorded at 540 nm on a VICTOR multi-label Plate reader. Our measurements used three larvae per sample, and we repeated the treatment with three biological samples.

#### Galleria injection assay

Precultures of *X. szentirmaii* DSM wild-type strain and the mutants thereof were grown in LB medium and inoculated into fresh cultures at an $OD_{600}$ of 0.1. Cells were grown to exponential phase ($OD_{600} \approx 1$) and then diluted to an $OD_{600}$ to 0.00025. 5 µL of the diluted bacterial culture was injected into the last left pro-leg of the larvae (LB medium as a negative control). *G. mellonella* larvae were kept at 4°C for 10 min prior to injection. After infection, the larvae were incubated at 25°C. Dead *Galleria* larvae were frozen at -20°C, then at -80°C, and freeze-dried for 1 day. Freeze-dried larvae were ground. Every injection experiment was aliquoted into two portions, one of which was extracted with 25 mL acetone/ethyl acetate (v/v, 1:1) while the other one was extracted with acetone/methanol. Extracts were dried and resuspended in 3 mL acetonitrile/water (1:1, v/v) with tenfold dilution for HPLC-MS-UV analysis.

#### Data availability

All data generated or analyzed in this study are available within the article and its Supplementary Information files.

**a**

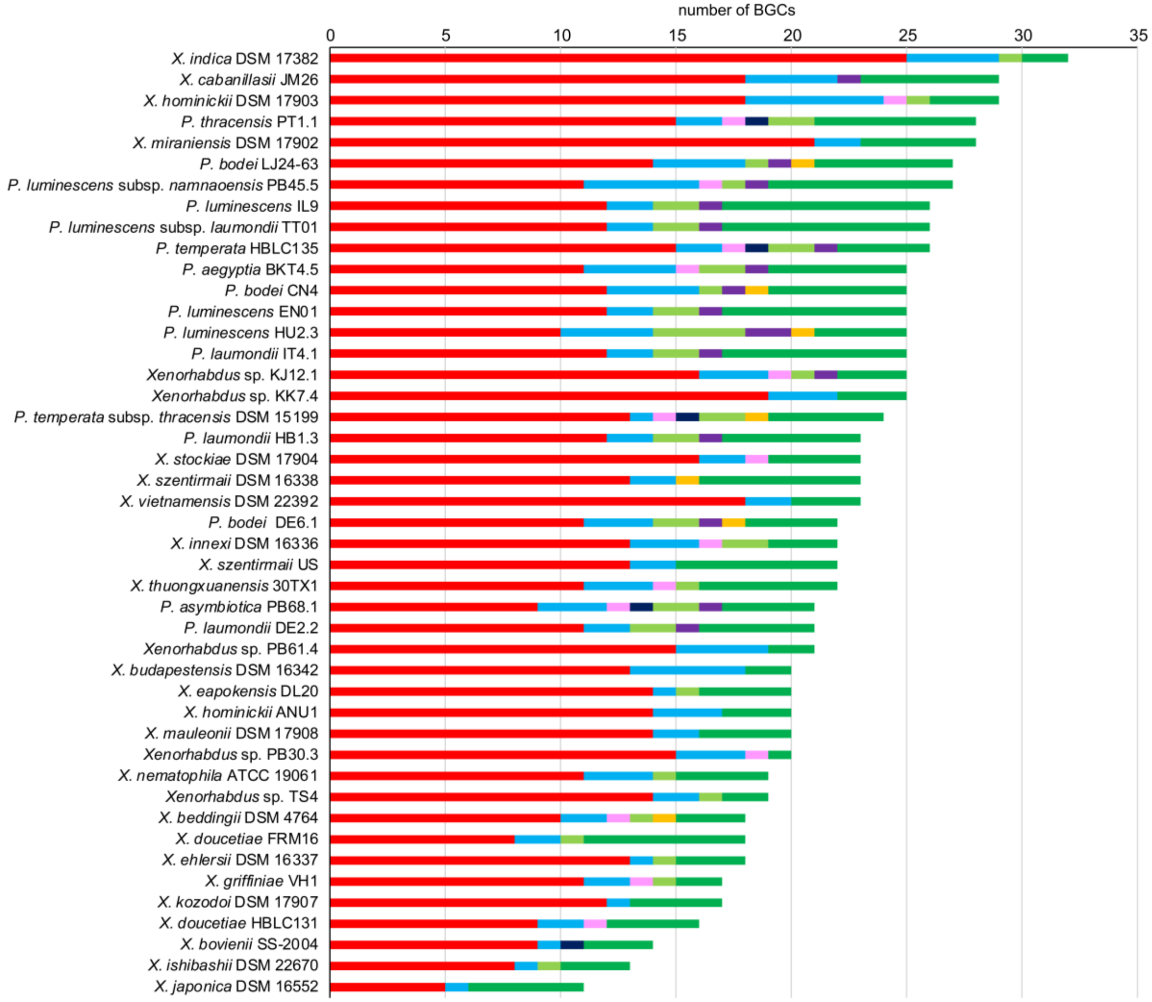
number of BGCs
0
5
10
15
20
25
30
35
X. indica DSM 17382
X. cabanillasii JM26
X. hominickii DSM 17903
P. thracensis PT1.1
X. miraniensis DSM 17902
P. bodei LJ24-63
P. luminescens subsp. namnaoensis PB45.5
P. luminescens IL9
P. luminescens subsp. laumondii TT01
P. temperata HBLC135
P. aegyptia BKT4.5
P. bodei CN4
P. luminescens EN01
P. luminescens HU2.3
P. laumondii IT4.1
Xenorhabdus sp. KJ12.1
Xenorhabdus sp. KK7.4
P. temperata subsp. thracensis DSM 15199
P. laumondii HB1.3
X. stockiae DSM 17904
X. szentirmaii DSM 16338
X. vietnamensis DSM 22392
P. bodei DE6.1
X. innexi DSM 16336
X. szentirmaii US
X. thuongxuanensis 30TX1
P. asymbiotica PB68.1
P. laumondii DE2.2
Xenorhabdus sp. PB61.4
X. budapestensis DSM 16342
X. eapokensis DL20
X. hominickii ANU1
X. mauleonii DSM 17908
Xenorhabdus sp. PB30.3
X. nematophila ATCC 19061
Xenorhabdus sp. TS4
X. beddingii DSM 4764
X. doucetiae FRM16
X. ehlersii DSM 16337
X. griffiniae VH1
X. kozodoi DSM 17907
X. doucetiae HBLC131
X. bovienii SS-2004
X. ishibashii DSM 22670
X. japonica DSM 16552

b

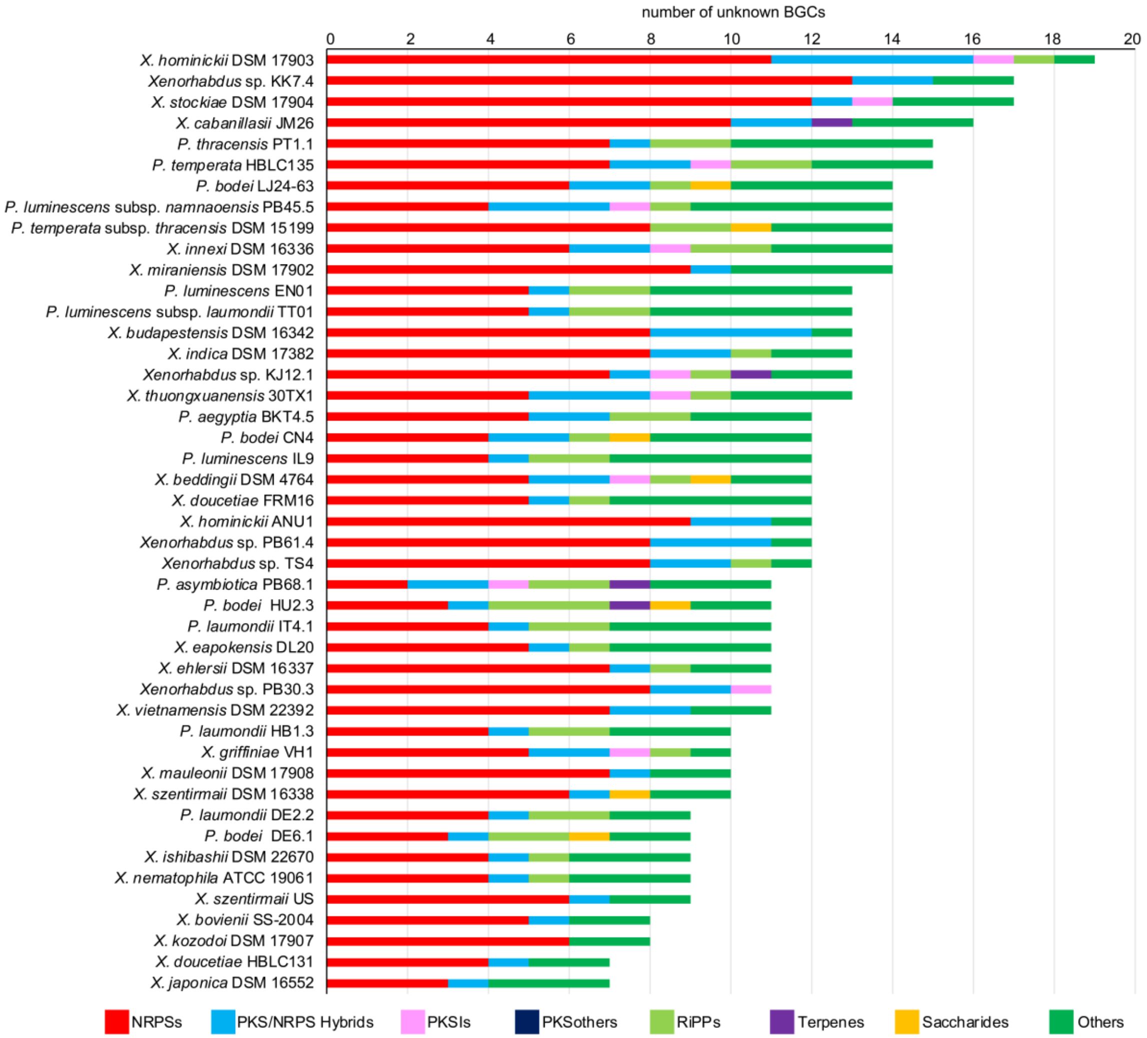

**Supplementary Fig. 1 | The number and classes of (a) total BGCs (including fragments) annotated by antiSMASH 5.0 and (b) unknown BGCs in each *XP* genome.**

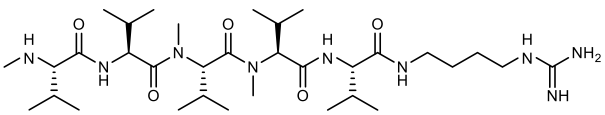

rhabdopeptide/xenortide-like peptide

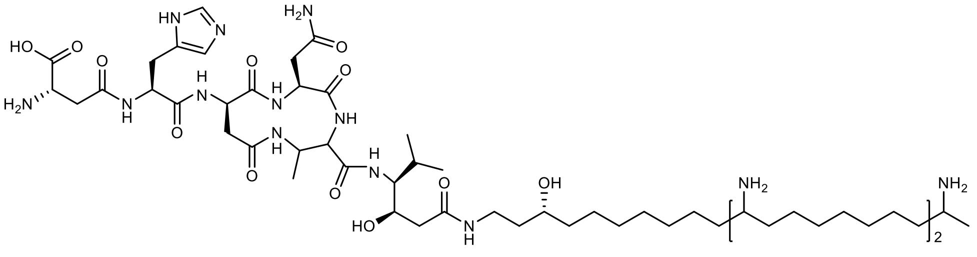

fabclavine IVa

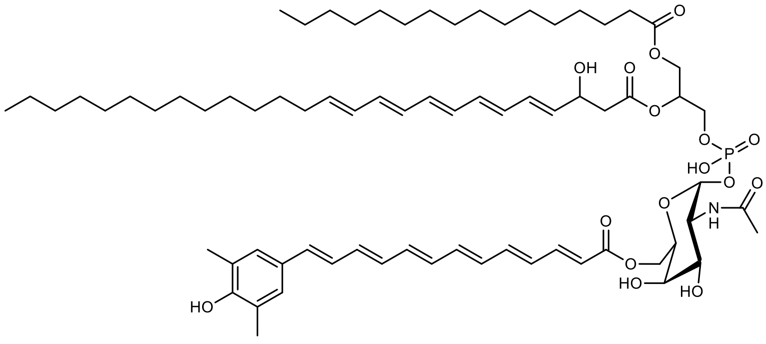

APEL-1284

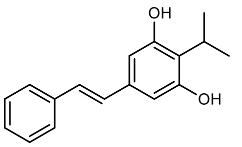

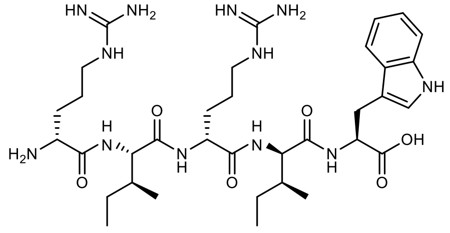

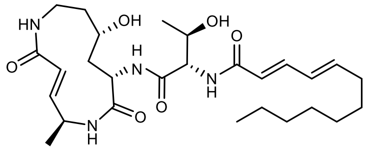

isopropylstilbene ririwpeptide A glidobactin A

**Supplementary Fig. 2 | Known natural products widely distributed in *Xenorhabdus* and/or *Photorhabdus*.** Rhabdopeptide/xenortide-like peptides [26] are the second most broadly distributed non-ribosomal peptide in *XP*. Fabclavines [27] are the most prevalent *Xenorhabdus*-specific polyketide/non-ribosomal peptide hybrid. APELs [28] are the most prominent compound class in Gram-negative bacteria [29] and exclusively present in *Xenorhabdus* but absent in *Photorhabdus*. Isopropylstilbene [30] is highly conserved across all *Photorhabdus*. Ririwpeptides [10] and glidobactins[31] are the most widespread *Photorhabdus*-specific non-ribosomal peptide and polyketide/non-ribosomal peptide hybrid, respectively.

**a**

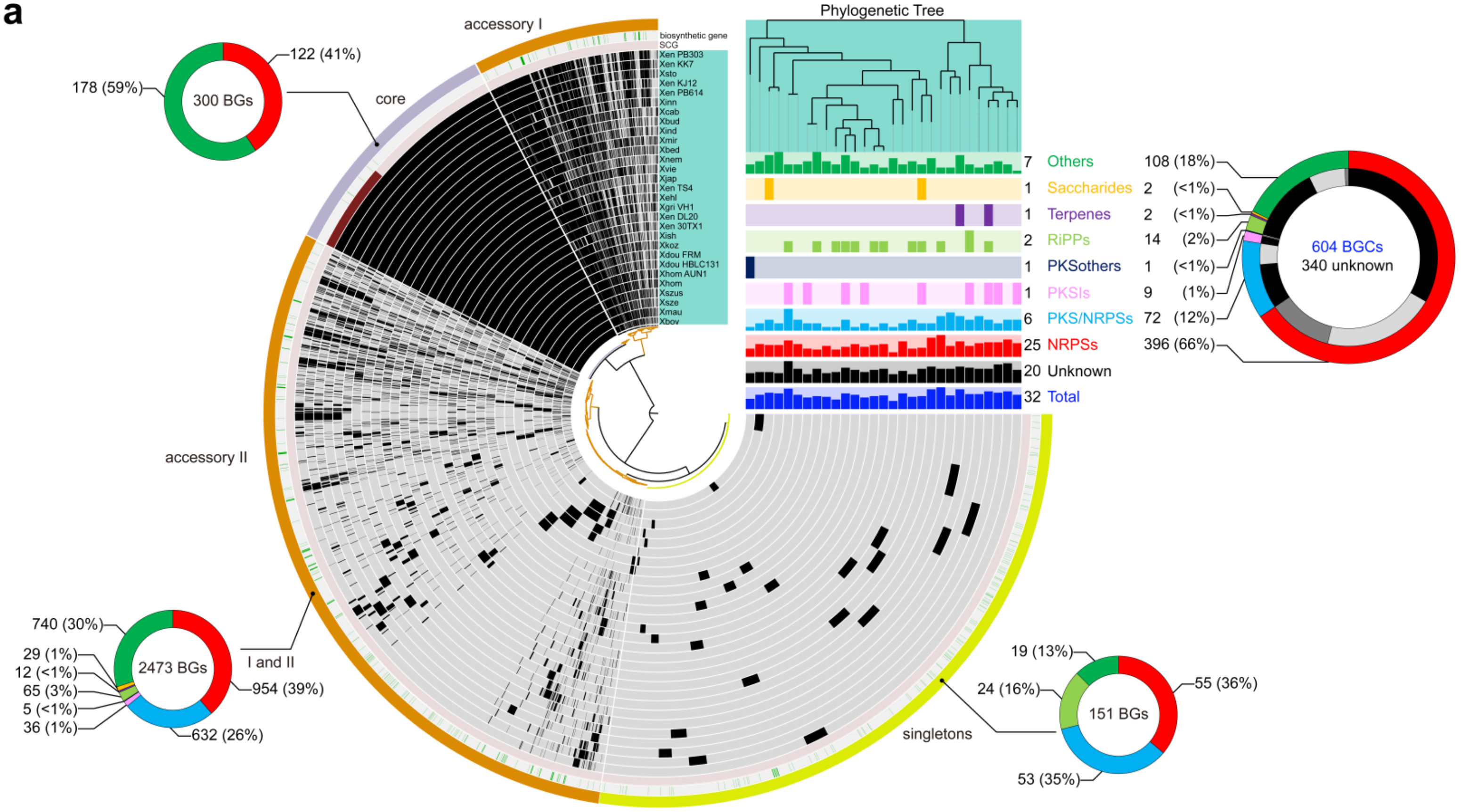
accessory I
core
accessory II
singletons
I and II
Phylogenetic Tree
178 (59%)
300 BGs
122 (41%)
740 (30%)
29 (1%)
12 (<1%)
65 (3%)
5 (<1%)
36 (1%)
2473 BGs
954 (39%)
632 (26%)
19 (13%)
24 (16%)
151 BGs
55 (36%)
53 (35%)
7 Others 108 (18%)
1 Saccharides 2 (<1%)
1 Terpenes 2 (<1%)
2 RiPPs 14 (2%)
1 PKSothers 1 (<1%)
1 PKSIs 9 (1%)
6 PKS/NRPSs 72 (12%)
25 NRPSs 396 (66%)
20 Unknown
32 Total
604 BGCs
340 unknown
biosynthetic gene
SCG
Xen PB303
Xen KK7
Xsto
Xen KJ12
Xen PB614
Xinn
Xcab
Xbud
Xind
Xmir
Xbed
Xnem
Xvie
Xjap
Xen TS4
Xehl
Xgri VH1
Xen DL20
Xen 30TX1
Xish
Xkoz
Xdou FRM
Xdou HBLC131
Xhom AUN1
Xhom
Xszus
Xsze
Xmau
Xbov

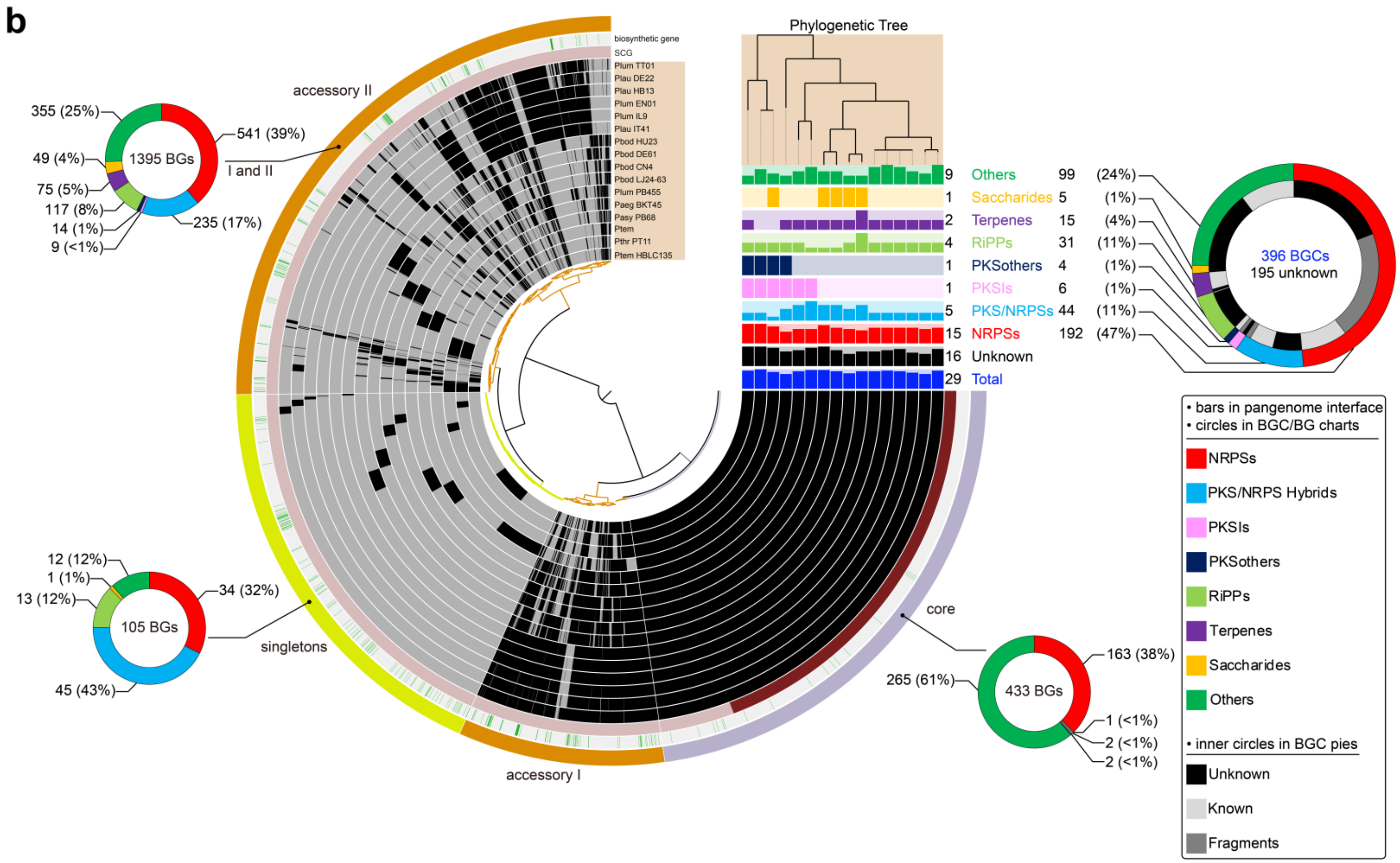

**Supplementary Fig. 3 | Pangenomic and BGs/BGCs analysis of 29 *Xenorhabdus* and 16 *Photorhabdus* genomes.** Overview of classifications and distribution of BGs/BGCs in (**a**) 29 *Xenorhabdus* and (**b**) 16 *Photorhabdus*. In the circle interface, each layer (grey) represents all genes (black) in a single genome; single-copy-core-gene (scg, dark red), biosynthetic gene (green), bin names (core, grey-purple; accessories I and II, orange; singletons, yellow). The number and classification of BGCs in a strain is represented by the bar charts under the phylogenetic tree. The maximum number of each BGC class is indicated on the right side of the bar charts. The number of identified BGCs in total, unknown BGCs, and BGs in different pangenomic regions are indicated by doughnut charts.

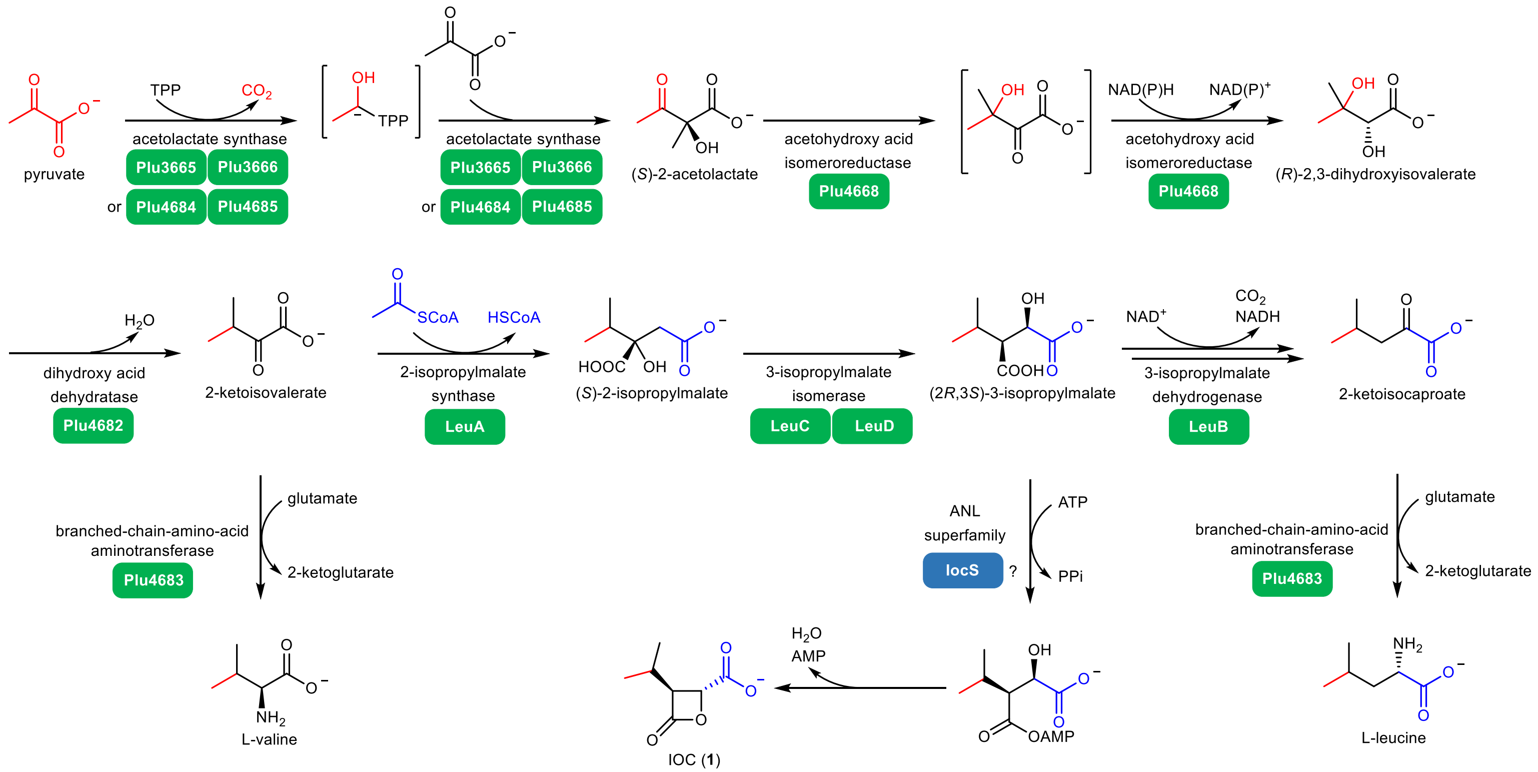

**Supplementary Fig. 4 | Proposed biosynthetic pathway of IOC (1) in *P. luminescens* subsp. *laumondii* TT01.**

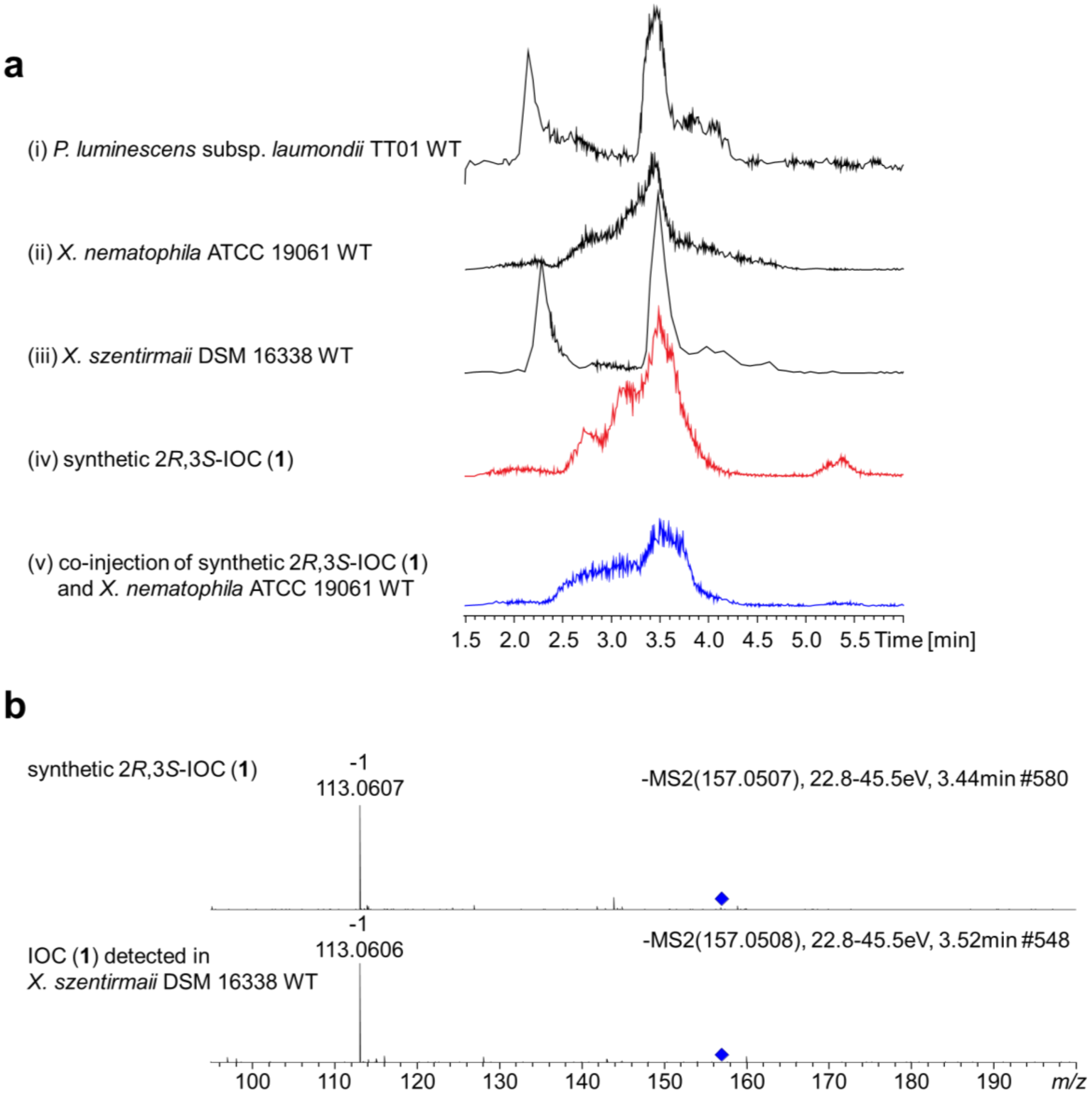

**Supplementary Fig. 5 | HPLC-MS analysis of IOC in different *Xenorhabdus* and *Photorhabdus* strains in Sf-900 medium. a,** 157.05075 $[M – H]^-$ EICs of the culture supernatants of (i) *P. luminescens* subsp. *laumondii* TT01 WT, (ii) *X. nematophila* ATCC 19061 WT, and (iii) *X. szentirmaii* DSM 16338 WT, as well as (iv) synthetic 2*R*,3*S*-IOC (**1**) and (v) co-injection of synthetic 2*R*,3*S*-IOC (**1**) and the culture supernatant of *X. nematophila* ATCC 19061 WT. **b,** Comparison of the MS/MS fragmentation patterns of the synthetic 2*R*,3*S*-IOC (**1**) and IOC (**1**) detected in the culture supernatant of *X. szentirmaii* DSM 16338 WT. The blue diamond indicates the parent ions. Representative data from three independent experiments are shown.

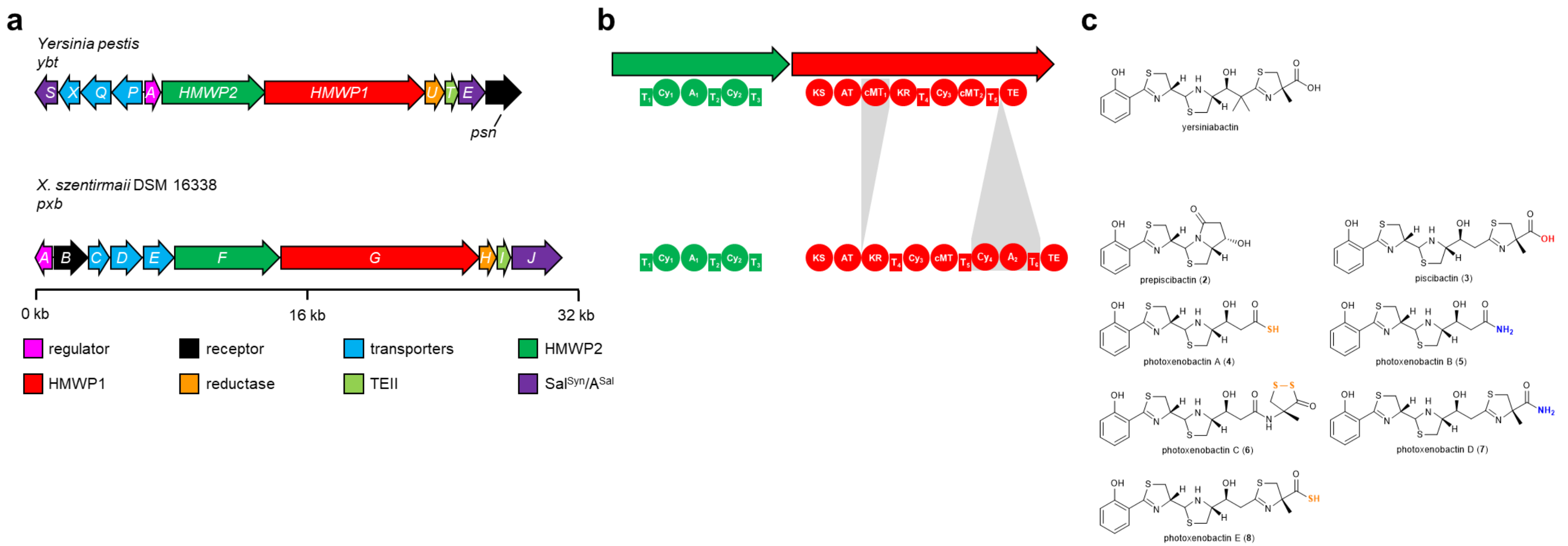

**Supplementary Fig. 6 | BGCs and chemical structures of yersiniabactin and photoxenobactins. a,** Comparison of yersiniabactin-related BGCs in *Yersinia pestis* (*ybt*) and *X. szentirmaii* (*pxb*). kb, kilobase. **b,** Domain organization of HMWP1 and HMWP2 homologs encoded by two BGCs. Domain differences are indicated with shades of gray. T, thiolation; A, adenylation; Cy, heterocyclization; KS, ketosynthase; AT, acyltransferase; KR, ketoreductase; cMT, carbon methyltransferase; TE, thioesterase. **c,** Known chemical structures, yersiniabactin from *Y. pestis* and prepiscibactin (**2**) and piscibactin (**3**) from *Photobacterium damselae* subsp. *piscida*[32], as well as previously unidentified photoxenobactins A-E (**4**–**8**) from *X. szentirmaii* DSM 16338. The terminal heteroatoms are highlighted.

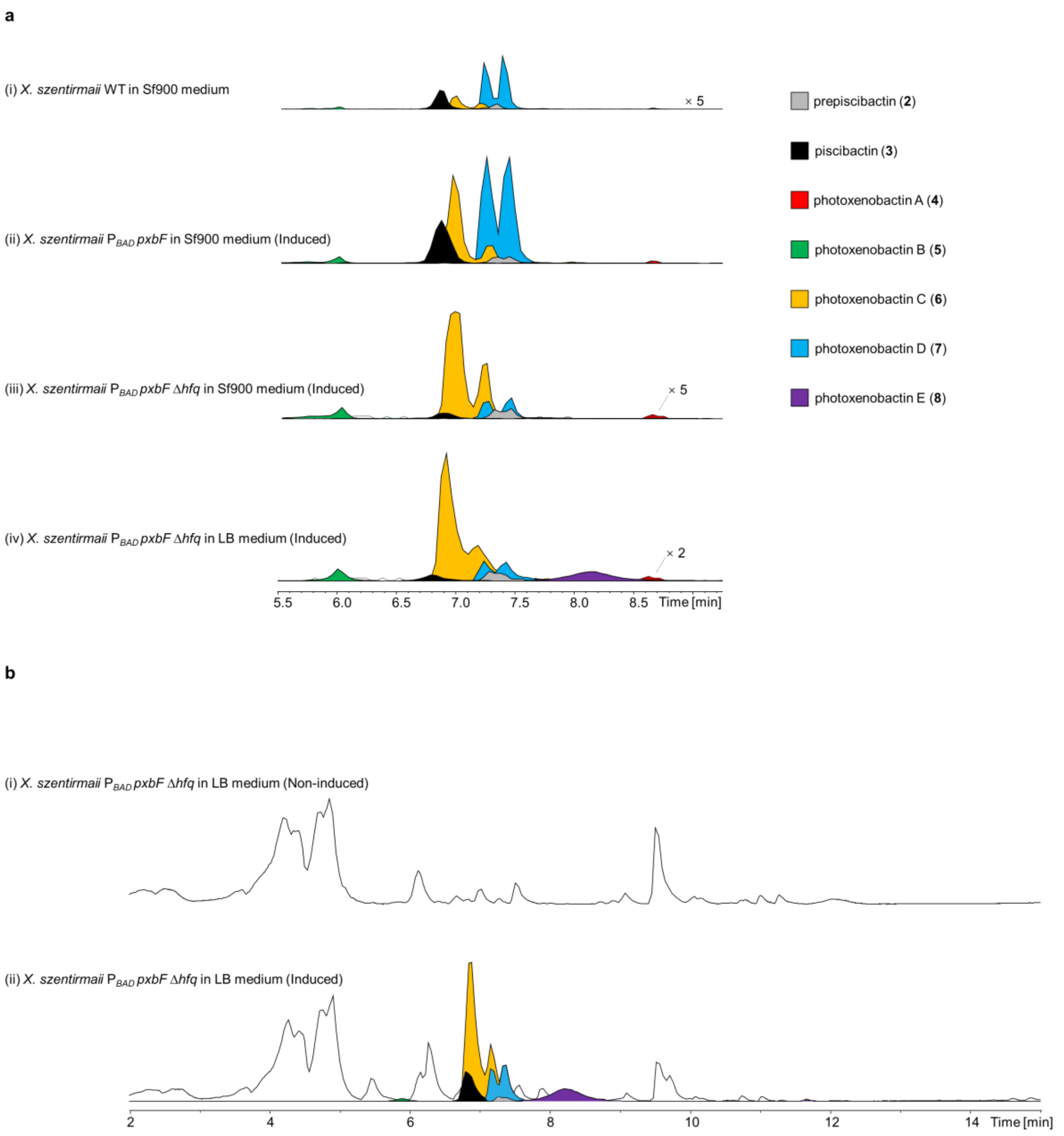

**Supplementary Fig. 7 | HPLC-MS analysis of photoxenobactins and piscibactins in *X. szentirmaii* DSM 16338 wild-type strain and the promoter exchange mutants thereof in different media. a,** EICs of the (i) wild-type strain and (ii-iv) mutants with L-arabinose induction. Shown are (i-iv) prepiscibactin (**2**), piscibactin (**3**), and photoxenobactins A (**4**), B (**5**), C (**6**), D (**7**), and E (**8**). Each compound contains a pair of C-10 epimers, which were not differentiated. Intensities in traces (i), (iii), and (iv) are magnified for visualizing tiny peaks. Magnifications are indicated on the right side of traces or on the top of the peak. **b,** BPCs of promoter exchange of the Δ*hfq* mutant (i) without and (ii) with L-arabinose induction. Desired peaks are highlighted in (**b**) the BPC of trace ii. Photoxenobactin E (**8**) was produced in a detectable amount in the *X. szentirmaii* $P_{BAD}$ *pxbF* Δ*hfq* mutant. Representative data from three independent experiments are shown.

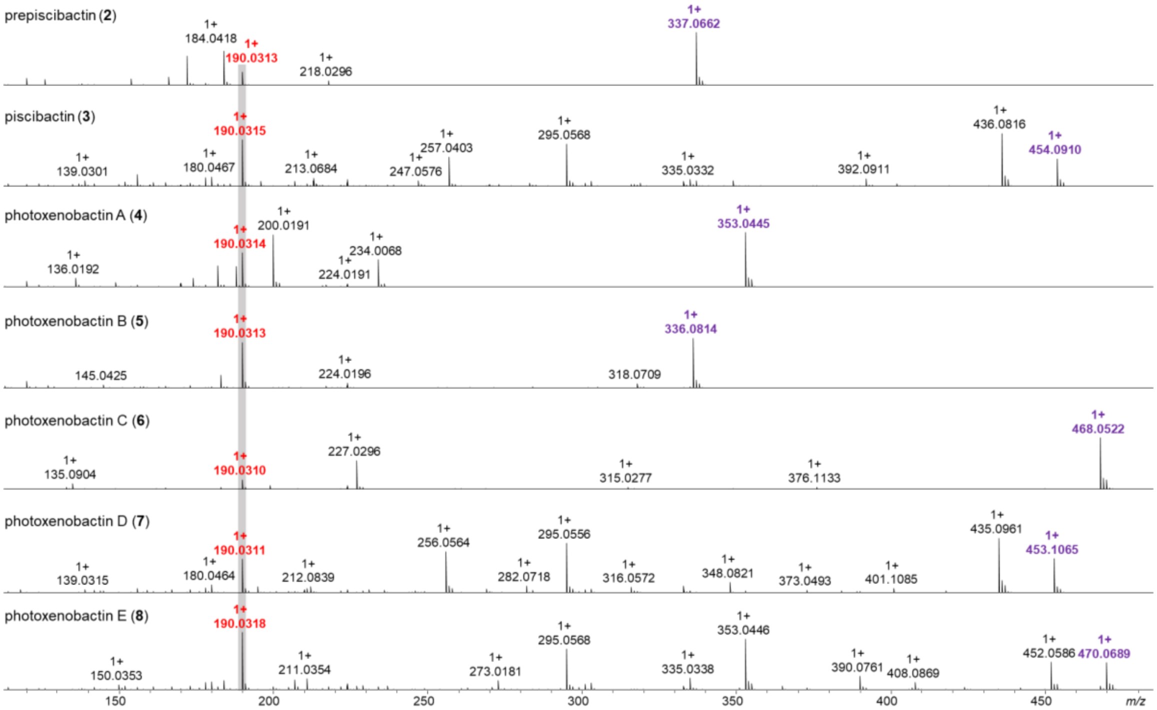

**Supplementary Fig. 8 | Comparison of MS/MS fragmentation patterns of prepiscibactin (2), piscibactin (3), and photoxenobactins A–E (4–8).** A diagnostic *m/z* 190.031 $[M + H]^+$ fragment ion indicating a hydroxyphenylthiazoline moiety [33] is highlighted in red. Purple masses indicate parent ions.

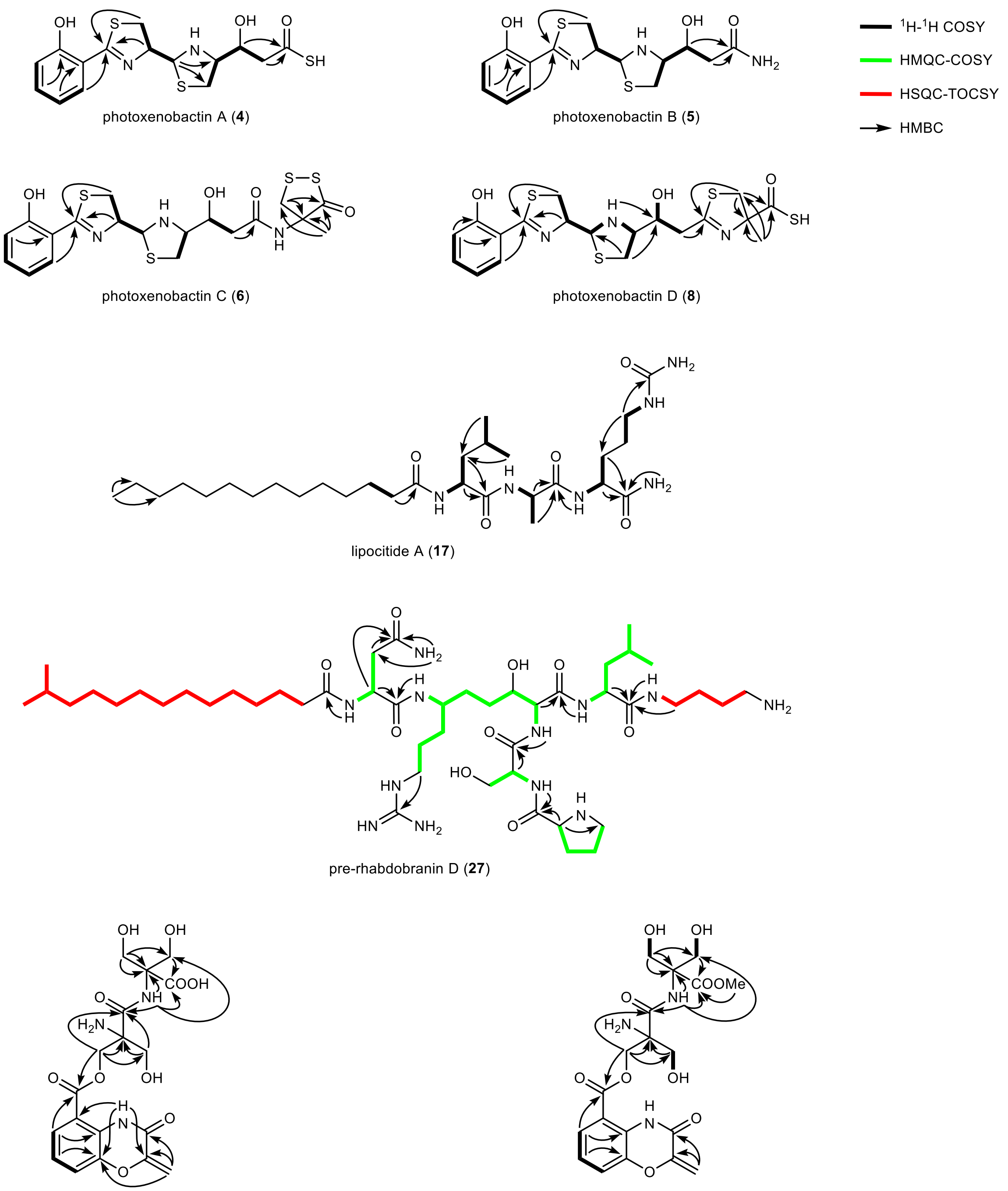

**Supplementary Fig. 9 | 2D NMR correlations of new natural products produced by *XP* strains.**

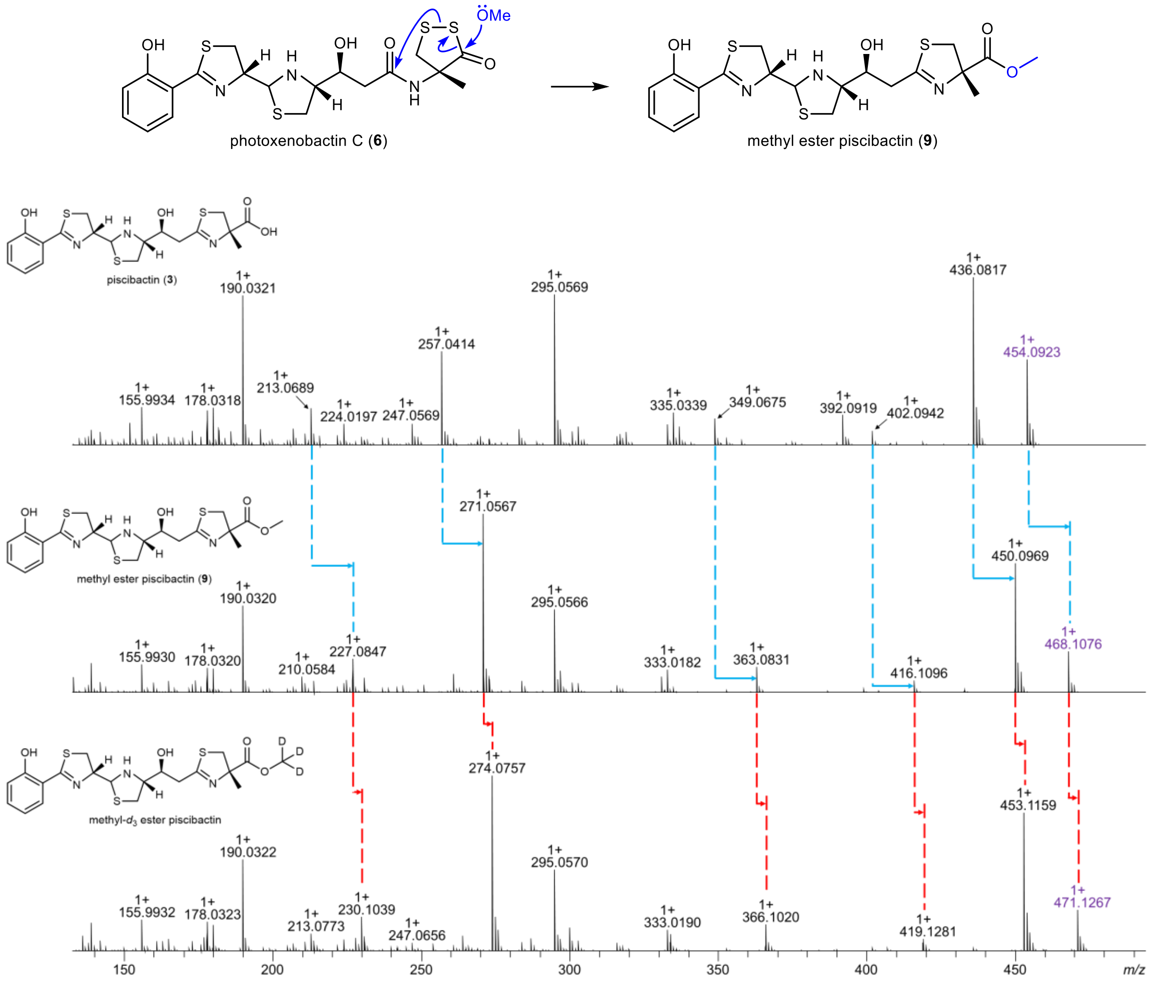

**Supplementary Fig. 10 | Proposed conversion of photoxenobactin C (6) in methanol.** The conversion of photoxenobactin C (**6**) into piscibactin (**3**) in methanol is dramatically accelerated under heating and UV light. Fragmentation patterns with a 14-Da shift between piscibactin (**3**) and methyl ester piscibactin (**9**) are shown with blue arrows, indicating a difference in the methyl group. Fragmentation patterns with a 3-Da shift between methyl ester piscibactin (**9**) and methyl-$d_3$ ester piscibactin that was obtained by incubation of photoxenobactin C (**6**) in methanol-$d_4$ are shown with red arrows, indicating a difference in the $d_3$-methyl group.

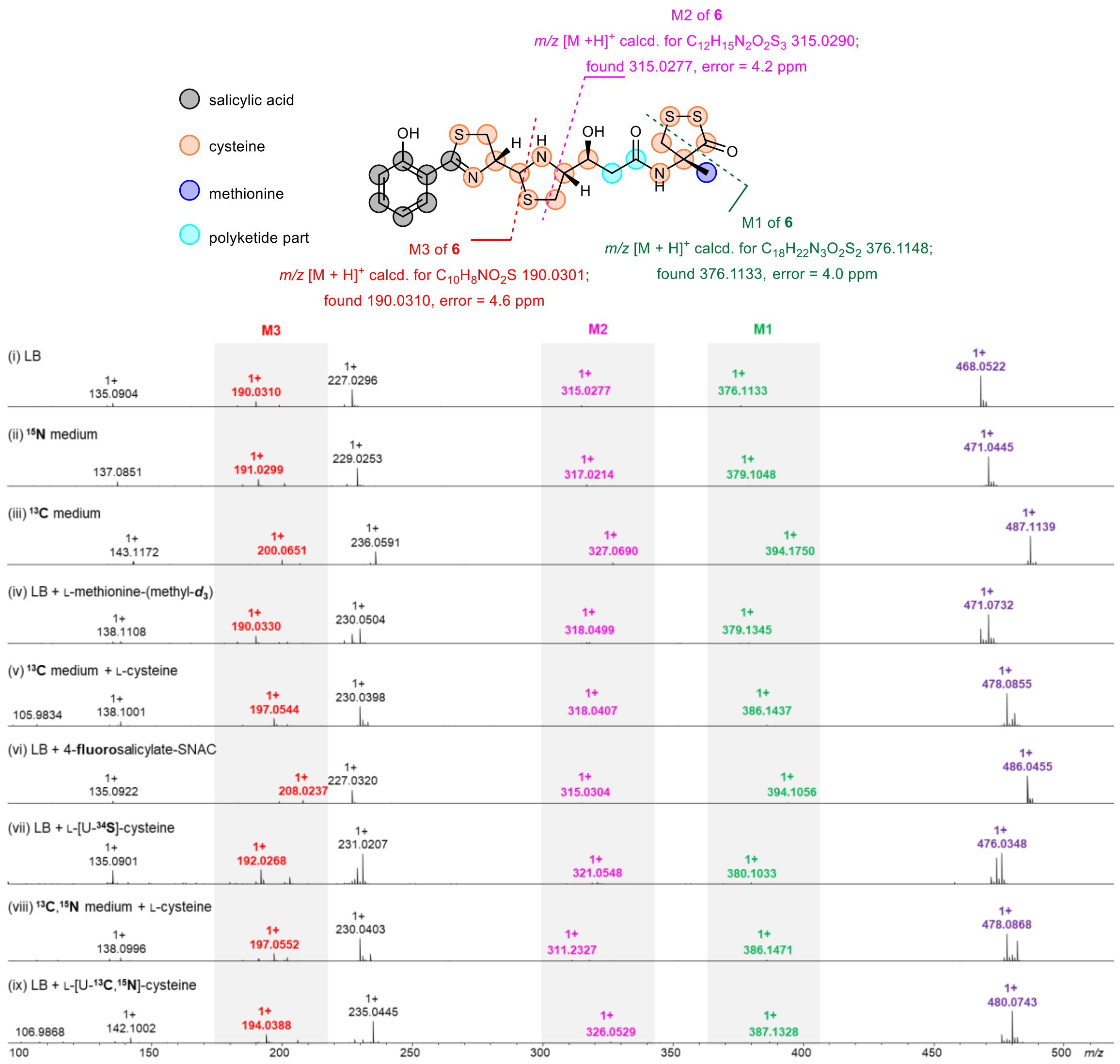

**Supplementary Fig. 11 | Mass spectrometry fragmentation patterns of photoxenobactin C (6) resulting from (ii-ix) labeling experiments.** Positions incorporated with different building blocks that were supported by labels and/or $MS^2$ are shown as colored spheres. Purple masses indicate parent ions ($M - H_2O + H^+$) and fragment ions M1–3 are green, pink, and red.

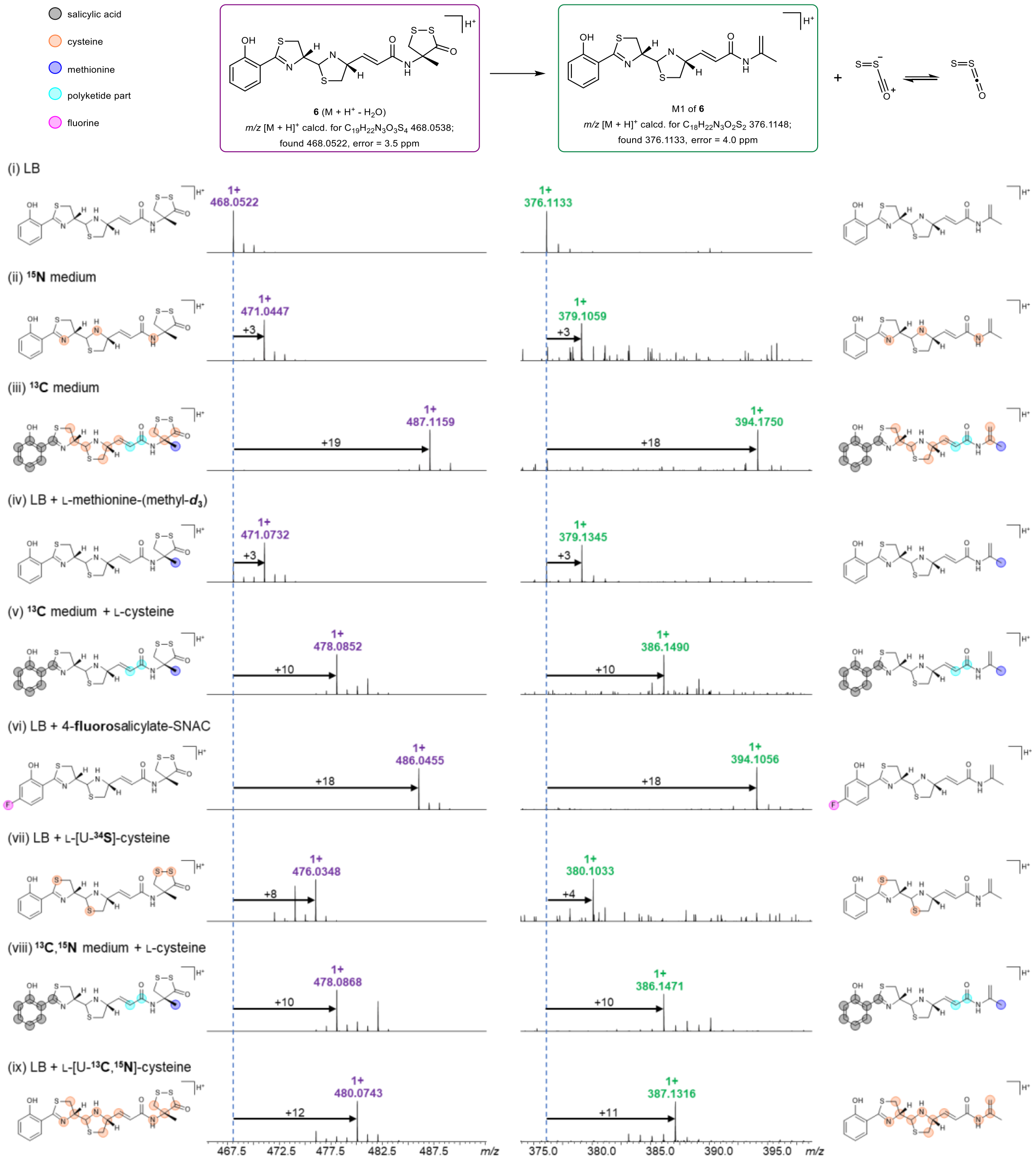

**Supplementary Fig. 12 | Mass spectrometry identification of photoxenobactin C (6) and a dithioperoxoate moiety thereof by (ii-ix) labeling experiments.** Positions shown as colored spheres are incorporated with labels in corresponding experiments. Purple and green masses/frames indicate parent ions (M – $H_2O$ + $H^+$) and fragment ions M1, respectively. Black arrows indicate mass shifts. The number of nitrogen and carbon atoms was confirmed by (ii) $^{15}N$ and (iii) $^{13}C$ labeling media. A mass shift of 3 Da in (iv) L-methionine-(methyl-$d_3$) feeding showed the incorporation of one *S*-adenosylmethionine derived methyl group. (vi) 4-Fluorosalicylate-SNAC supplement with a mass shift of 18 Da confirmed the incorporation of salicylate. (v and viii) Inverse feeding experiments with L-cysteine in $^{13}C$ & $^{13}C$,$^{15}N$ media background together with (ix) L-[U-$^{13}C$,$^{15}N$]-cysteine feeding confirmed three cysteine building blocks being part of photoxenobactin C (**6**). However, four sulfur atoms were found to be incorporated by (vii) L-[U-$^{34}S$]-cysteine feeding. Comparison of the parent masses and fragment ions M1 suggested the C terminus lost a $COS_2$ moiety which contains a carbon atom as observed in (iii) and (ix) as well as two sulfur atoms as in (vii).

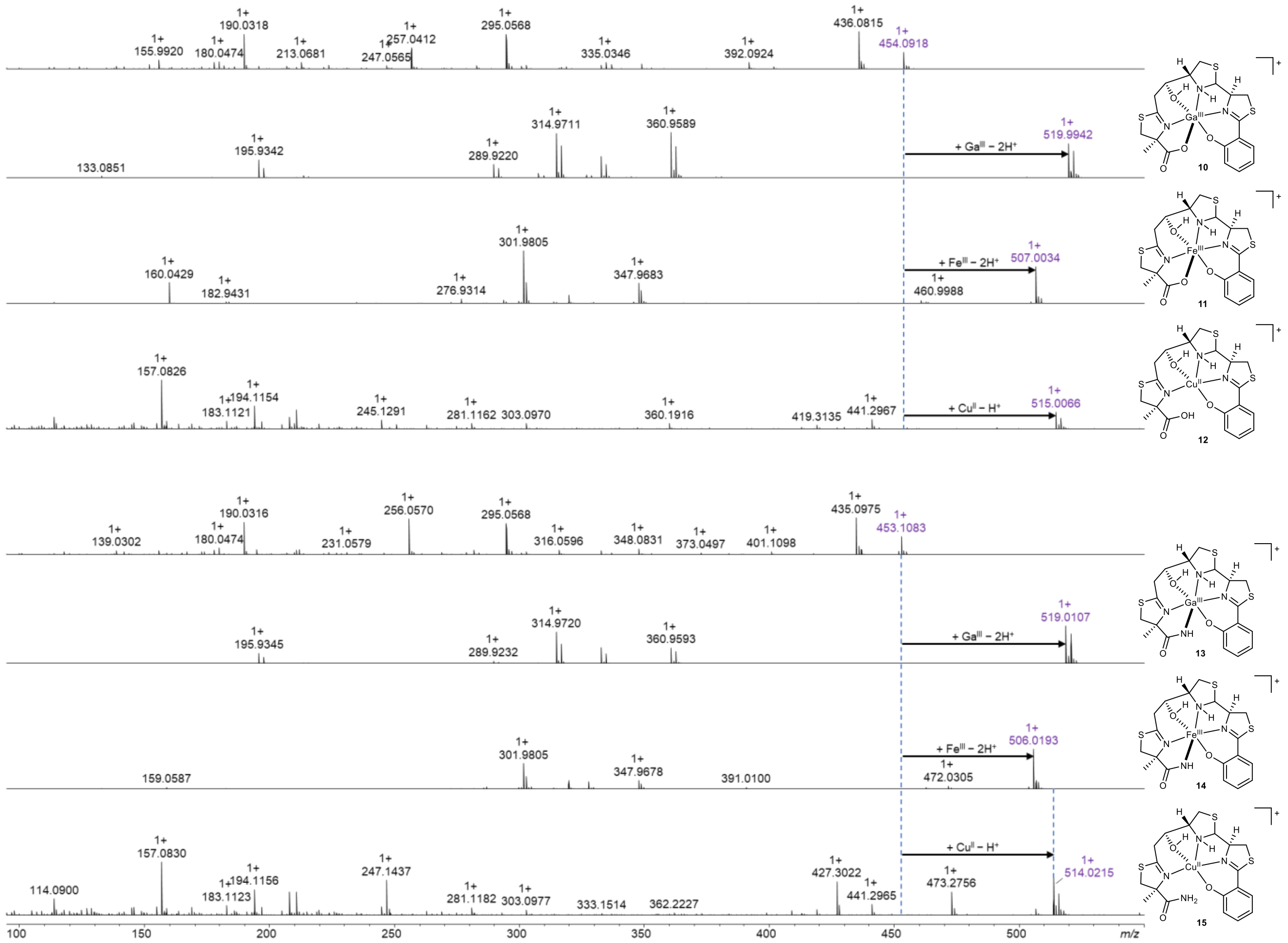

**Supplementary Fig. 13 | Metal chelating properties of piscibactin (10–12) and photoxenobactin D (13–15).** Ethyl acetate extracts of *X. szentirmaii* $P_{BAD}$ *pxbF* (induced) were incubated with $Fe(NO_3)_3$, $Ga(NO_3)_3$, and $CuCl_2$. Both piscibactin (**3**) and photoxenobactin D (**7**) can chelate $Fe^{III}$, $Ga^{III}$, and $Cu^{II}$. Purple indicates parent ions ($M + H^+$ or $M^+$). Black arrows indicate mass shifts.

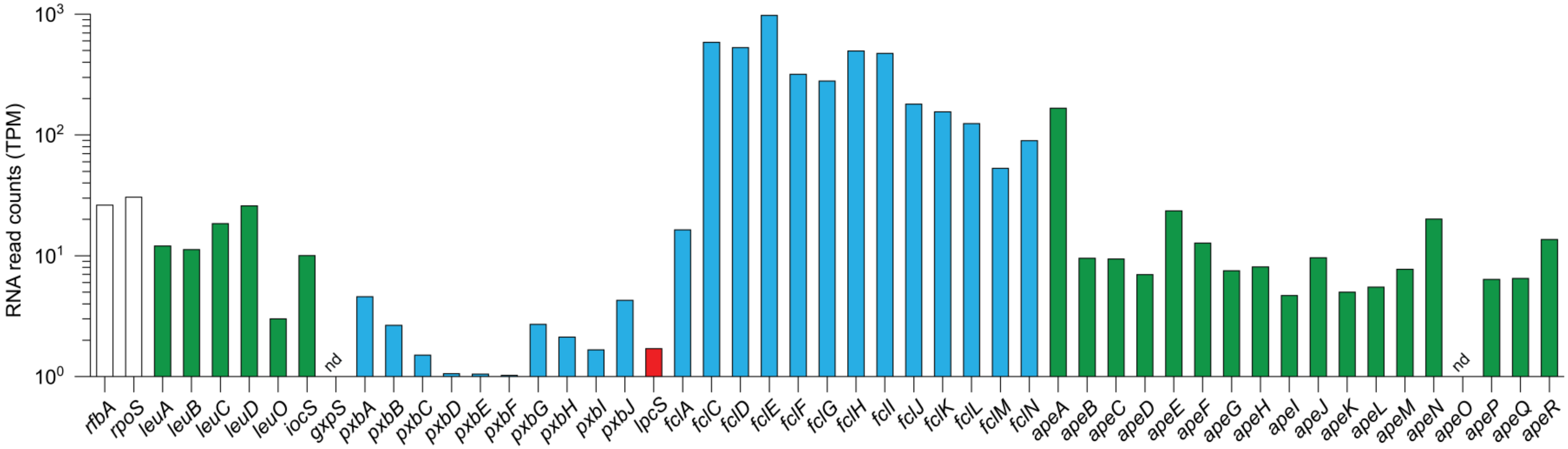

**Supplementary Fig. 14 | Comparison of transcriptional levels of biosynthetic genes in the conserved BGCs (*ioc/leu, gxp, pxb*, *lpc, fcl*, and *ape*) in *X. szentirmaii* US with the housekeeping genes (*rfbA* and *rpoS*).** See Supplementary Table 13 for actual values.

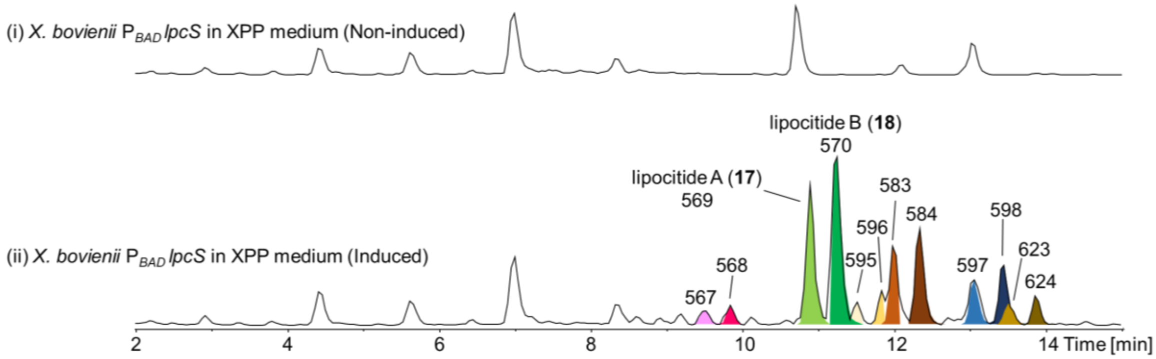

**Supplementary Fig. 15 | HPLC-MS analysis of lipocitides in the *X. bovienii* SS-2004 P$_{BAD}$ *lpcS* mutant in XPP medium.** BPCs of the promoter exchange mutant (i) without and (ii) with L-arabinose induction. Lipocitides are highlighted with colors with corresponding $[M + H]^+$ ions. Representative data from three independent experiments are shown.

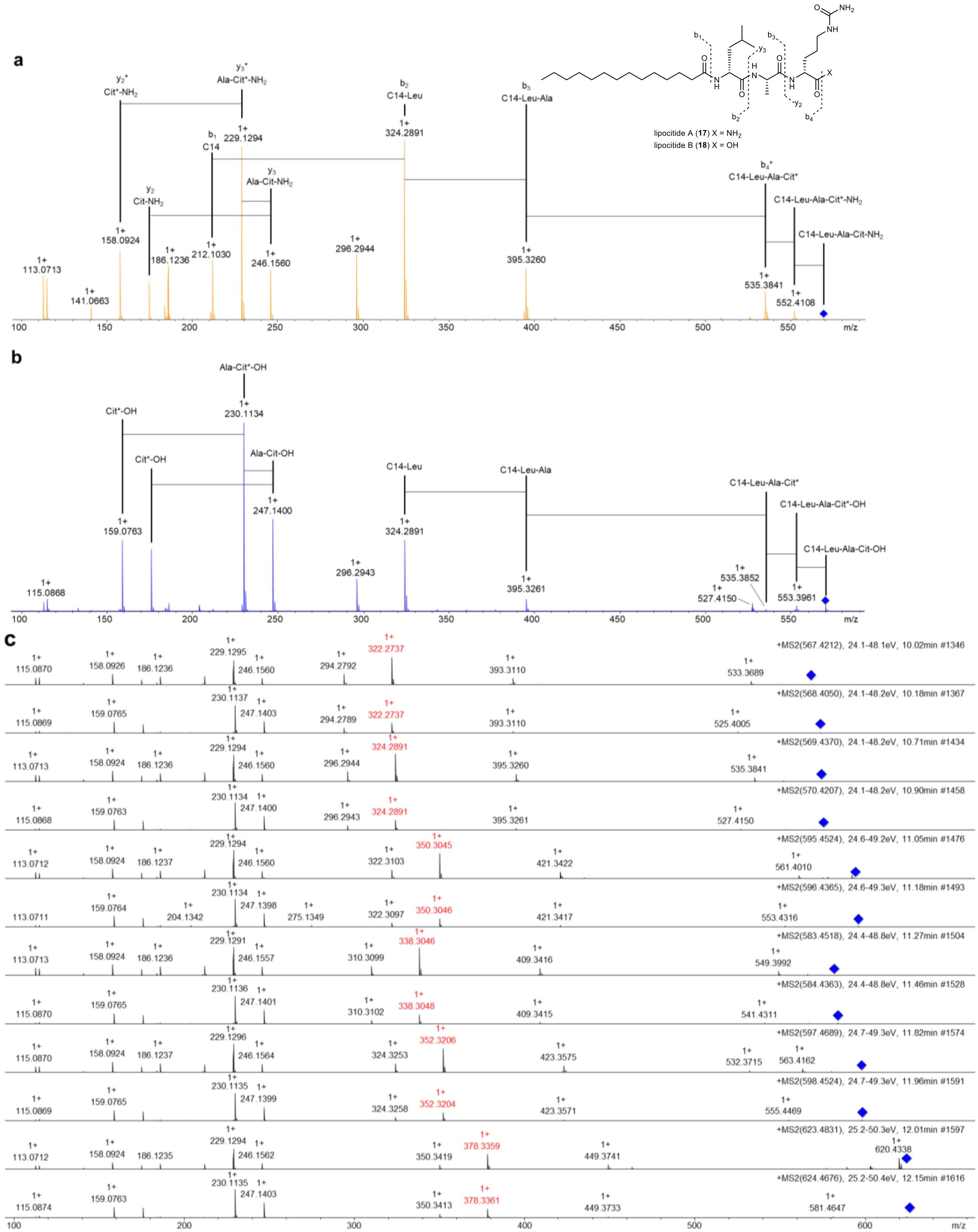

**Supplementary Fig. 16 | MS/MS fragmentation patterns of lipocitides. a,** Fragmentation patterns of lipocitide A (**17**). **b,** Fragmentation patterns of lipocitide B (**18**). The asterisk in a citrulline and fragment ion represents a loss of $NH_2$ in the ureido group. **c,** Comparison of MS/MS fragmentation patterns between lipocitides. Fragment ions highlighted in red represent an acyl/Leu pattern, indicating that lipocitides differ in N-acyl groups. The blue diamond indicates the parent ions.

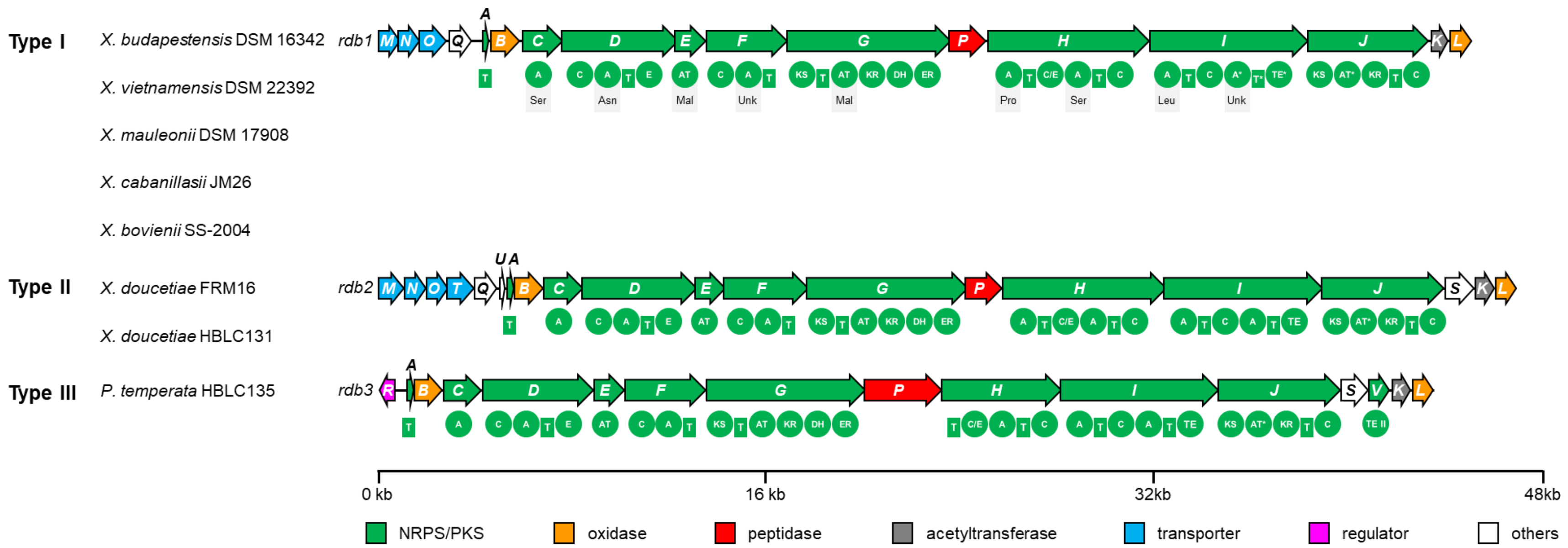

**Supplementary Fig. 17 | Domain organization of three types of *rdb* BGCs with predicted substrates of adenylation and acyltransferase domains.** T, thiolation; A, adenylation; C, condensation; E, epimerization; AT, acyltransferase; KS, ketosynthase; KR, ketoreductase; DH, dehydratase; ER, enoyl reductase; cMT, carbon methyltransferase; TE, thioesterase; Ser, serine; Asn, asparagine; Mal, malonyl; Pro, proline; Leu, leucine; Unk, unknown. Presumably inactive domain is labeled with an asterisk.

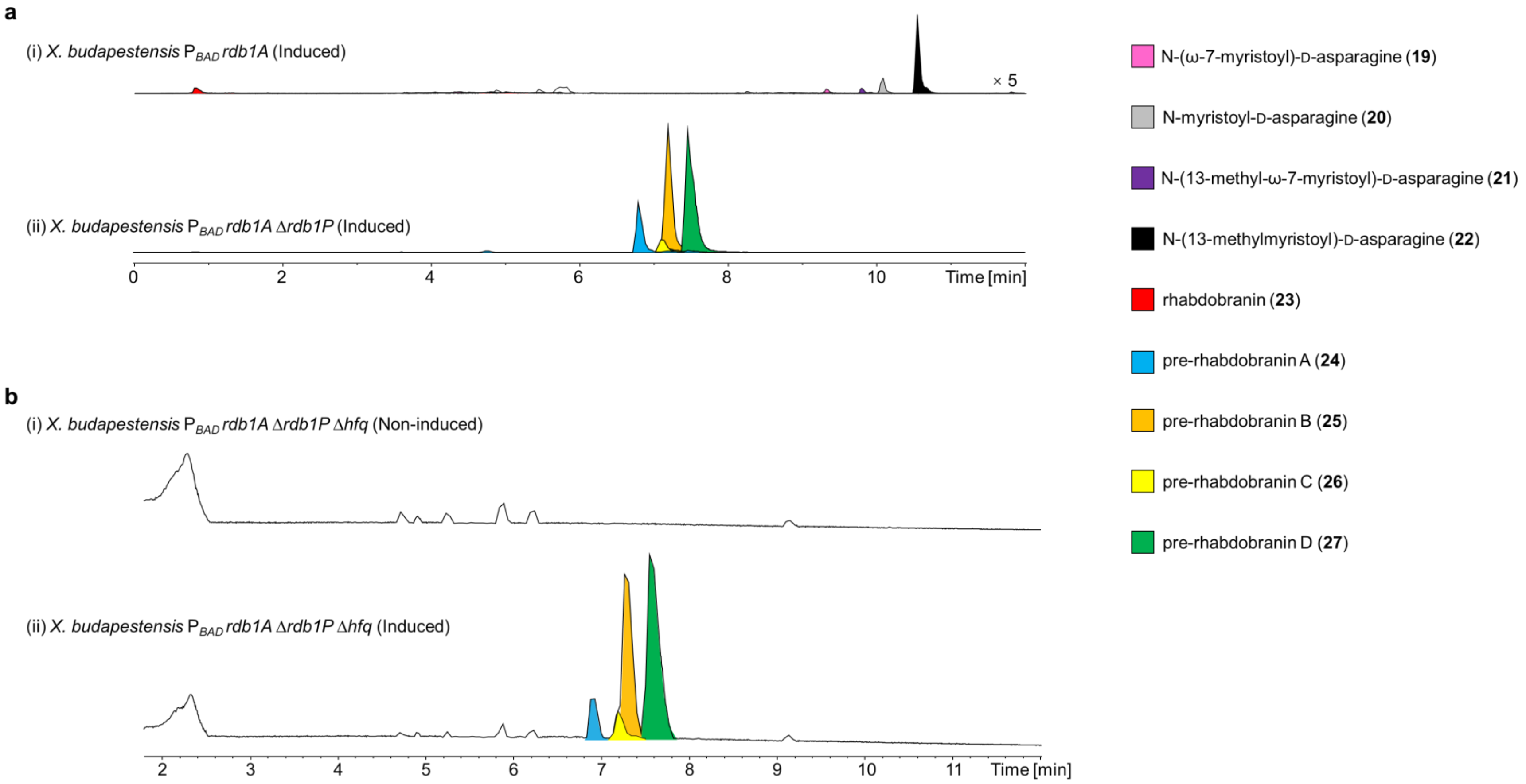

**Supplementary Fig. 18 | HPLC-MS analysis of (pre-)rhabdobranins in the promoter exchange mutants of *X. budapestensis* DSM 16342 in LB medium. a,** EICs of the *X. budapestensis* $P_{BAD}$ *rdb1A* and *X. budapestensis* $P_{BAD}$ *rdb1A Δrdb1P* mutants. Shown are (i) N-(ω-7-myristoyl)-D-asparagine (**19**), N-myristoyl-D-asparagine (**20**), N-(13-methyl-ω-7-myristoyl)-D-asparagine (**21**), N-(13-methylmyristoyl)-D-asparagine (**22**), and rhabdobranin (**23**); (ii) pre-rhabdobranin A (**24**), pre-rhabdobranin B (**25**), pre-rhabdobranin C (**26**), and pre-rhabdobranin D (**27**). Intensities in trace (i) are magnified for visualizing tiny peaks. Magnifications are indicated on the right side of traces. **b,** BPCs of the (i) non-induced and (ii) induced promoter exchange mutants of the *X. budapestensis* $P_{BAD}$ *rdb1A Δrdb1P Δhfq*. Desired peaks are highlighted in trace (ii). Mutants were induced with L-arabinose. Representative data from three independent experiments are shown.

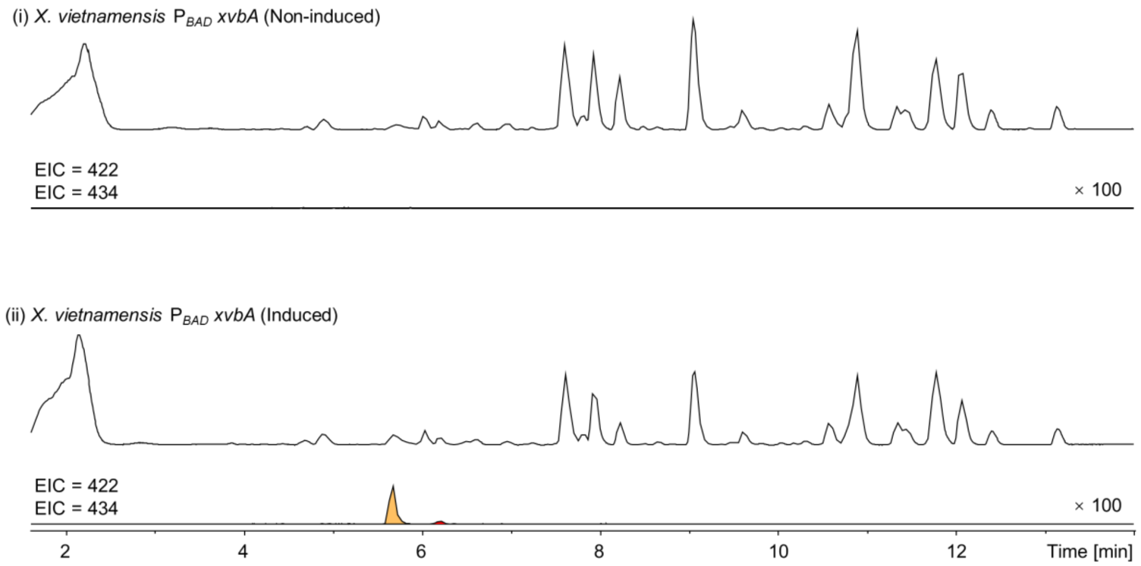

**Supplementary Fig. 19 | HPLC-MS analysis of benzobactins in the promoter exchange mutants of *X. vietnamensis* DSM 22392 in XPP medium.** (i) BPCs and EICs of the non-induced mutant. (ii) BPCs and EICs of the induced mutant. Desired peaks are highlighted in trace (ii). Benzobactins A (**28**, orange) and the methyl ester thereof (**29**, red) are highlighted in the EICs. Intensities in EIC traces (i) and (ii) are magnified for visualizing tiny peaks. Magnifications are indicated on the right side of traces. Mutants were induced with L-arabinose. Representative data from three independent experiments are shown.

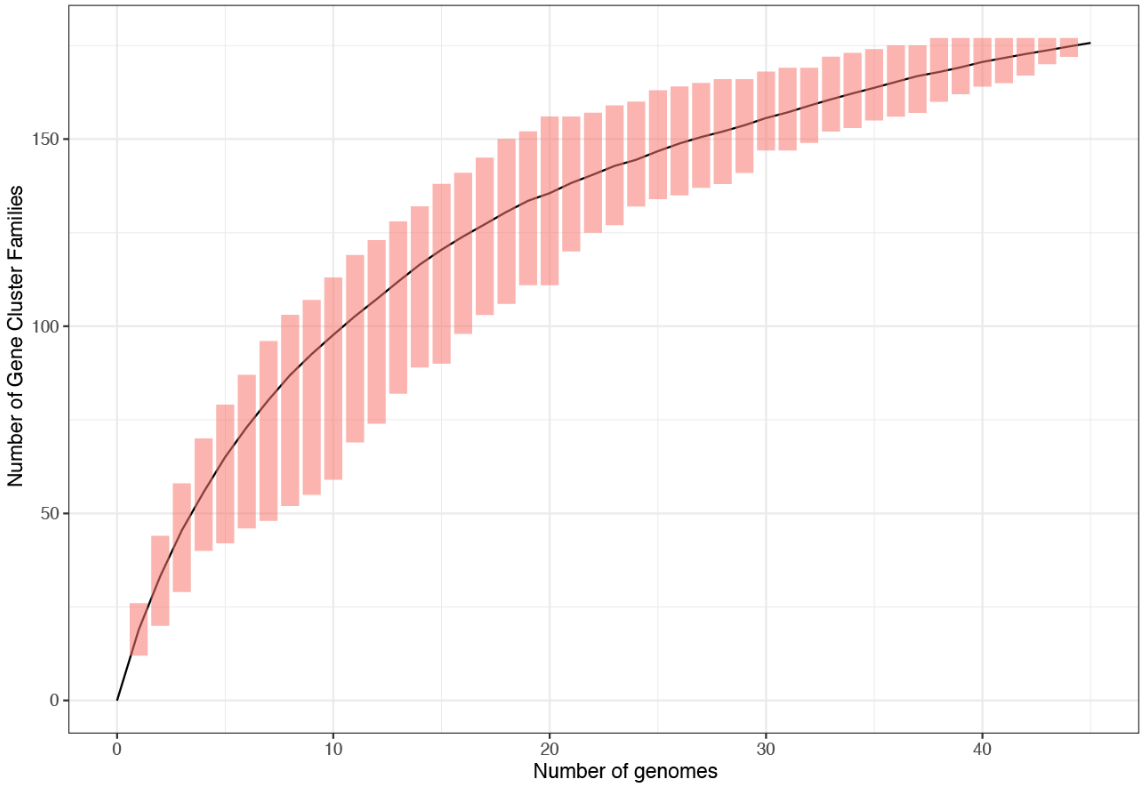

**Supplementary Fig. 20 | Rarefaction analysis for the 176 GCFs in 45 *XP* strains.** The line shows the mean accumulation of 100 permutations with bars indicating the maximum and minimum values over these random shuffles. A rarefaction analysis was computed in R using the rarefaction function as a part of the mircopan package with n.perm set to 100. GCFs are defined by BiG-SCAPE distance metrics with a raw distance cut-off of 0.65 and refined based on our in-house database.

**a**

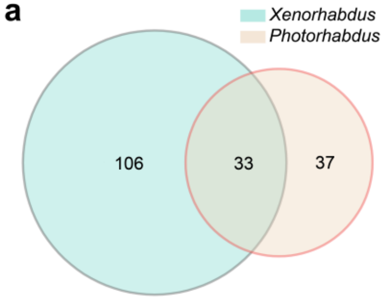

**b**

| BGCs/GCFs | *Xenorhabdus* | | *Photorhabdus* | |
|---|---|---|---|---|
| | sum | mean | sum | mean |
| BGCs | 604 | 20.8 | 396 | 24.8 |
| unknown BGCs | 340 | 11.7 | 195 | 12.2 |
| GCFs | 139 | 4.8 | 70 | 4.4 |
| genus-specific GCFs | 106 | 3.7 | 37 | 2.3 |
| unique genus-specific GCFs* | 40 | 1.4 | 16 | 1.0 |
| unique genus-specific isolated clade GCFs* | 17 | 0.6 | 8 | 0.5 |
| singletons | 23 | 0.8 | 8 | 0.5 |

**c**

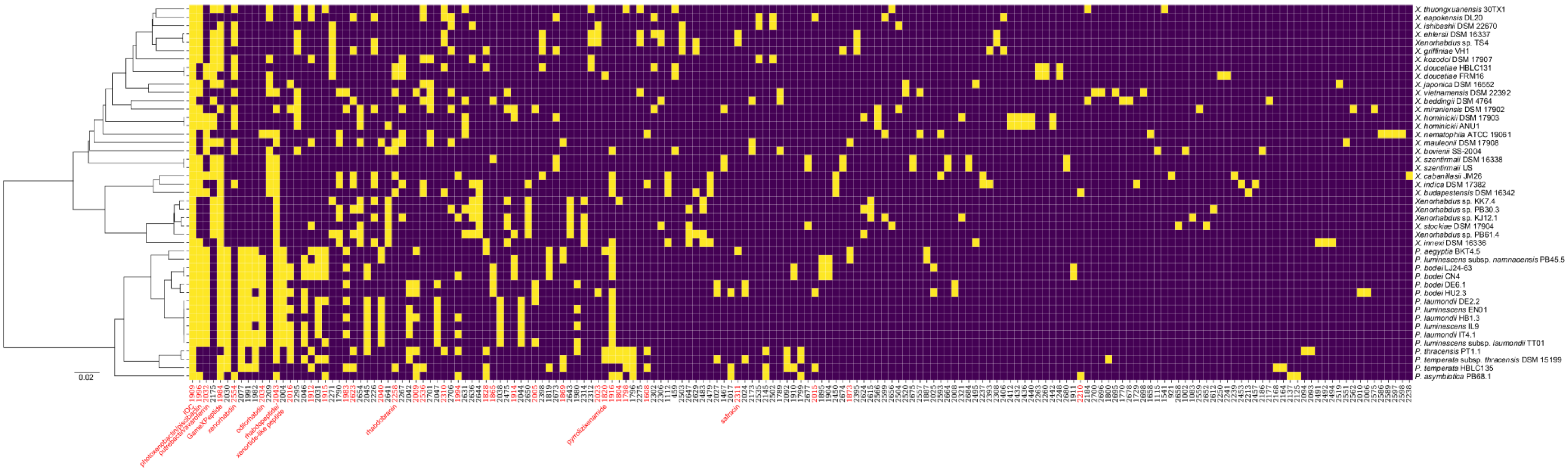

**Supplementary Fig. 21 | Distributions of GCFs on genus and species levels**. **a,** Venn diagram on the distribution of GCFs between *XP* genera. **b,** Distribution of BGCs and GCFs in *Xenorhabdus* and/or *Photorhabdus*. Asterisk indicates BGCs within a given GCF having no connections with MIBiG reference BGCs and the main BiG-SCAPE network. **c,** Phyletic distribution of GCFs in *XP*. A yellow square indicates the presence of a GCF in the respective species. GCFs shared by *XP* are highlighted in red and compound annotations are indicated for those that have been identified. GCFs are defined by BiG-SCAPE distance metrics with a raw distance cut-off of 0.65 and refined based on our in-house database.

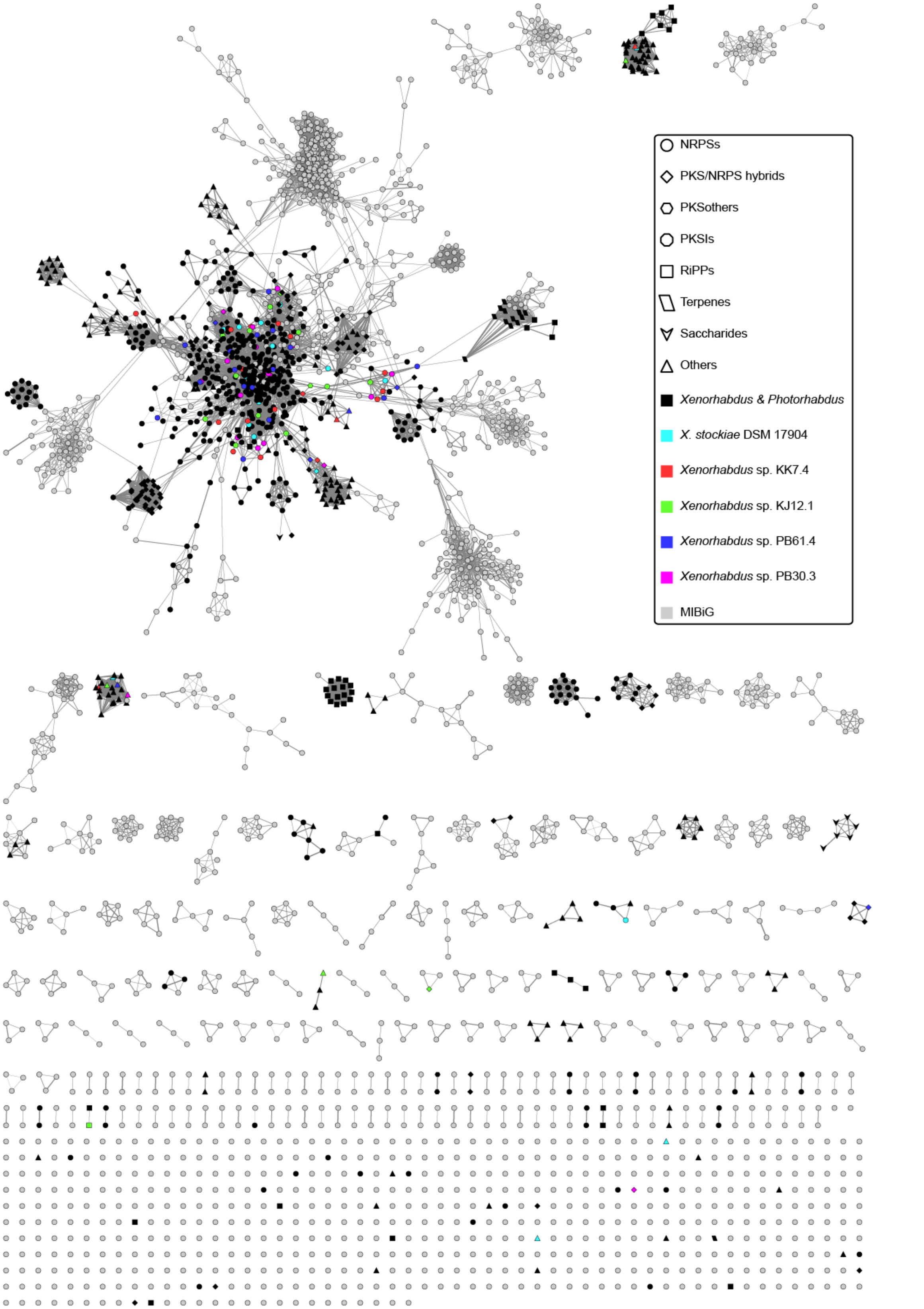

**Supplementary Fig. 22 | BGC distribution of *Xenorhabdus stockiae* DMS 17904, *Xenorhabdus* sp. KK7.4, *Xenorhabdus* sp. KJ12.1, *Xenorhabdus* sp. PB30.3, and *Xenorhabdus* sp. PB61.4 in the sequence similarity network.**
