## Supplementary Note for "Global analysis of biosynthetic gene clusters reveals conserved and unique natural products in entomopathogenic nematode-symbiotic bacteria"

### Synthetic procedures and spectra

### General synthetic procedures

The Fmoc protecting group was removed with 2 mL 40% piperidine/DMF (5 min) followed by 2mL 20% piperidine/DMF (10 min). Washings between coupling and deprotection steps were performed with DMF (5 syringe volumes) and DCM (5 syringe volumes). Resin loadings were determined by Fmoc-cleavage from a weighted resin sample [1]. The combined filtrates containing Fmoc cleavage products were quantified spectrophotometrically at 301 nm using a UV/Vis spectrophotometer with Hellma absorption cuvettes with a path length of 1 cm. Loadings were calculated in mmol resin using the Lambert-Beer's law with ε = 7800 $Mz^{-1}$ $cm^{-1}$: loading (mmol) = $\frac{Abs\,(sample)}{\varepsilon l} \times V$. Final cleavage was achieved by shaking the resin in a mixture of 2 mL TFA/TIPS/$H_2O$ (95:2.5:2.5) for 1 h. Then the filtrate was collected and the resin was washed three times (2 mL each) with DCM, and the combined filtrates were dried under reduced pressure.

### Syntheses of lipocitide A (17)

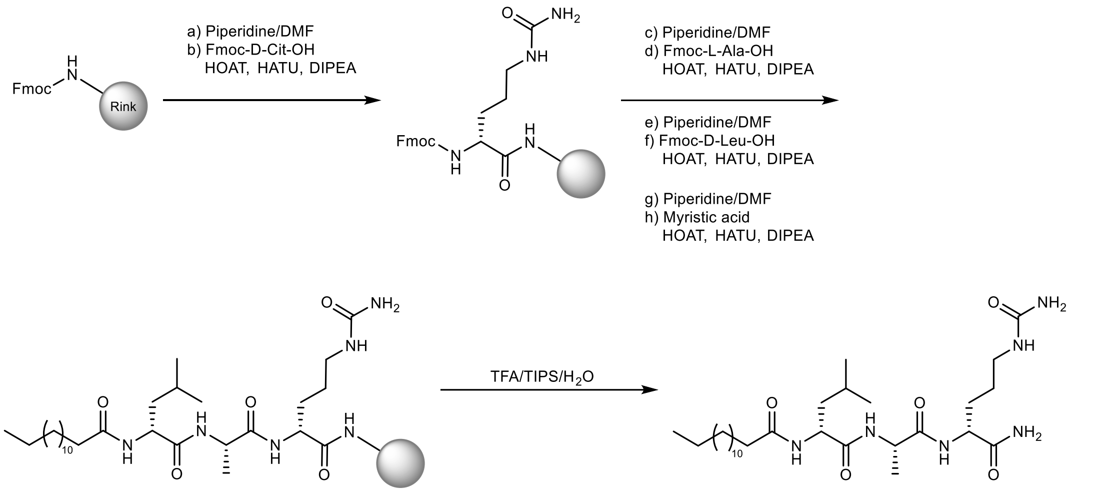


Fmoc-protected Rink Amide resin (192 mg, 0.52 mmol/g, 0.1 mmol) was placed in a polypropylene 6 mL syringe vessel fitted with polyethylene porous filter disks and swollen in 3 mL DMF for 10 min. Subsequently, the Fmoc-protected resin was deprotected and then washed as described in the general synthetic procedures. Fmoc-D-Cit-OH (198.0 mg, 0.5 mmol, 5 equiv.), HOAT (0.83 mL, 0.5 mmol, 5 equiv.), HATU (190.5 mg, 0.5 mmol, 5 equiv.), and DIPEA (170 µL, 1.0 mmol, 10 equiv.) were dissolved in 1.5 mL dry DMF. After 5 min the clear solution was added to the resin and shaken at room temperature overnight. The resin was washed and loading was calculated (79.2%) as described in the general synthetic procedures. Acylation of Fmoc-L-Ala-OH (74.1 mg, 0.24 mmol, 3 equiv.), Fmoc-D-Leu-OH (84.8 mg, 0.24 mmol, 3 equiv.), and myristic acid (54.8 mg, 0.24 mmol, 3 equiv.) were carried out using the abovementioned procedure. Final cleavage was performed as described in the general synthetic procedures, and the crude (70.8 mg) was purified by HPLC to obtain lipocitide A 24.3 mg (54.0 %) as a white solid.

### Syntheses of lipocitide B (18)

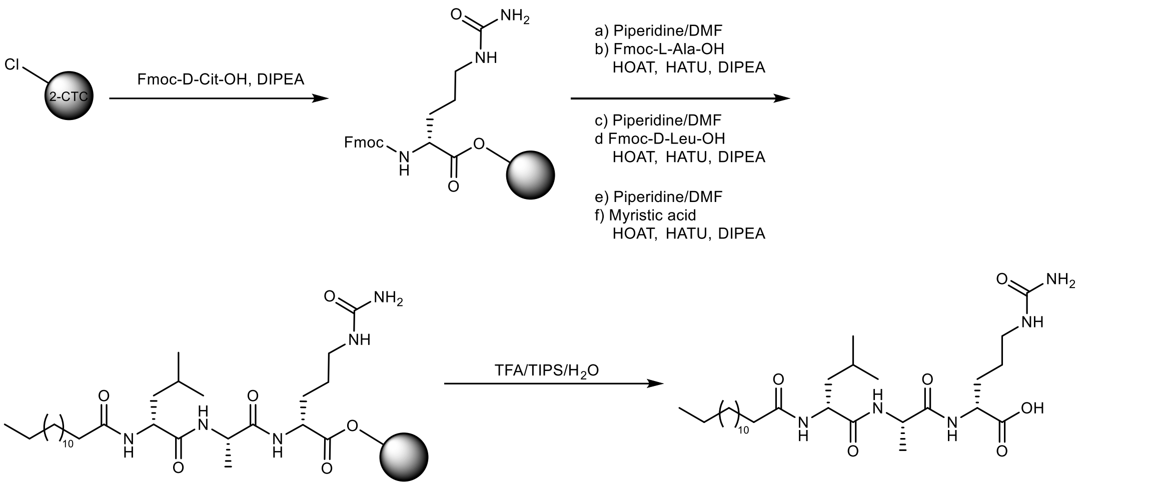


2-CTC resin (63 mg, 1.6 mmol/g, 0.1 mmol) was placed in a polypropylene 6 mL syringe vessel fitted with polyethylene porous filter disks. The resin was incubated with Fmoc-D-Cit-OH (119.0 mg, 0.3 mmol, 3 equiv.) and DIPEA (153 µL, 0.9 mmol, 9 equiv.) in 1.5 mL dry DCM at room temperature overnight. The resin was washed and loading was calculated (56.7%) as described in the general synthetic procedures. Acylation of Fmoc-L-Ala-OH (52.9 mg, 0.17 mmol, 3 equiv.), Fmoc-D-Leu-OH (60.1 mg, 0.17 mmol, 3 equiv.), and myristic acid (38.9 mg, 0.24 mmol, 3 equiv.) were performed with additional HOAT (0.47 mL, 0.28 mmol, 5 equiv.), HATU (108 mg, 0.28 mmol, 5 equiv.), and DIPEA (96 µL, 0.56 mmol, 10 equiv.). Final cleavage was carried out as described in the general synthetic procedures, and the crude (54.2 mg) was purified by HPLC to obtain lipocitide B 18.6 mg (57.6 %) as a white solid.

### Synthesis of *S*-(2-acetamidoethyl)4-fluoro-2-hydroxybenzothioate (4-Fluoro-salicylate-SNAC).

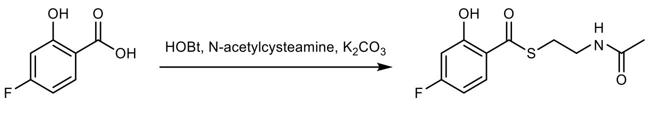


To a solution of 4-fluorosalicylic acid (156 mg, 1.0 mmol, 1.0 equiv.) and HOBt (162 mg, 1.2 mmol, 1.2 equiv.) in 45 mL THF, DCC (248 mg, 1.2 mmol, 1.2 equiv.) was added, followed by *N*-acetylcysteamine (112 µL, 1.0 mmol, 1.0 equiv.). After 1 h at room temperature, $K_2CO_3$ (138 mg, 1.0 mmol, 1.2 equiv.) was added and the reaction was stirred for an additional 2 h. The reaction mixture was then filtered and concentrated by rotary evaporation. The solid residue was dissolved in ethyl acetate and washed with sat. $NaHCO_3$ (50 mL) and water (50 mL). The organic layer was dried over $MgSO_4$, concentrated and purified by flash chromatography (1-10% MeOH in $CHCl_3$) to give 26 mg (10%) *S*-(2-acetamidoethyl)4-fluoro-2-hydroxybenzothioate.

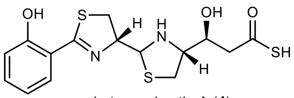

photoxenobactin A (**4**)

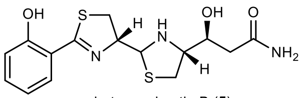

photoxenobactin B (**5**)

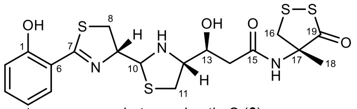

photoxenobactin C (**6**)

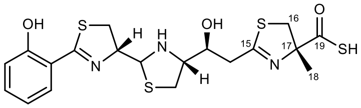

photoxenobactin E (**8**)

**Supplementary Note Table 1 |** [1]H and [13]C NMR data assignments for photoxenobactins A–C (**4**–**6**) and E (**8**) in DMSO-$d_6$ (for NMR spectra see Supplementary Note Figs. 4–24).

| | No. | Photoxenobactin A (**4**) | | Photoxenobactin B (**5**) | | Photoxenobactin C (**6**) | | Photoxenobactin E (**8**) | |
|---|---|---|---|---|---|---|---|---|---|
| | | $\delta_H$ (mult., *J*)[a] | $\delta_C$, mult.[f] | $\delta_H$ (mult., *J*)[b] | $\delta_C$, mult.[f] | $\delta_H$ (mult., *J*)[c] | $\delta_C$, mult.[f] | $\delta_H$ (mult., *J*)[d] | $\delta_C$, mult.[e] |
| SA | | | | | | | | | |
| | 1 | - | 158.9, C | - | 158.9, C | - | nd | - | 172.4, C |
| | 2 | 6.99 (br d, 7.7) | 117.1, CH | 6.98 (dd, 7.3, 1.2) | 117.2, CH | 6.97 (ov) | 119.7, CH | 6.50 (d, 8.5) | 124.4, CH |
| | 3 | 7.44 (ov) | 134.1, CH | 7.44 (ov) | 133.8, CH | 7.44 (ov) | 130.7, CH | 7.09 (td, 8.5, 1.8) | 133.9, CH |
| | 4 | 6.97 (td, 7.7, 1.1) | 119.5, CH | 6.95 (dd, 7.3, 1.2) | 119.6, CH | 6.97 (ov) | 117.4, CH | 6.32 (br t, 7.3) | 112.1, CH |
| | 5 | 7.46 (br d 7.7) | 130.7, CH | 7.42 (ov) | 130.8, CH | 7.44 (ov) | 134.0, CH | 7.16 (dd, 7.9, 1.8) | 132.3, CH |
| | 6 | - | 115.9, C | - | 116.5, C | - | 116.1, C | - | 116.7, C |
| | 7 | - | 173.8, C | - | 171.8, C | - | 173.3, C | - | 172.1, C |
| L-Cys-1 | | | | | | | | | |
| | 8 | 3.61 (dd, 11.5, 9.1) | 33.7, $CH_2$ | 3.39 (ov) | 31.1, $CH_2$ | 3.52 (ov) | 34.6, $CH_2$ | 3.45 (ov) | 33.7, $CH_2$ |
| | | 3.34 (ov) | | 3.29 (ov) | | 3.52 (ov) | | 3.09 (ov) | |
| | 9 | 5.41 (ddd, 9.1, 7.5, 5.6) | 78.3, CH | 6.36 (td, 8.8, 3.6) | 76.0, CH | 4.94 (m) | 81.5, CH | 4.40 (ddd, 13.1, 10.3, 7.5) | 78.7, CH |
| | 10 | 5.65 (br d, 5.6) | 64.3, CH | 5.36 (br d, 3.6) | 61.7, CH | 5.35 (d, 8.3) | 64.5, CH | 4.71 (dd, 10.3, 6.8) | 70.6, CH |
| | 10NH | nd | - | nd | - | nd | - | 5.31 (dd, 10.3, 6.9) | - |
| L-Cys-2 | | | | | | | | | |
| | 11 | 3.27 (ov) | 27.2, $CH_2$ | 3.01 (t, 10.4) | 26.3, $CH_2$ | 2.97 (dd, 10.4, 8.7) | 30.8, $CH_2$ | 3.45 (dd, 12.3, 7.6) | 37.5, $CH_2$ |
| | | 3.02 (dd, 10.3, 7.8) | | 2.65 (ov) | | 2.90 (dd, 10.4, 6.6) | | 3.03 (br t, 11.4) | |
| | 12 | 4.70 (td, 7.8, 4.5) | 74.9, CH | 4.35 (ddd, 10.4, 5.5, 3.5) | 75.1, CH | 4.23 (m) | 68.9, CH | 3.72 (dd, 18.1, 10.3) | 66.3, CH |
| | 13 | 4.43 (t, 4.9) | 67.2, CH | 4.09 (t, 3.9) | 65.5, CH | 4.46 (m) | 66.8, CH | 3.99 (br s) | 67.8, CH |
| | 13OH | nd | - | nd | - | nd | - | 7.52 (s) | - |
| Polyketide | | | | | | | | | |
| | 14 | 3.47 (ov) | 57.8, $CH_2$ | 3.08 (dd, 16.5, 4.5) | 48.0, $CH_2$ | 2.79 (16.4, 6.6) | 39.3, $CH_2$ | 3.08 (ov) | 40.0, $CH_2$ |
| | | 3.94 (d, 17.5) | | 2.60 (ov) | | 2.47 (ov) | | 2.88 (d, 17.0) | |
| | 15 | - | 198.4, C | - | 162.6, C | - | 162.8, C | - | 170.6, C |
| L-Cys-3 | | | | | | | | | |
| | 16 | - | - | - | - | 3.46 (ov) | 45.7, $CH_2$ | 3.59 (d, 11.5) | 39.4, $CH_2$ |
| | | | | | | 3.37 (ov) | | 3.15 (d, 11.5) | |
| | 17 | - | - | - | - | - | 68.4, C | - | 93.1, C |
| | 18 | - | - | - | - | 1.42 (s) | 22.8, $CH_3$ | 1.56 (s) | 25.6, $CH_3$ |
| | 19 | - | - | - | - | - | 208.3, C | - | 212.1, C |

Data were recorded at [a]700, [b]500, [c]600, [d]800, and [e]200 MHz, respectively. [f]Data were extracted from HSQC and HMBC spectra. nd = not detectable. The stereochemistry was predicted by analyzing the *pxb* BGC and comparison of chemical shifts with piscibactins and yersiniabactin.

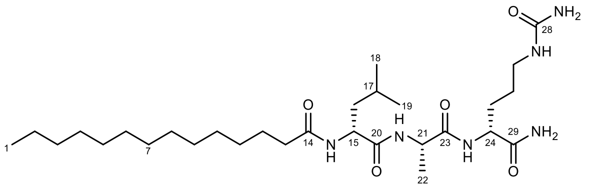


lipocitide A (**17**)

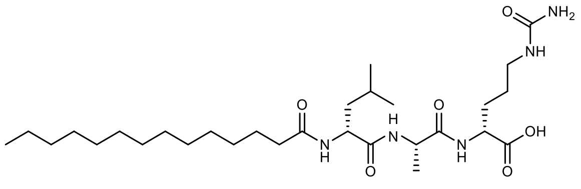


lipocitide B (**18**)

**Supplementary Note Table 2 |** $^{1}H$ (500 MHz) and $^{13}C$ (125 MHz) NMR data assignments for lipocitides A (**17**) and B (**18**) in DMSO-$d_6$ (for NMR spectra see Supplementary Note Figs. 28–32 and 35–39).

| | No. | Lipocitide A (**17**) | | Lipocitide B (**18**) | |
|---|---|---|---|---|---|
| | | $\delta_H$ (mult., *J*) | $\delta_C$, mult. | $\delta_H$ (mult., *J*) | $\delta_C$, mult. |
| FA | | | | | |
| | 1 | 0.85(ov) | 14.4, $CH_3$ | 0.85 (ov) | 14.4, $CH_3$ |
| | 2 | 1.24 (ov) | 22.6, $CH_2$ | 1.24 (ov) | 22.6, $CH_2$ |
| | 3 | 1.24 (ov) | 31.8, $CH_2$ | 1.24 (ov) | 31.8, $CH_2$ |
| | 4 | 1.24 (ov) | 29.2, $CH_2$ | 1.24 (ov) | 29.2, $CH_2$ |
| | 5 | 1.24 (ov) | 29.5, $CH_2$ | 1.24 (ov) | 29.5, $CH_2$ |
| | 6 | 1.24 (ov) | 29.5, $CH_2$ | 1.24 (ov) | 29.5, $CH_2$ |
| | 7 | 1.24 (ov) | 29.5, $CH_2$ | 1.24 (ov) | 29.5, $CH_2$ |
| | 8 | 1.24 (ov) | 29.5, $CH_2$ | 1.24 (ov) | 29.5, $CH_2$ |
| | 9 | 1.24 (ov) | 29.5, $CH_2$ | 1.24 (ov) | 29.5, $CH_2$ |
| | 10 | 1.24 (ov) | 29.2, $CH_2$ | 1.24 (ov) | 29.2, $CH_2$ |
| | 11 | 1.28 (ov) | 29.0, $CH_2$ | 1.28(m) | 29.0, $CH_2$ |
| | 12 | 1.48 (ov) | 25.7, $CH_2$ | 1.56 (ov) | 25.7, $CH_2$ |
| | 13 | 2.11 (m) | 35.5, $CH_2$ | 2.10 (m) | 35.6, $CH_2$ |
| | 14 | - | 173.1, C | - | 173.0, C |
| D-Leu | | | | | |
| | 15NH | 8.00 (t, 7.9) | - | 8.06 (br s) | - |
| | 15 | 4.24 (ov) | 51.9, CH | 4.26 (ov) | 51.8, CH |
| | 16 | 1.48 (ov) | 40.9, $CH_2$ | 1.41 (m) | 40.9, $CH_2$ |
| | 17 | 1.48 (ov) | 24.7, CH | 1.56 (ov) | 24.7, CH |
| | 18 | 0.84 (d, 6.4) | 23.4, $CH_3$ | 0.83 (d, 6.6) | 23.4, $CH_3$ |
| | 19 | 0.89 (d, 6.4) | 22.0 $CH_3$ | 0.88 (d, 6.6) | 22.0, $CH_3$ |
| | 20 | - | 172.6, C | - | 172.5, C |
| L-Ala | | | | | |
| | 21NH | 8.13 (t, 6.9) | - | 8.18 (br s) | - |
| | 21 | 4.24 (ov) | 48.7, CH | 4.26 (ov) | 48.6, CH |
| | 22 | 1.20 (d, 7.1) | 18.6, $CH_3$ | 1.20 (d, 7.1) | 19.0, $CH_3$ |
| | 23 | - | 172.4, C | - | 172.3, C |
| D-Cit | | | | | |
| | 24NH | 7.90 (dd, 8.0, 5.7) | - | 7.90 (br d, 8.2) | - |
| | 24 | 4.14(m) | 52.6, CH | 4.12 (m) | 52.5, CH |
| | 25a | 1.68 (m) | 29.7, $CH_2$ | 1.71 (m) | 29.5, $CH_2$ |
| | 25b | 1.58 (m) | | 1.56 (ov) | |
| | 26 | 1.30 (ov) | 27.1, $CH_2$ | 1.33 (m) | 26.9, $CH_2$ |
| | 27NH | 5.92 (t, 5.2) | - | 5.91 (t, 5.0) | - |
| | 27 | 2.91 (m) | 39.1, $CH_2$ | 2.91 (dd, 12.4, 6.2) | 39.3, $CH_2$ |
| | 28 | - | 159.2, C | - | 159.2, C |
| | 28$NH_2$ | 5.37 (s)<br>5.36 (s) | - | 5.36 (s) | - |
| | 29 | - | 174.1, C | - | 174.1, C |
| | 29$NH_2$ | 7.28 (d, 5.0)<br>7.02 (br s) | - | - | - |

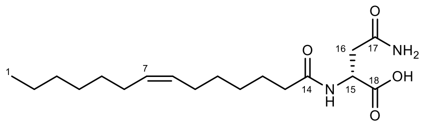

N-(ω-7-myristoyl)-D-asparagine (**19**)

**Supplementary Note Table 3 |** $^{1}H$ (500 MHz) and $^{13}C$ (125 MHz) NMR data assignments for N-(ω-7-myristoyl)-D-asparagine (**19**) in DMSO-$d_6$ (for NMR spectra see Supplementary Note Figs. 41–45).

| | No. | $\delta_H$ (mult., $J$) | $\delta_C$, mult. |
|---|---|---|---|
| FA[a] | | | |
| | 1 | 0.86 (t, 6.8) | 14.4, $CH_3$ |
| | 2 | 1.26 (ov) | 22.5, $CH_2$ |
| | 3 | 1.26 (ov) | 31.6, $CH_2$ |
| | 4 | 1.26 (ov) | 28.7, $CH_2$ |
| | 5 | 1.26 (ov) | 29.5, $CH_2$ |
| | 6 | 1.97 (ov) | 27.0, $CH_2$ |
| | 7 | 5.33 (ov) | 130.1, CH |
| | 8 | 5.33 (ov) | 130.1, CH |
| | 9 | 1.97 (ov) | 27.1, $CH_2$ |
| | 10 | 1.26 (ov) | 29.6, $CH_2$ |
| | 11 | 1.26 (ov) | 28.7, $CH_2$ |
| | 12 | 1.47 (ov) | 25.6, $CH_2$ |
| | 13 | 2.08 (t, 7.4) | 35.7, $CH_2$ |
| | 14 | - | 172.3, C |
| D-Asn | | | |
| | 15 | 4.35 (br s) | 40.9, CH |
| | 15NH | 7.79 (br s) | - |
| | 16 | 2.38 (m) | 38.3, $CH_2$ |
| | 17 | - | 172.3, C |
| | 17$NH_2$ | 7.54 (br s)<br>6.81 (br s) | - |
| | 18 | - | 172.2, C |

[a]The geometry of C-7/C-8 double bond was determined to be *cis* by comparing its chemical shifts with a known compound[2] that is a moiety of colibactin.

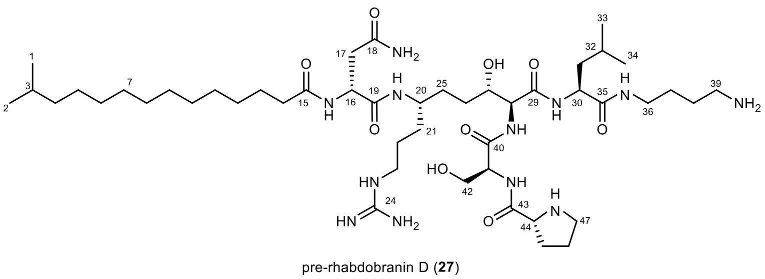

pre-rhabdobranin D (**27**)

**Supplementary Note Table 4 |** $^{1}H$ (700 MHz) and $^{13}C$ (175 MHz) NMR data assignments for pre-rhabdobranin D (**27**) in DMSO-$d_6$ (for NMR spectra see Supplementary Note Figs. 54–60)[a].

| | No. | $\delta_H$ (mult., *J*) | $\delta_C$, mult. |
|---|---|---|---|
| FA | | | |
| | 1,2 | 0.86 (d, 6.6) | 23.0, $CH_3$ |
| | 3 | 1.49 (ov) | 27.9, CH |
| | 4 | 1.13 (br dd, 14.2, 6,7) | 39.0, $CH_2$ |
| | 5 | 1.23 (ov) | 27.3, $CH_2$ |
| | 6 | 1.23 (ov) | 29.8, $CH_2$ |
| | 7 | 1.23 (ov) | 29.6, $CH_2$ |
| | 8 | 1.23 (ov) | 29.6, $CH_2$ |
| | 9 | 1.23 (ov) | 29.6, $CH_2$ |
| | 10 | 1.23 (ov) | 29.5, $CH_2$ |
| | 11 | 1.23 (ov) | 29.4, $CH_2$ |
| | 12 | 1.23 (ov) | 29.2, $CH_2$ |
| | 13 | 1.45 (ov) | 25.7, $CH_2$ |
| | 14 | 2.09 (t, 7.5) | 35.7, $CH_2$ |
| | 15 | - | 172.8, C |
| D-Asn | | | |
| | 16NH | 8.06 (d, 7.8) | - |
| | 16 | 4.49 (dd, 14.1, 7.8) | 50.5, CH |
| | 17 | 2.48 (ov) | 38.0, $CH_2$ |
| | | 2.38 (dd, 15.0, 7.8) | |
| | 18 | - | 172.2, C |
| | 18$NH_2$ | 7.40 (s) | - |
| | | 6.91 (s) | |
| | 19 | - | 171.5, C |
| L-Arg/polyketide | | | |
| | 20NH | 7.63 (d, 8.7) | - |
| | 20 | 3.66 (m) | 47.9, CH |
| | 21 | 1.43 (ov) | 31.9, $CH_2$ |
| | | 1.30 (m) | |
| | 22 | 1.45 (ov) | 20.4, $CH_2$ |
| | 23 | 3.02 (ov) | 40.9, $CH_2$ |
| | 24 | - | 157.8, C |
| | 25 | 1.44 (ov) | 30.9, $CH_2$ |
| | 26 | 1.42 (ov) | 29.8, $CH_2$ |
| | | 1.36 (m) | |
| | 27 | 3.62 (m) | 70.7, CH |
| | 28 | 4.15 (m) | 58.7, CH |
| | 28NH | 8.19 (d, 7.0) | - |
| | 29 | - | 170.8, C |
| L-Leu | | | |
| | 30NH | 8.15 (d, 7.4) | - |
| | 30 | 4.12 (m) | 52.0, CH |
| | 31 | 1.48 (ov) | 40.5, $CH_2$ |
| | 32 | 1.61 (ov) | 24.6, CH |
| | 33 | 0.86 (d, 6.6) | 23.6, $CH_3$ |
| | 34 | 0.79 (d, 6.6) | 21.5, $CH_3$ |
| | 35 | - | 172.4, C |
| putrescine | | | |
| | 36NH | 7.69 (t, 5.5) | - |
| | 36 | 3.02 (ov) | 38.4, $CH_2$ |
| | 37 | 1.57 (ov) | 26.1, $CH_2$ |
| | | 1.43 (ov) | |
| | 38 | 1.48 (ov) | 25.0, $CH_2$ |

| | | | |
|---|---|---|---|
| | 39 | 2.74 (t, 7.2) | 38.9, $CH_2$ |
| L-Ser | | | |
| | 40 | - | 171.1, C |
| | 41NH | 8.29 (d, 7.4) | - |
| | 41 | 4.36 (dd, 12.9, 5.6) | 55.0, CH |
| | 42 | 3.68 (dd, 10.8, 5.6) | 64.4, $CH_2$ |
| | | 3.51 (dd, 10.8, 5.6) | |
| | 43 | - | 174.8, C |
| D-Pro | | | |
| | 44 | 3.60 (m) | 60.4, CH |
| | 45 | 1.94 (ddd, 15.4, 12.4, 8.3) | 30.6, $CH_2$ |
| | | 1.69 (dt, 12.4, 6.8) | |
| | 46 | 1.57 (ov) | 26.3, $CH_2$ |
| | | 1.43 (ov) | |
| | 47 | 2.87 (dt, 10.1, 6.6) | 47.0, $CH_2$ |
| | | 2.78 (dt, 10.1, 6.6) | |

The stereochemistry was predicted by analyzing the *rdb1* BGC.

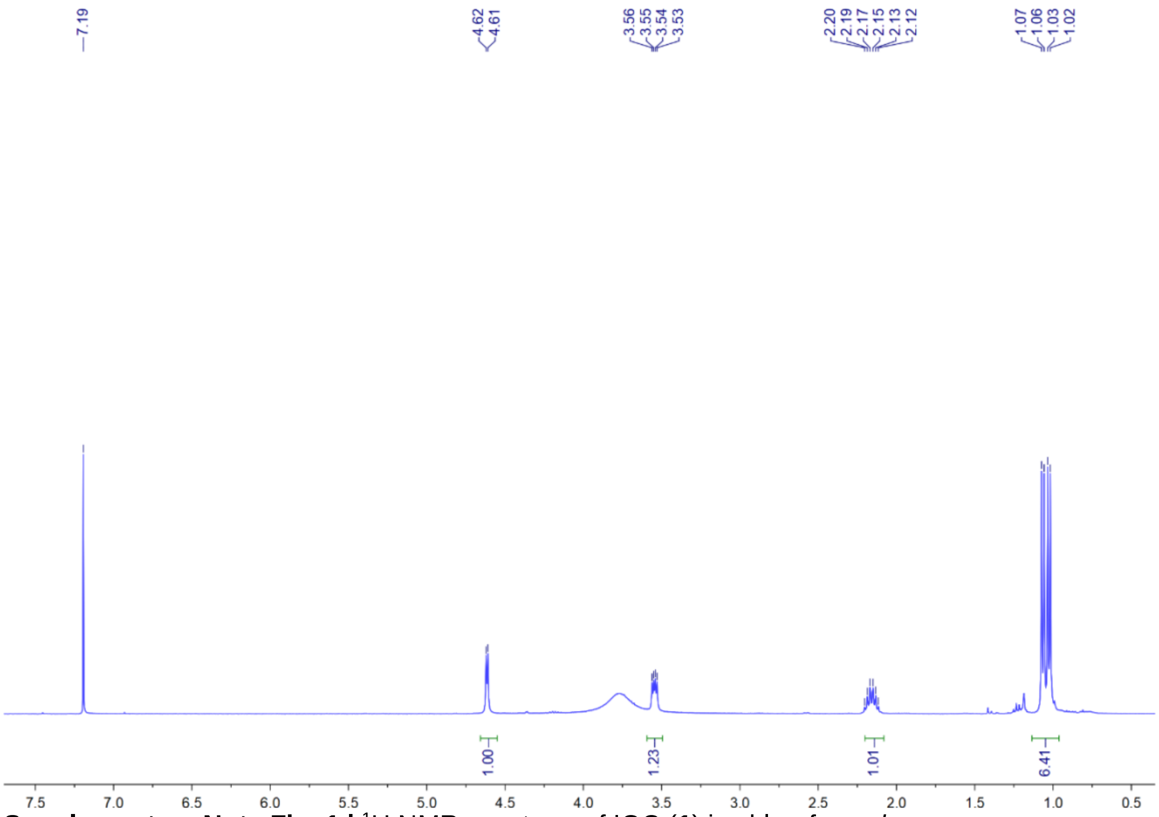


**Supplementary Note Fig. 1 |** $^{1}H$ NMR spectrum of IOC (**1**) in chloroform-*d*.

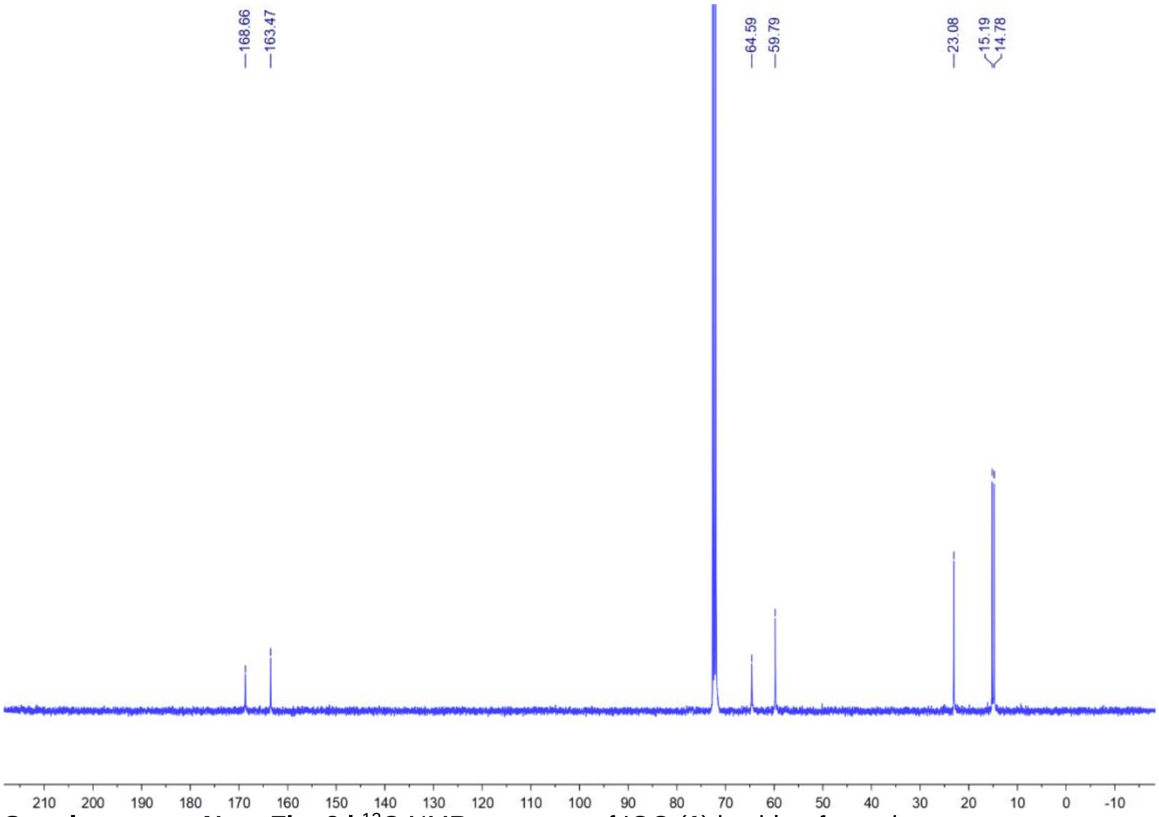


**Supplementary Note Fig. 2 |** $^{13}C$ NMR spectrum of IOC (**1**) in chloroform-*d*.

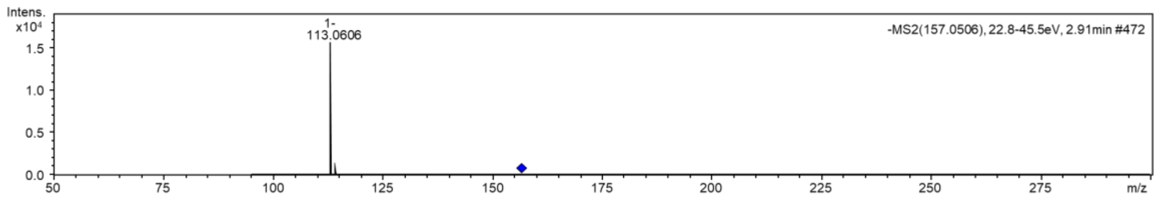

**Supplementary Note Fig. 3 |** HR-ESI-MS of IOC (**1**).

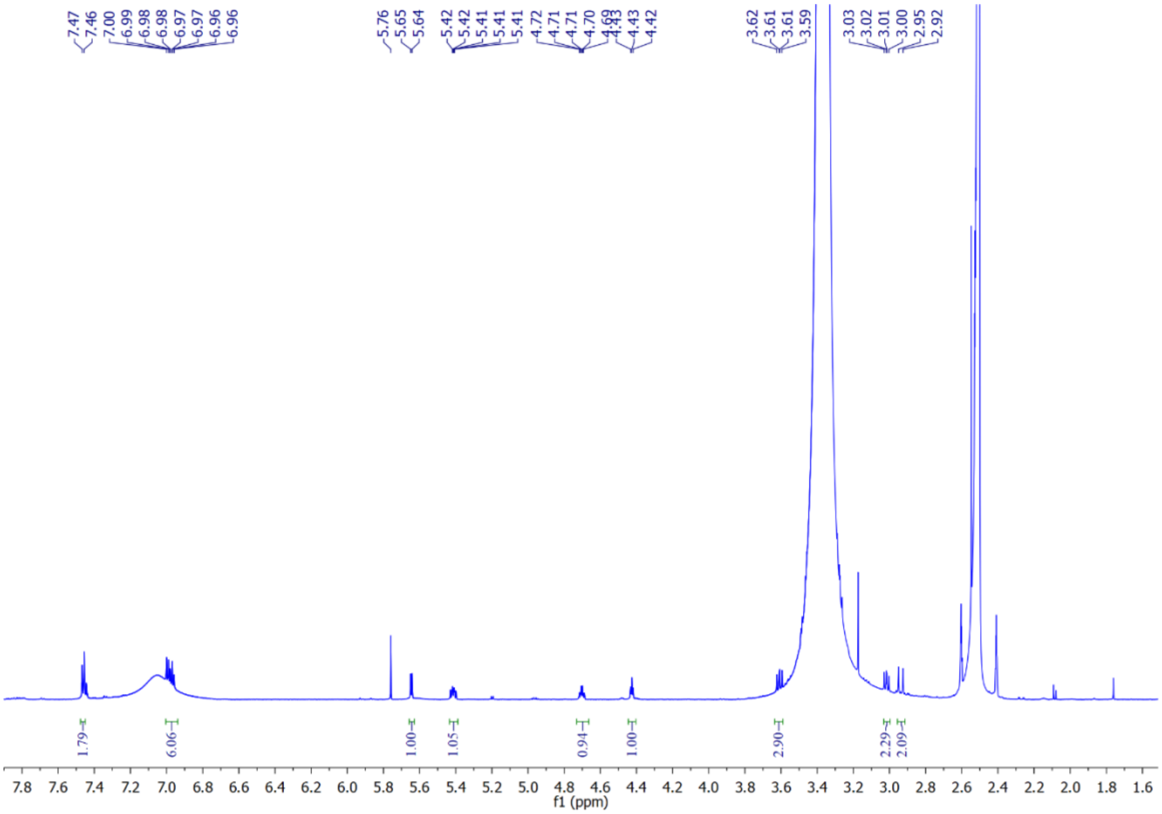

**Supplementary Note Fig. 4 |** [1]H NMR spectrum of photoxenobactin A (**4**) in DMSO-*d*6.

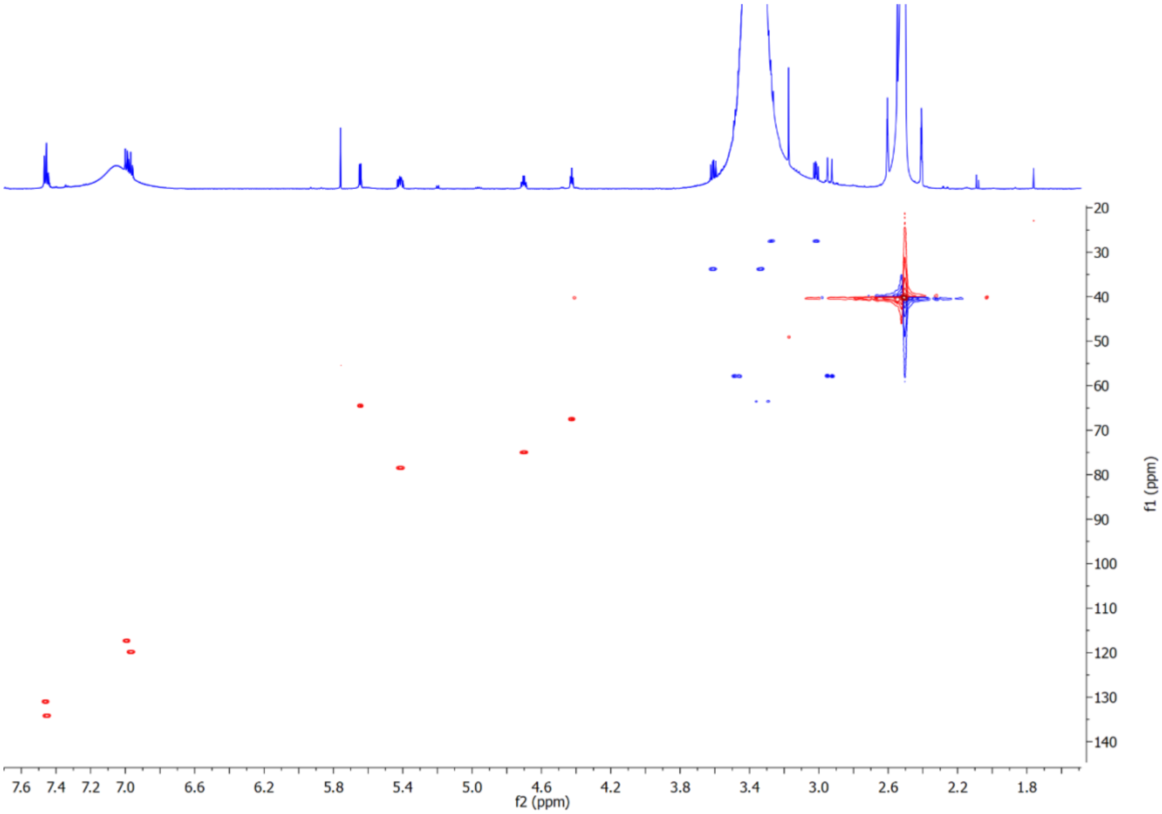

**Supplementary Note Fig. 5 |** HSQC spectrum of photoxenobactin A (**4**) in DMSO-*d*6.

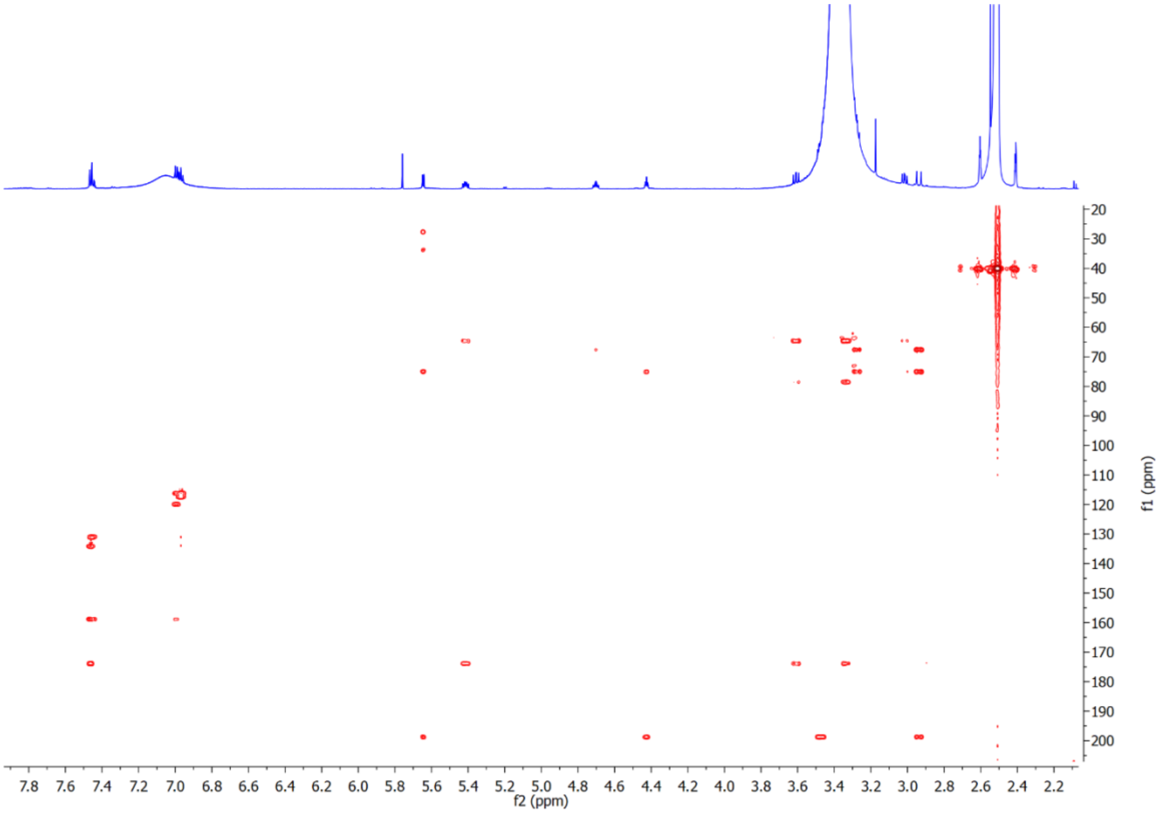


**Supplementary Note Fig. 6 |** HMBC spectrum of photoxenobactin A (**4**) in DMSO-$d_6$.

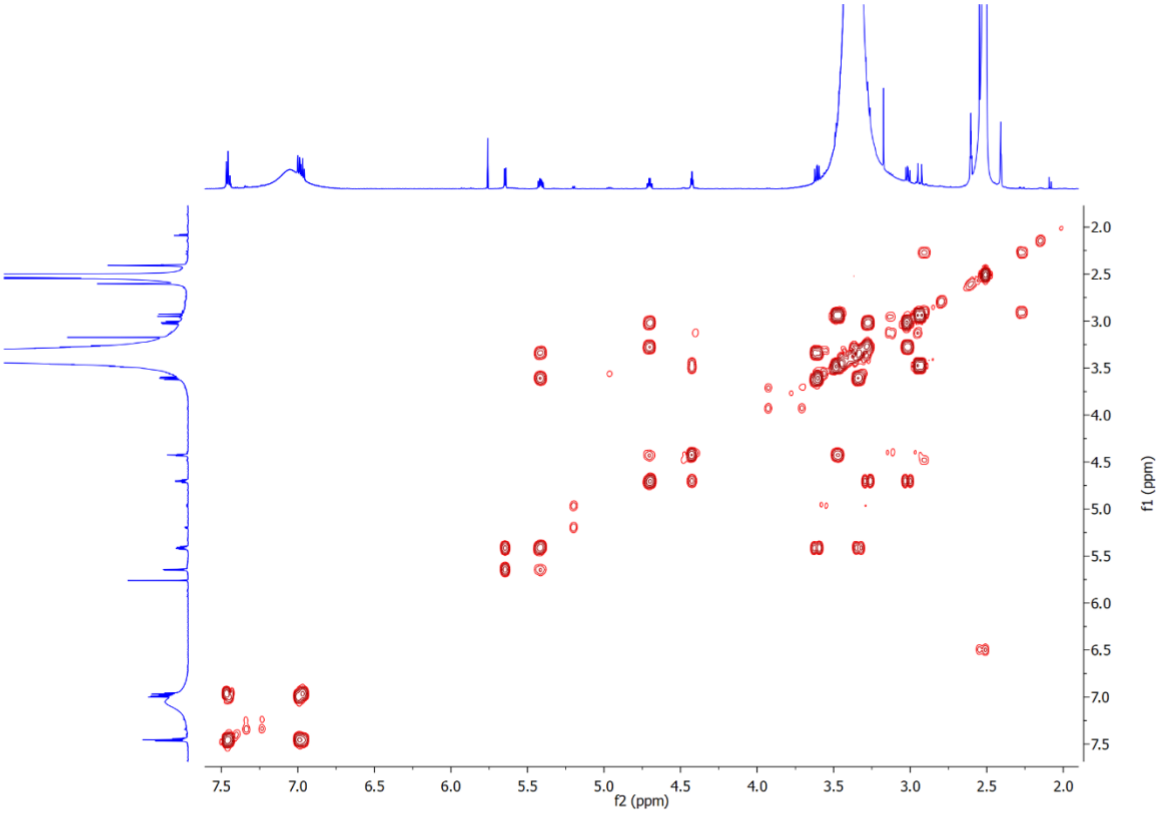


**Supplementary Note Fig. 7 |** $^1$H-$^1$H COSY spectrum of photoxenobactin A (**4**) in DMSO-$d_6$.

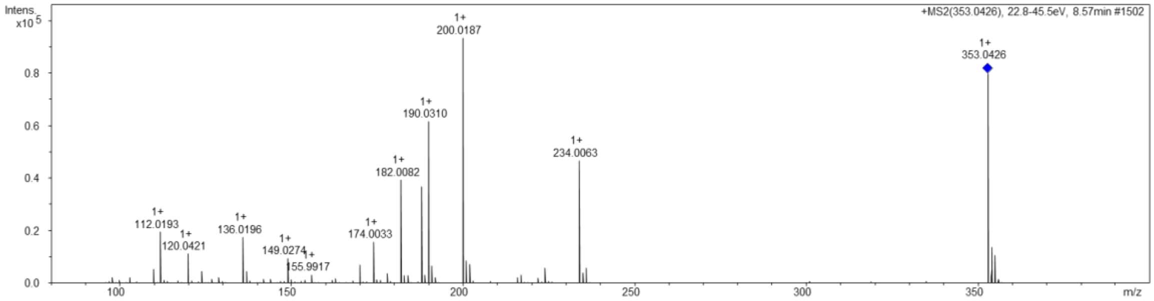


**Supplementary Note Fig. 8 |** HR-ESI-MS of photoxenobactin A (**4**).

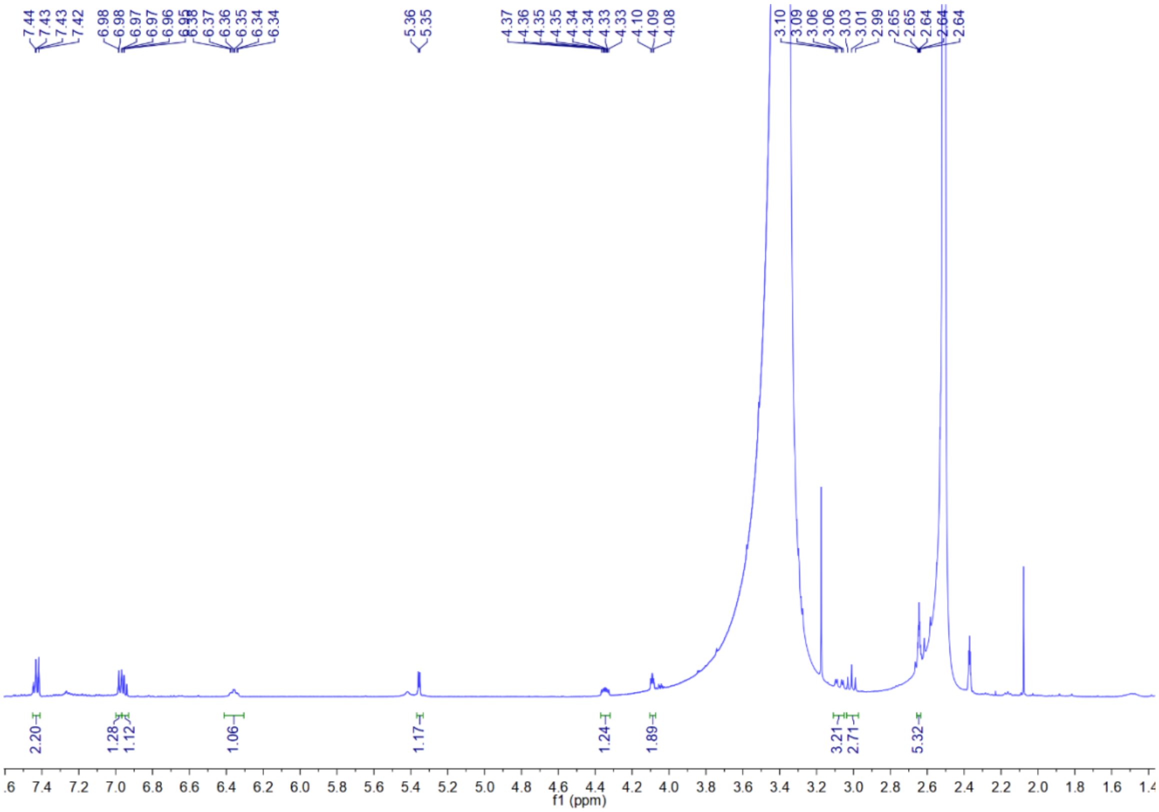

**Supplementary Note Fig. 9 |** $^{1}$H NMR spectrum of photoxenobactin B (**5**) in DMSO-$d_6$.

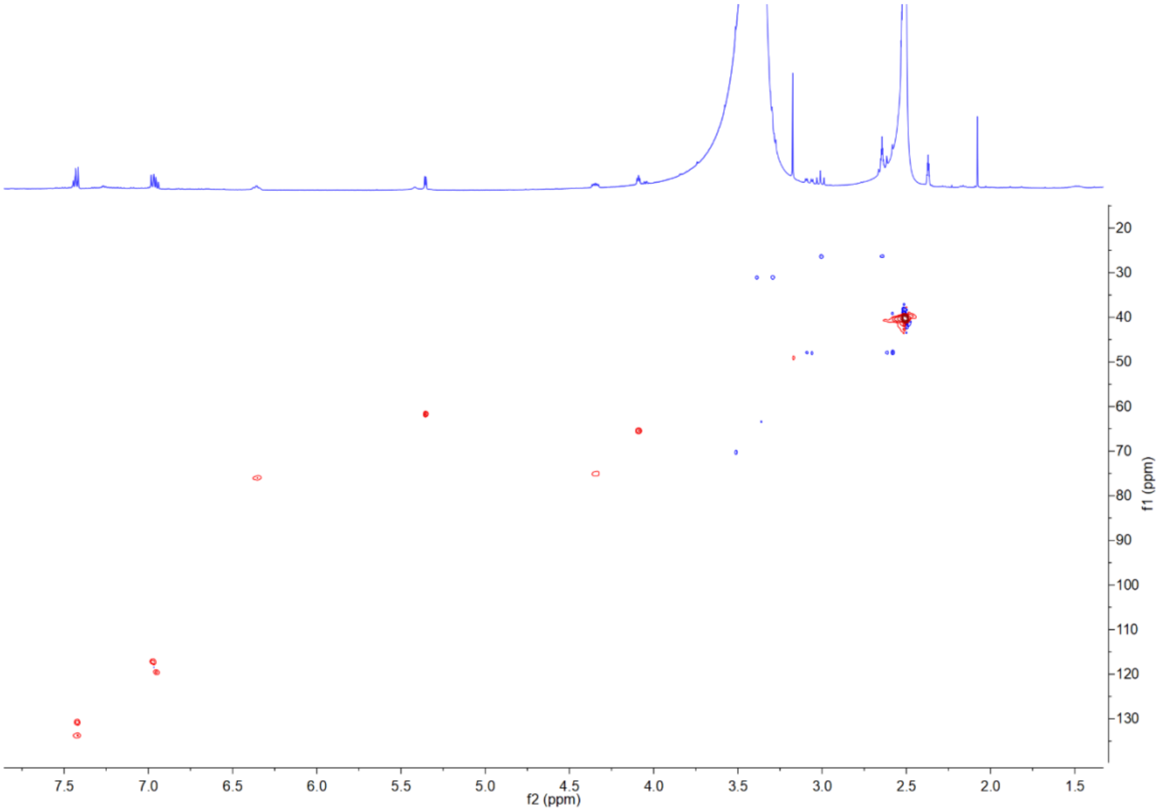

**Supplementary Note Fig. 10 |** HSQC spectrum of photoxenobactin B (**5**) in DMSO-$d_6$.

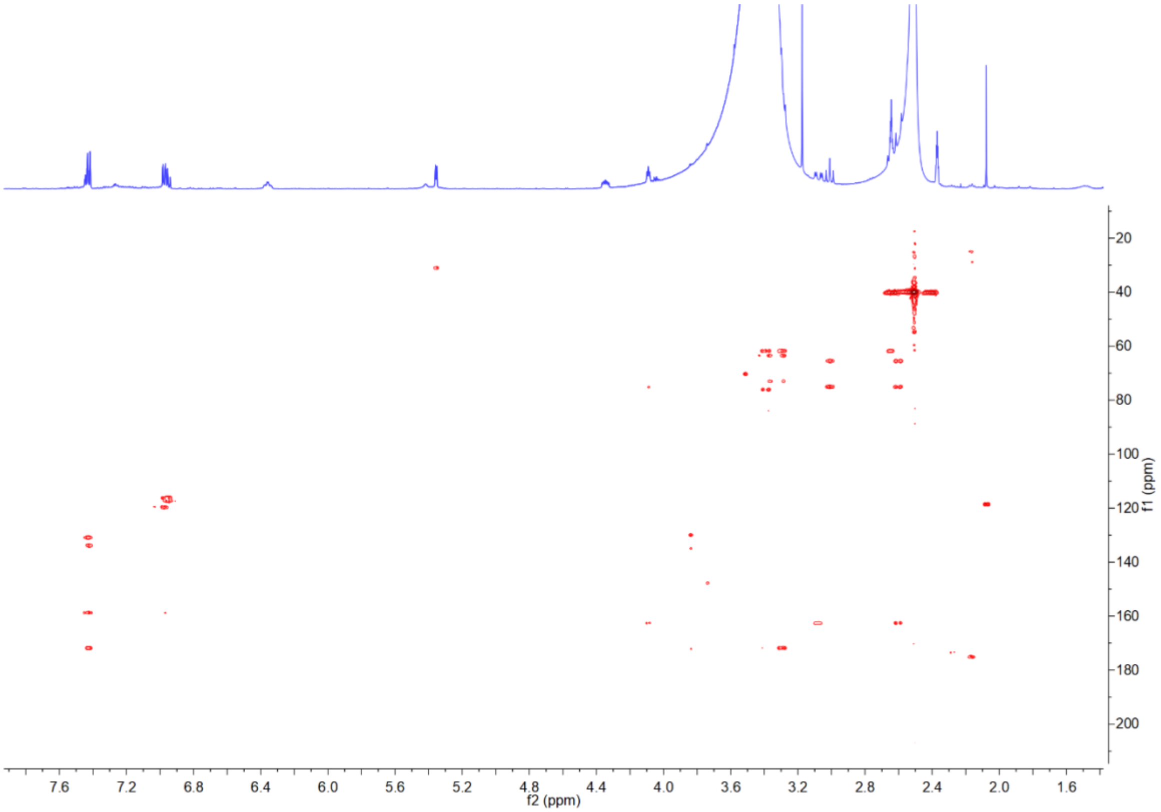


**Supplementary Note Fig. 11 |** HMBC spectrum of photoxenobactin B (**5**) in DMSO-$d_6$.

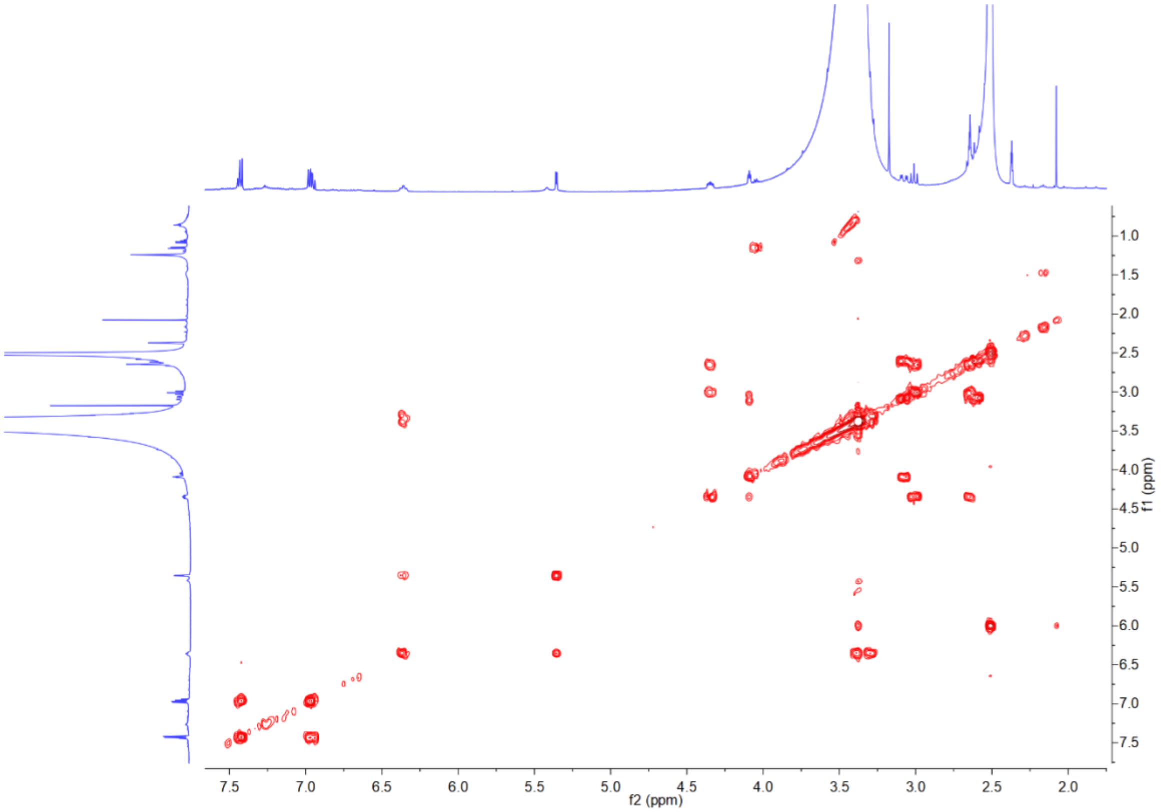


**Supplementary Note Fig. 12 |** $^1$H-$^1$H COSY spectrum of photoxenobactin B (**5**) in DMSO-$d_6$.

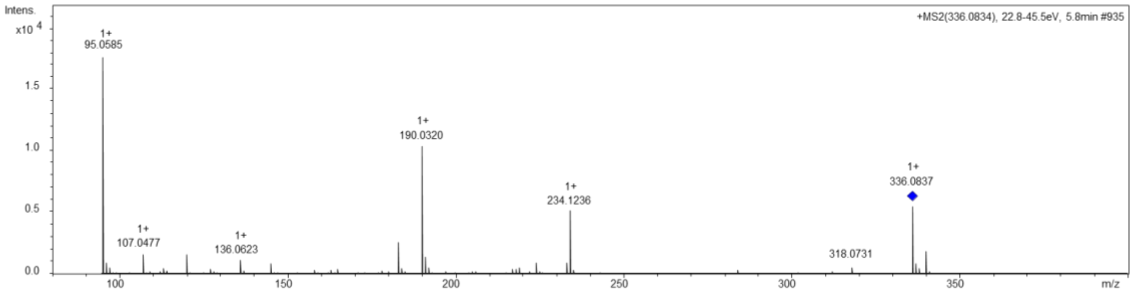


**Supplementary Note Fig. 13 |** HR-ESI-MS of photoxenobactin B (**5**).

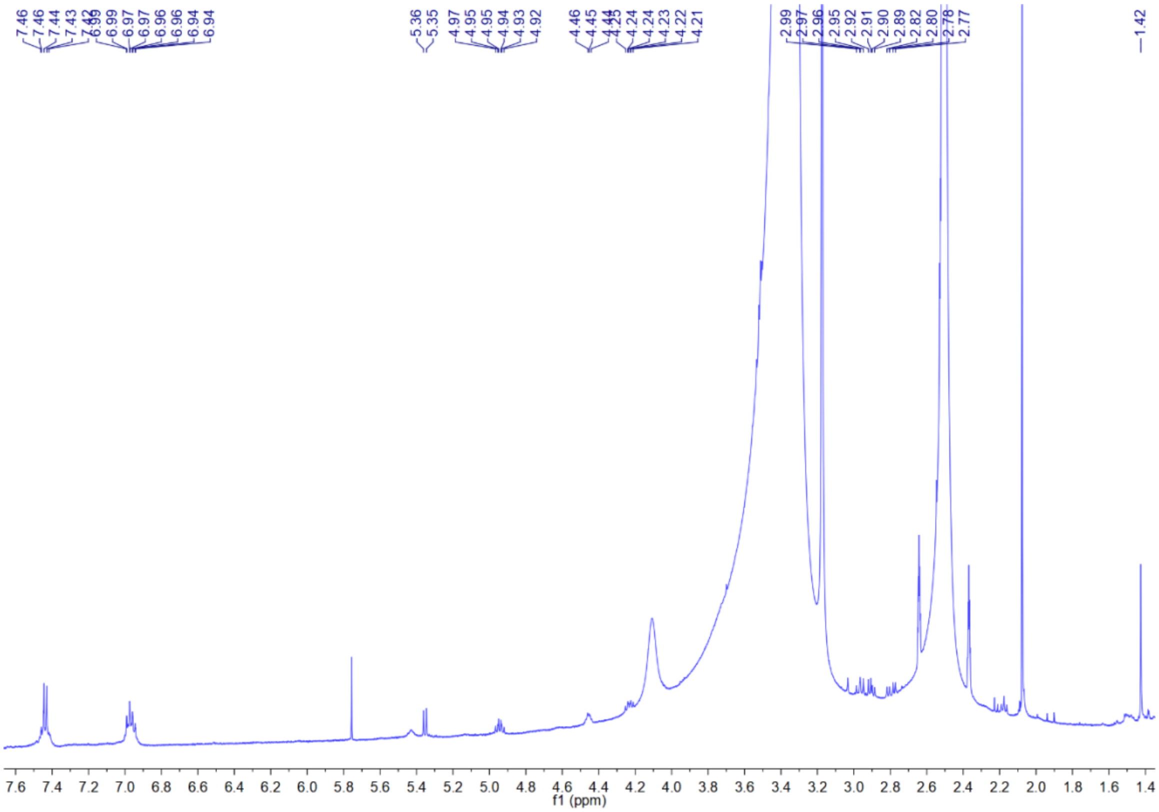

**Supplementary Note Fig. 14 |** $^{1}H$ NMR spectrum of photoxenobactin C (**6**) in DMSO-$d_6$.

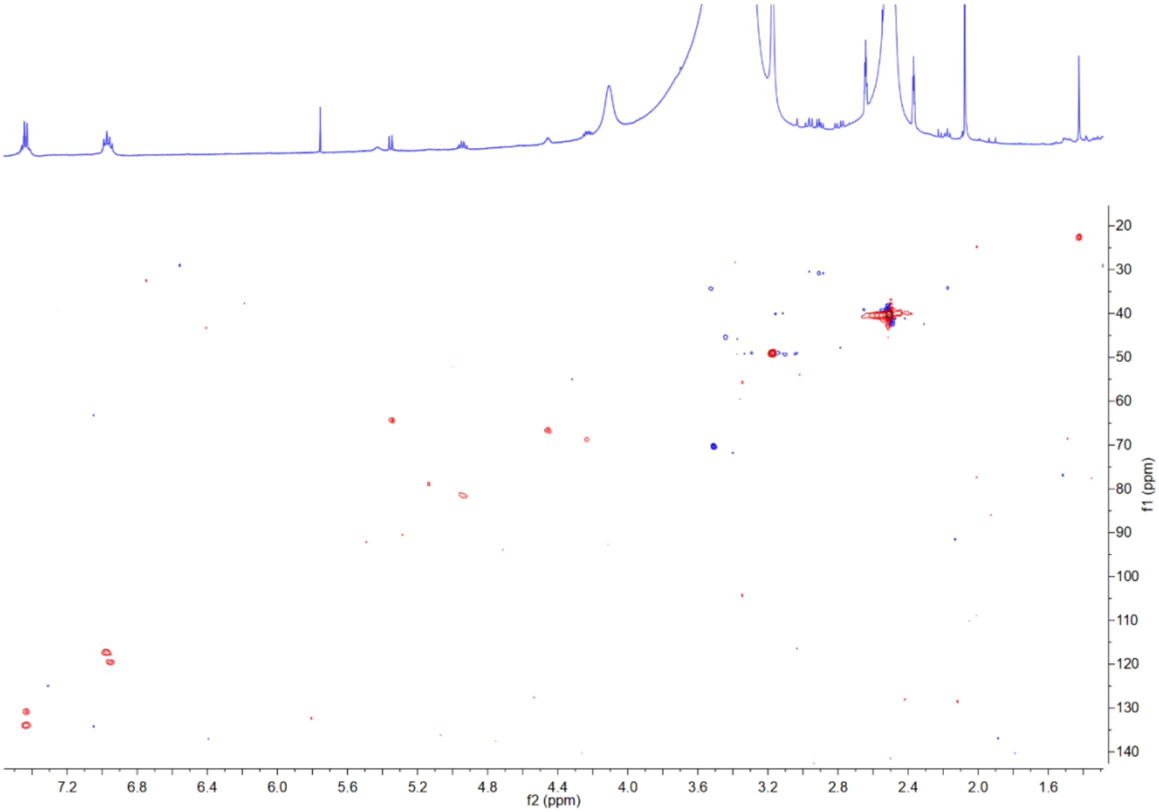

**Supplementary Note Fig. 15 |** HSQC spectrum of photoxenobactin C (**6**) in DMSO-$d_6$.

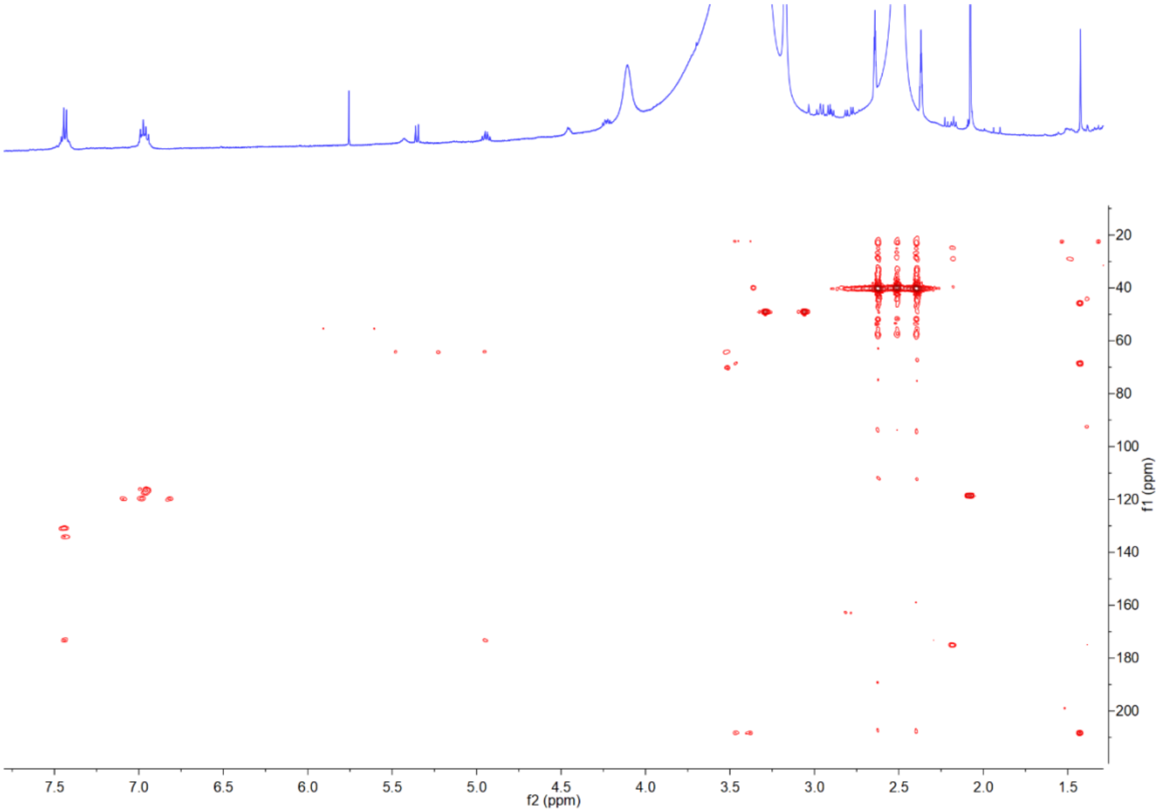


**Supplementary Note Fig. 16 |** HMBC spectrum of photoxenobactin C (**6**) in DMSO-$d_6$.

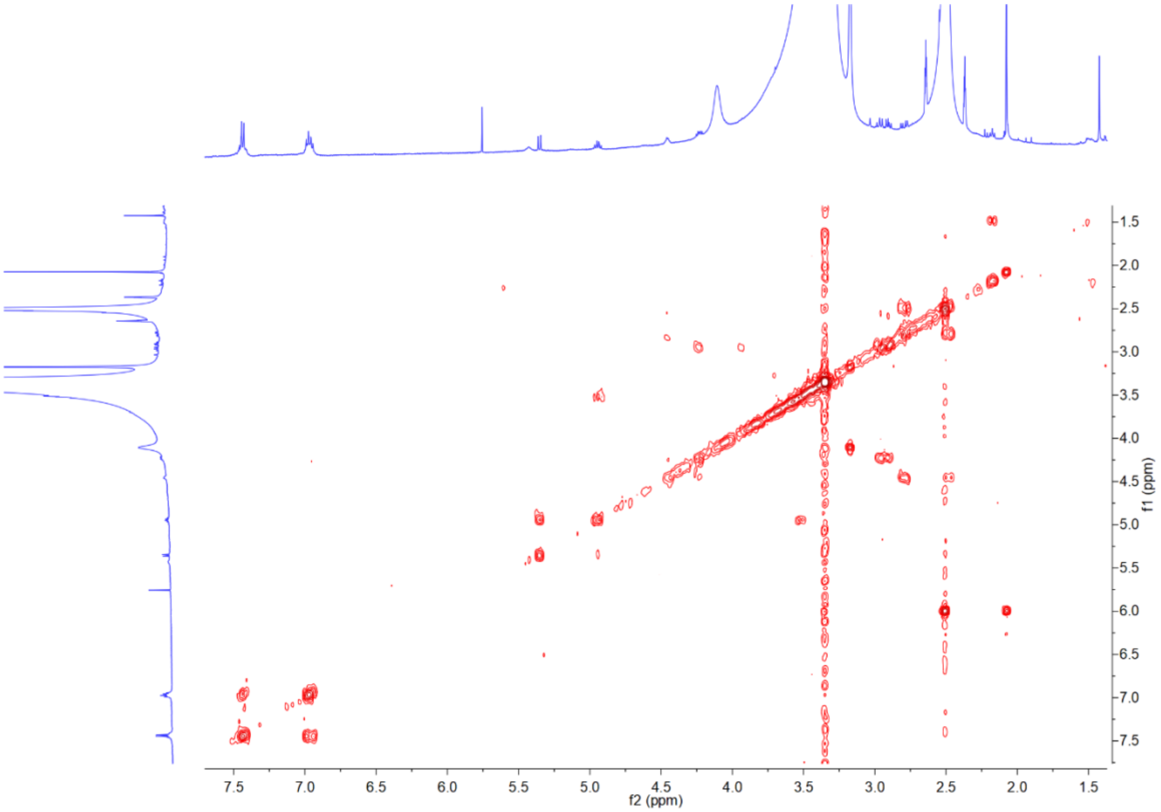


**Supplementary Note Fig. 17 |** $^1$H-$^1$H COSY spectrum of photoxenobactin C (**6**) in DMSO-$d_6$.

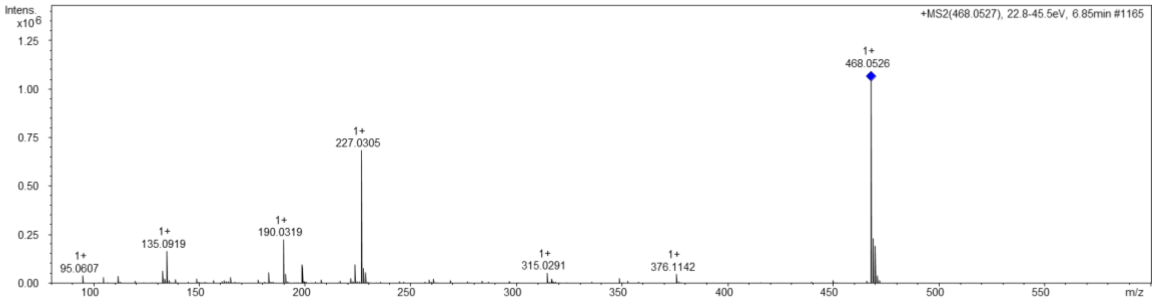


**Supplementary Note Fig. 18 |** HR-ESI-MS of photoxenobactin C (**6**).

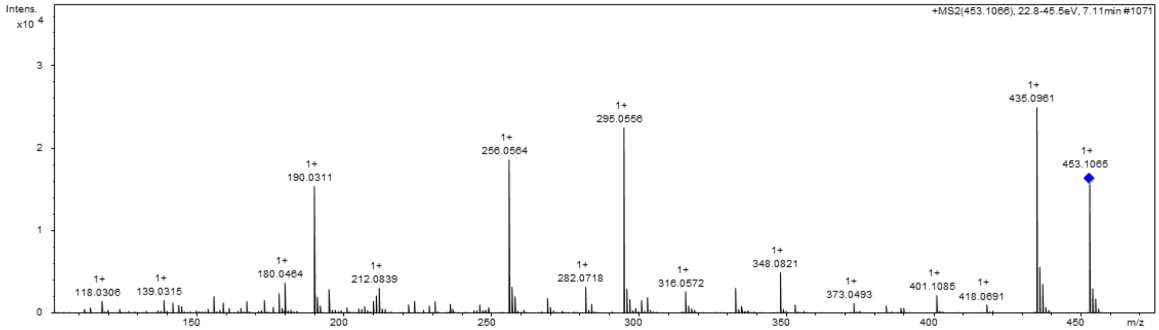


**Supplementary Note Fig. 19 |** HR-ESI-MS of photoxenobactin D (**7**).

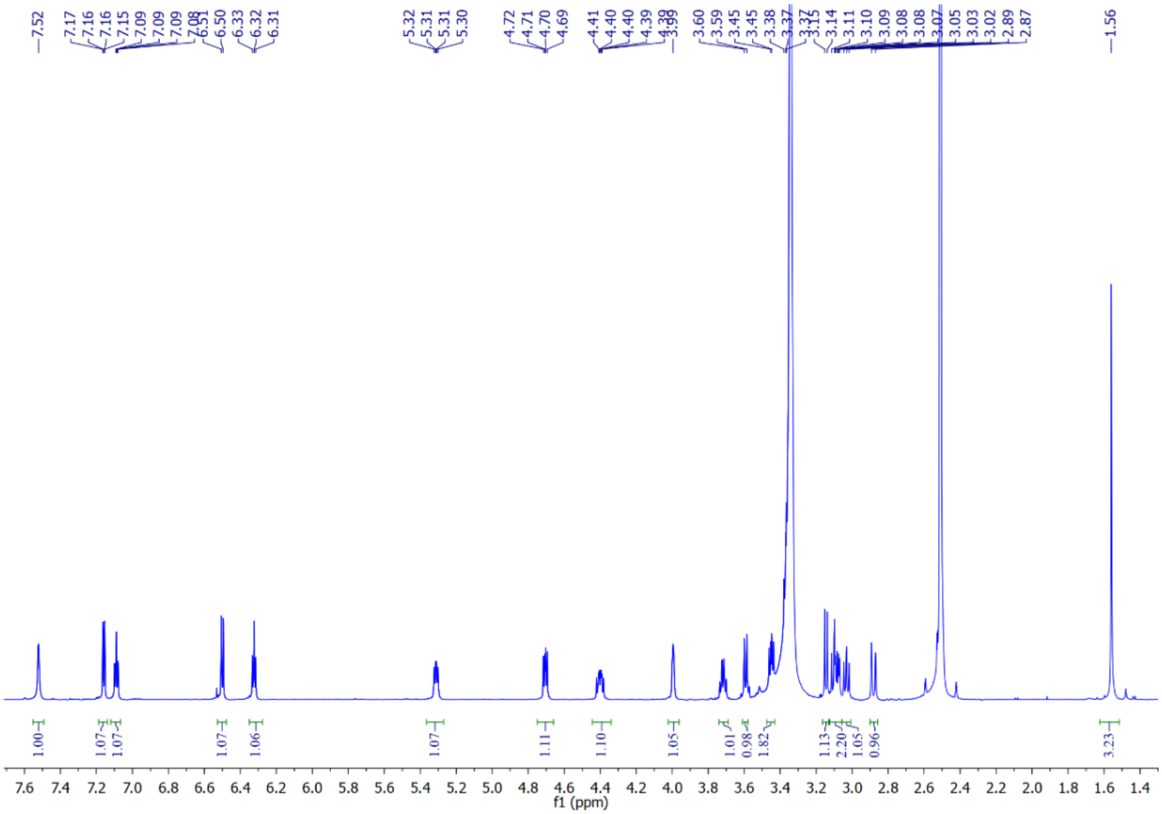

**Supplementary Note Fig. 20 |** $^{1}$H NMR spectrum of photoxenobactin E (**8**) in DMSO-$d_6$.

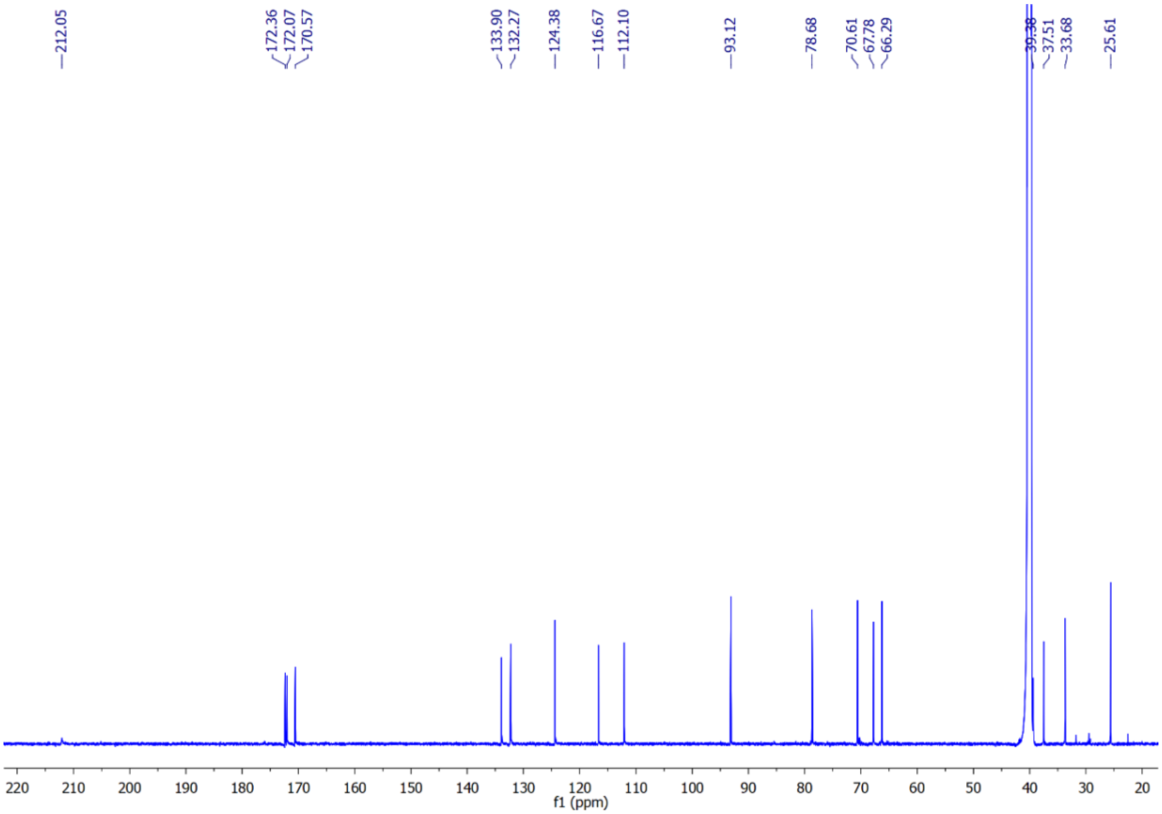

**Supplementary Note Fig. 21 |** $^{13}$C NMR spectrum of photoxenobactin E (**8**) in DMSO-$d_6$.

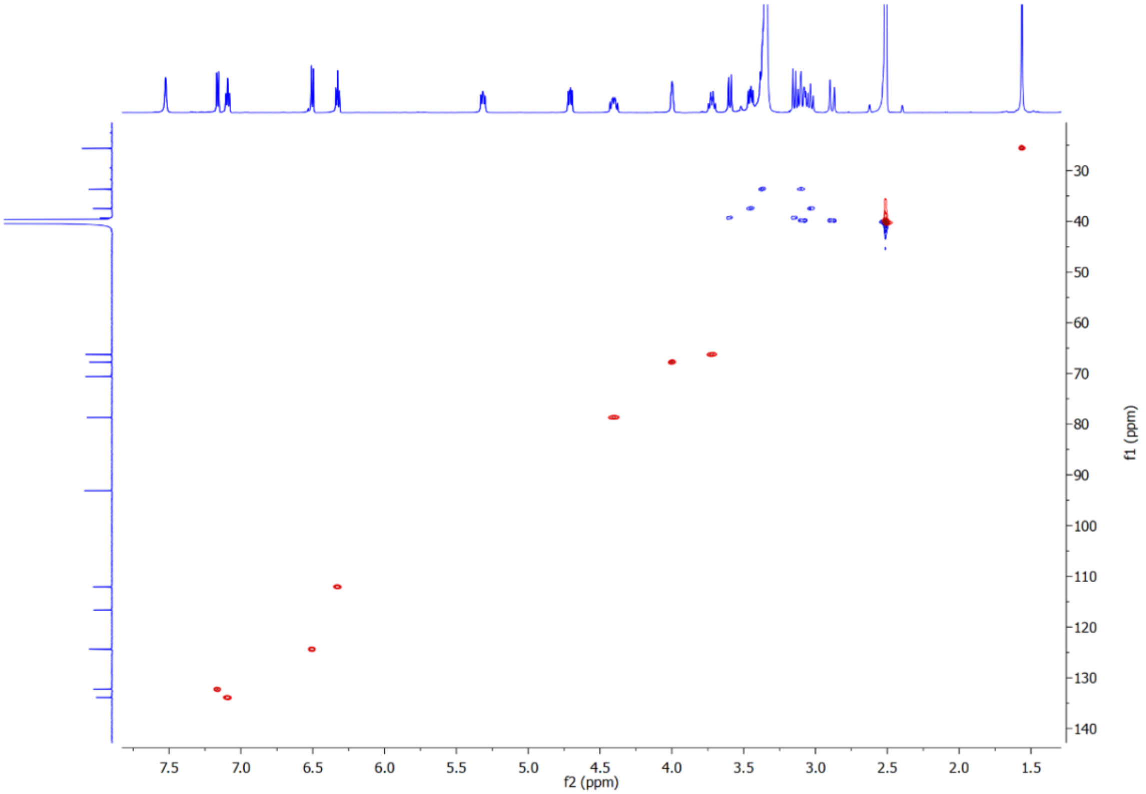


**Supplementary Note Fig. 22 |** HSQC spectrum of photoxenobactin E (**8**) in DMSO-$d_6$.

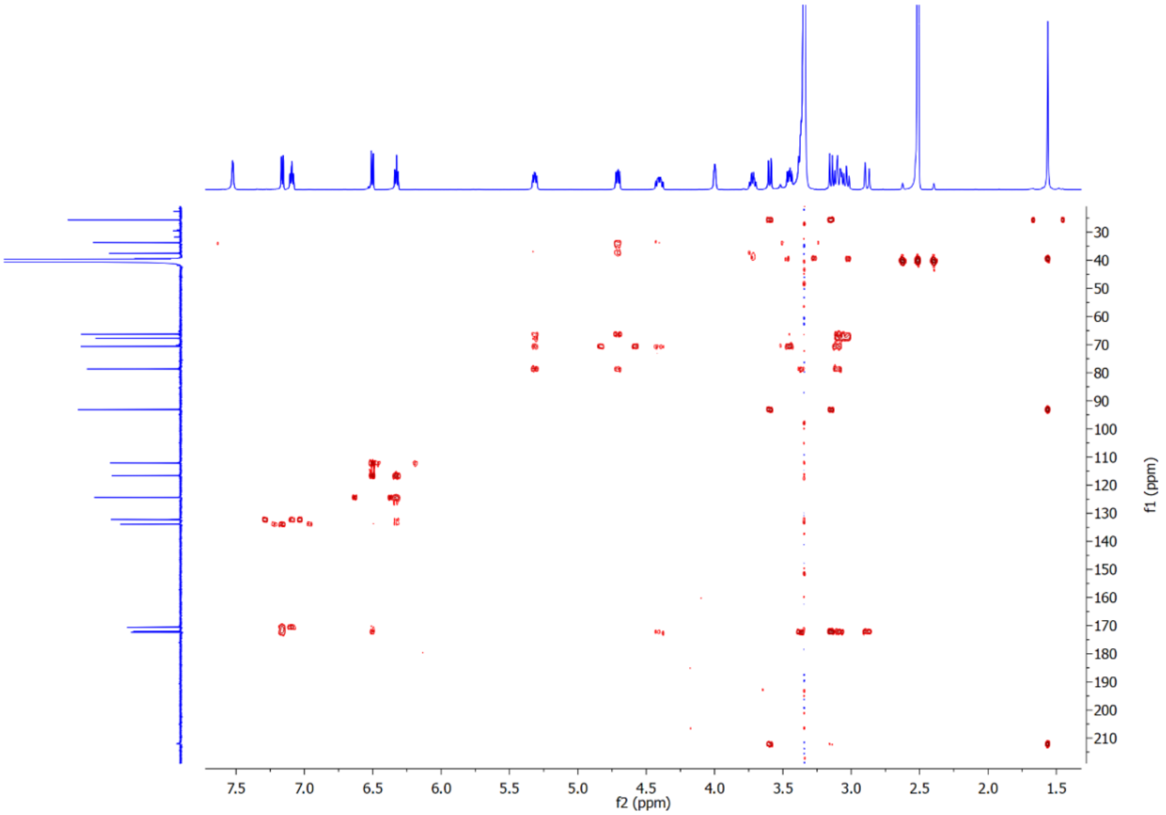


**Supplementary Note Fig. 23 |** HMBC spectrum of photoxenobactin E (**8**) in DMSO-$d_6$.

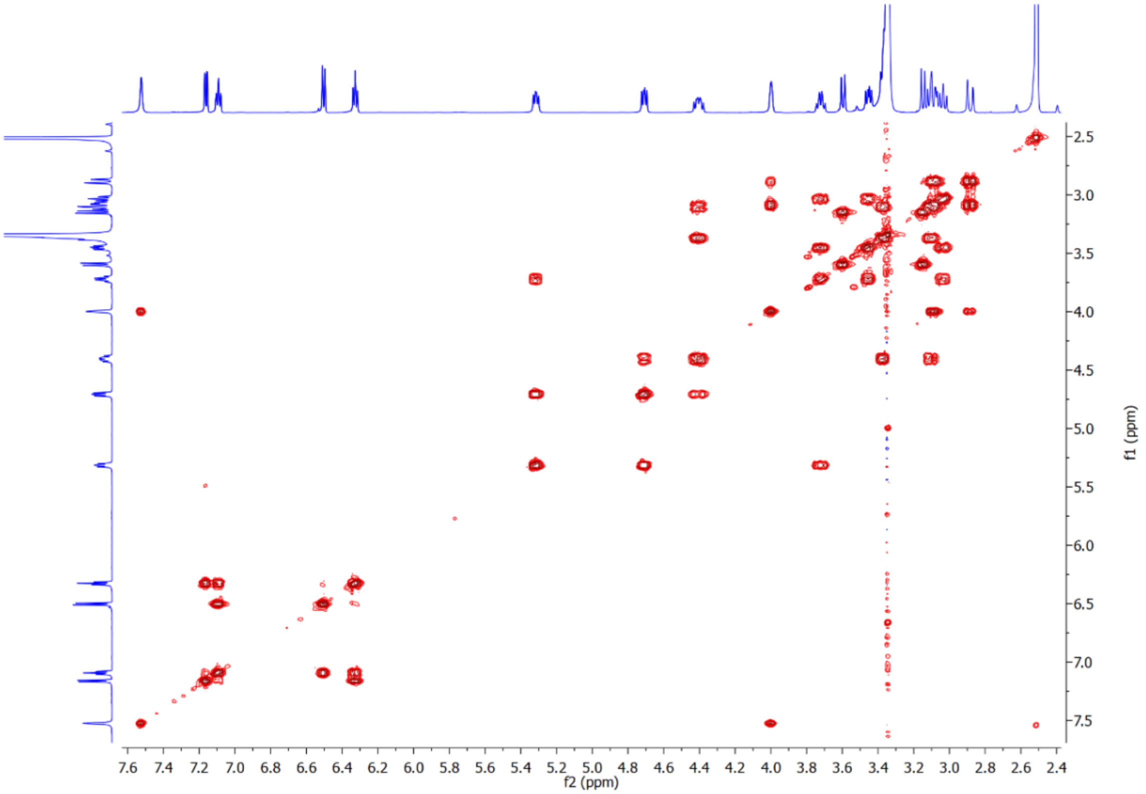


**Supplementary Note Fig. 24 |** $^1H$-$^1H$ COSY spectrum of photoxenobactin E (**8**) in DMSO-$d_6$.

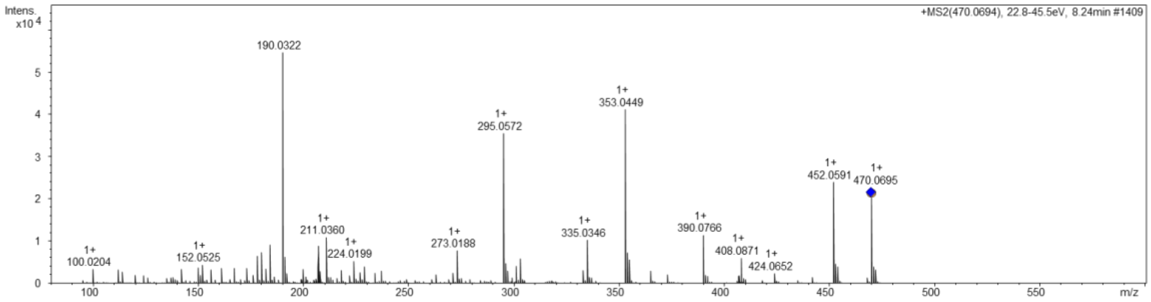


**Supplementary Note Fig. 25 |** HR-ESI-MS of photoxenobactin E (**8**).

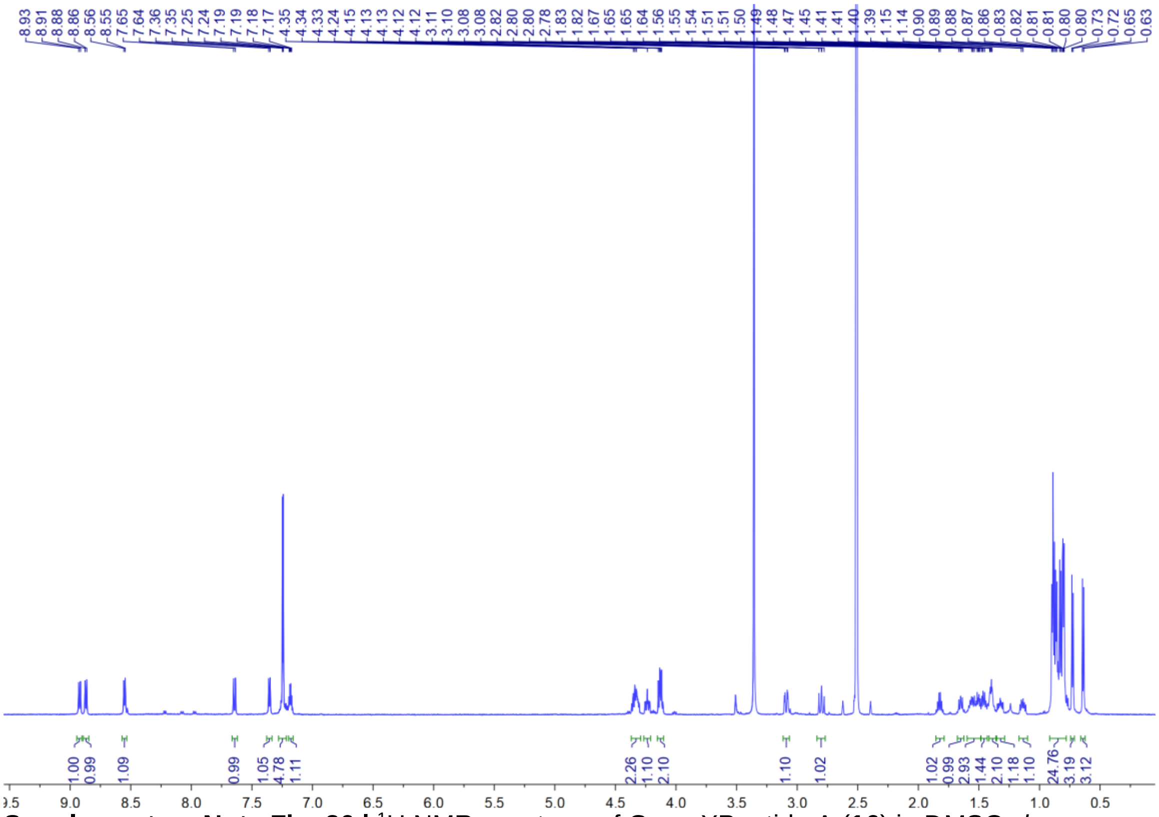

**Supplementary Note Fig. 26 |** $^{1}H$ NMR spectrum of GameXPeptide A (**16**) in DMSO-*d*$_6$.

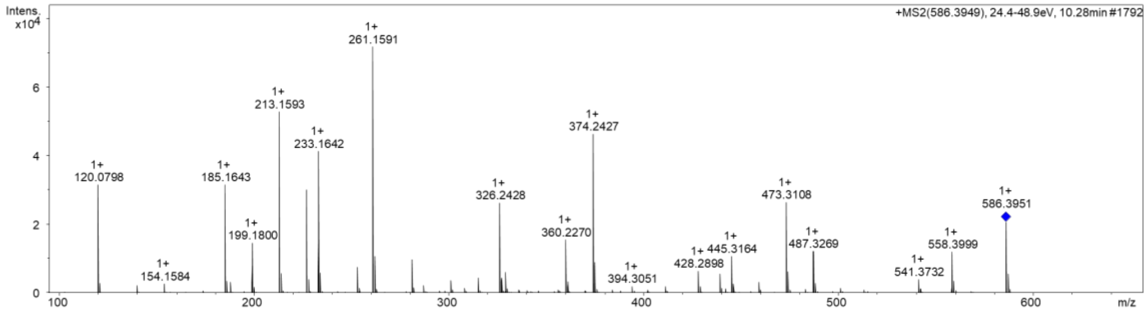


**Supplementary Note Fig. 27 |** HR-ESI-MS of GameXPeptide A (**16**).

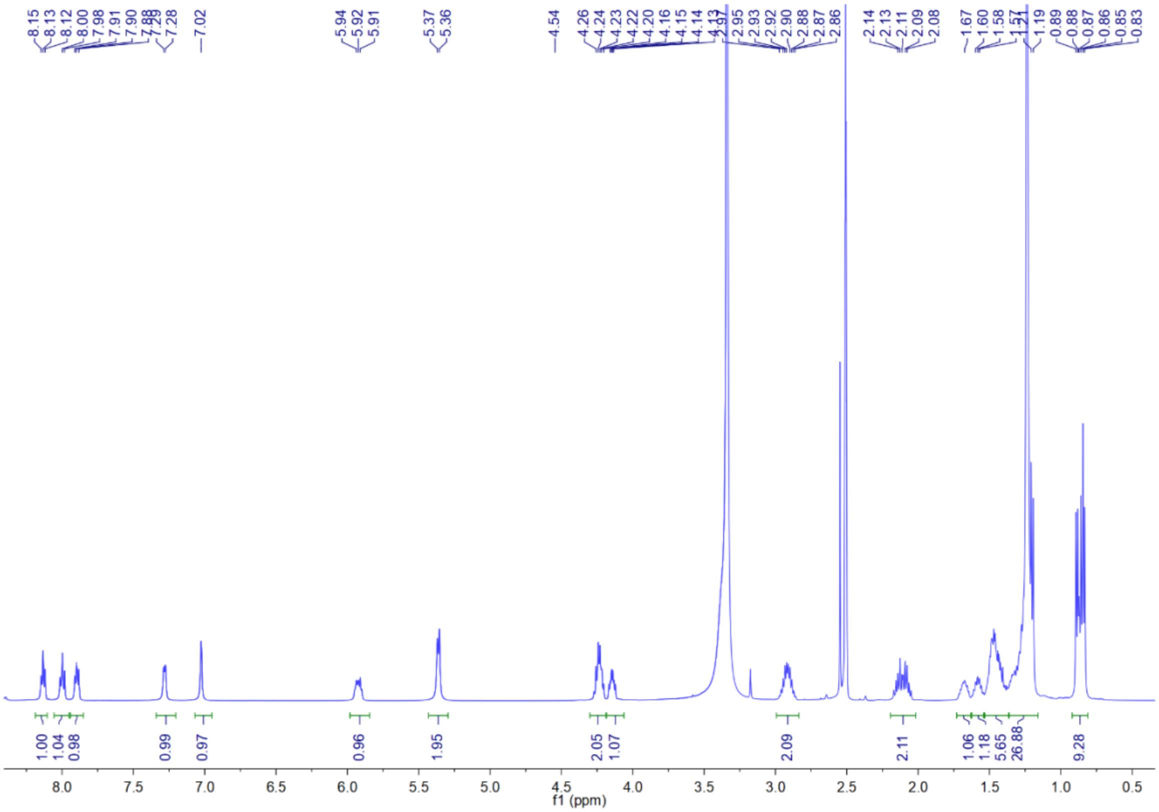

**Supplementary Note Fig. 28 |** $^{1}$H NMR spectrum of lipocitide A (**17**) in DMSO-$d_6$.

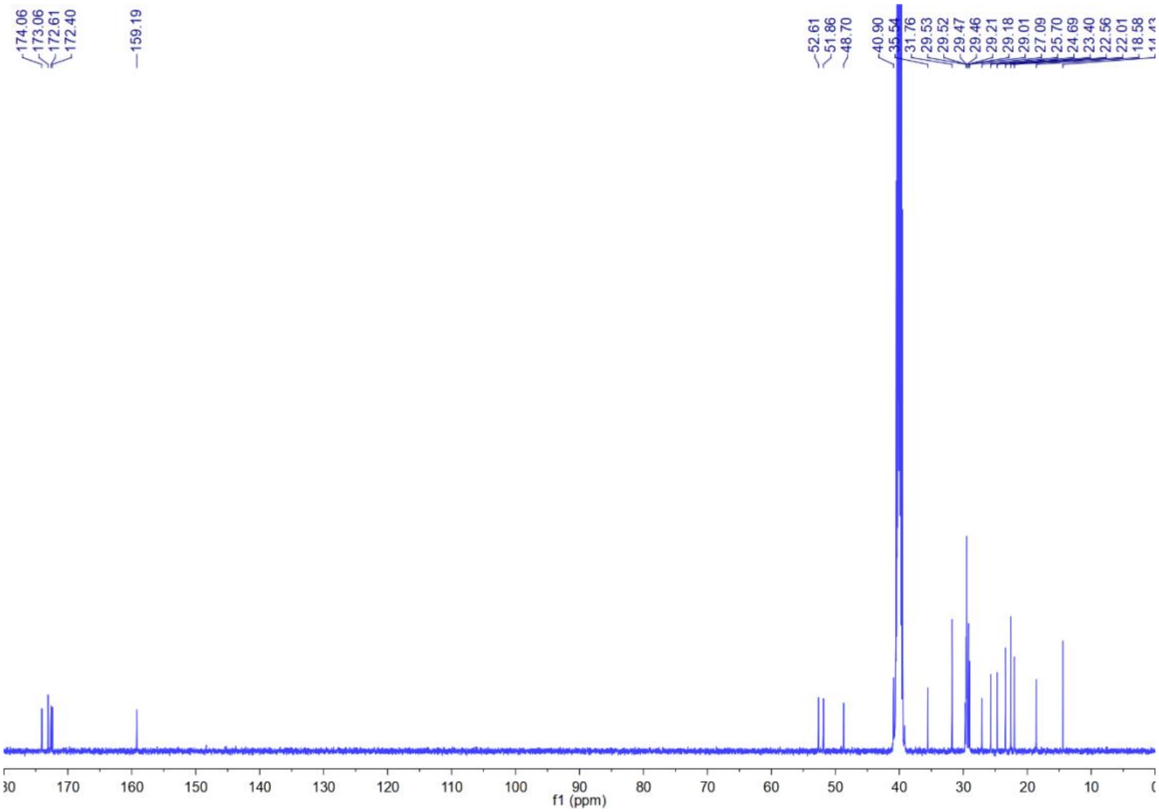

**Supplementary Note Fig. 29 |** $^{13}$C NMR spectrum of lipocitide A (**17**) in DMSO-$d_6$.

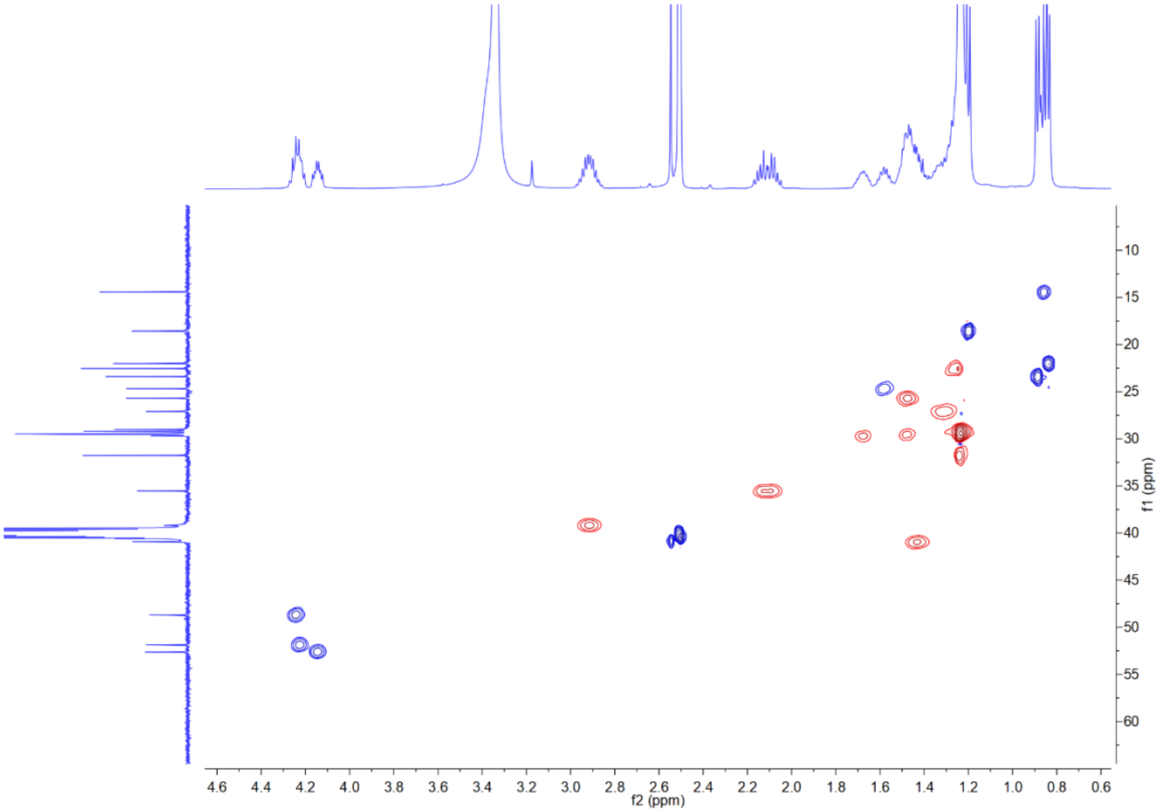

**Supplementary Note Fig. 30 |** HSQC spectrum of lipocitide A (**17**) in DMSO-$d_6$.

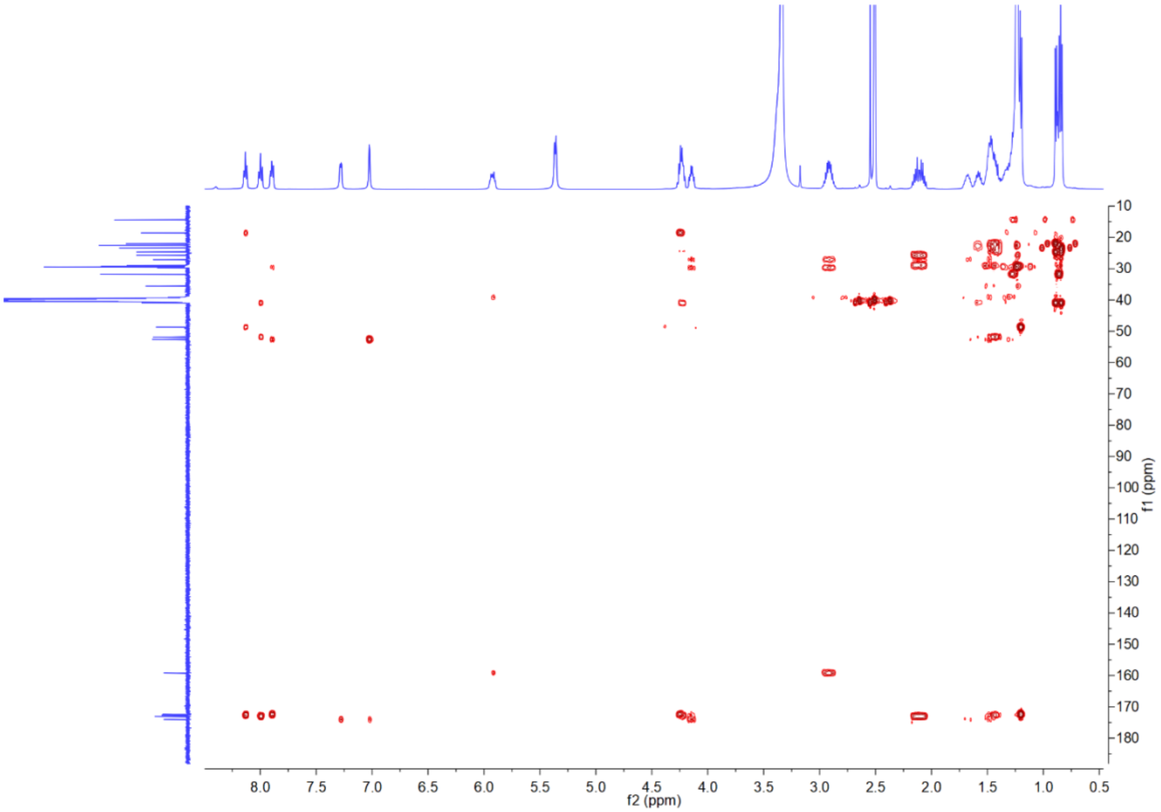

**Supplementary Note Fig. 31 |** HMBC spectrum of lipocitide A (**17**) in DMSO-$d_6$.

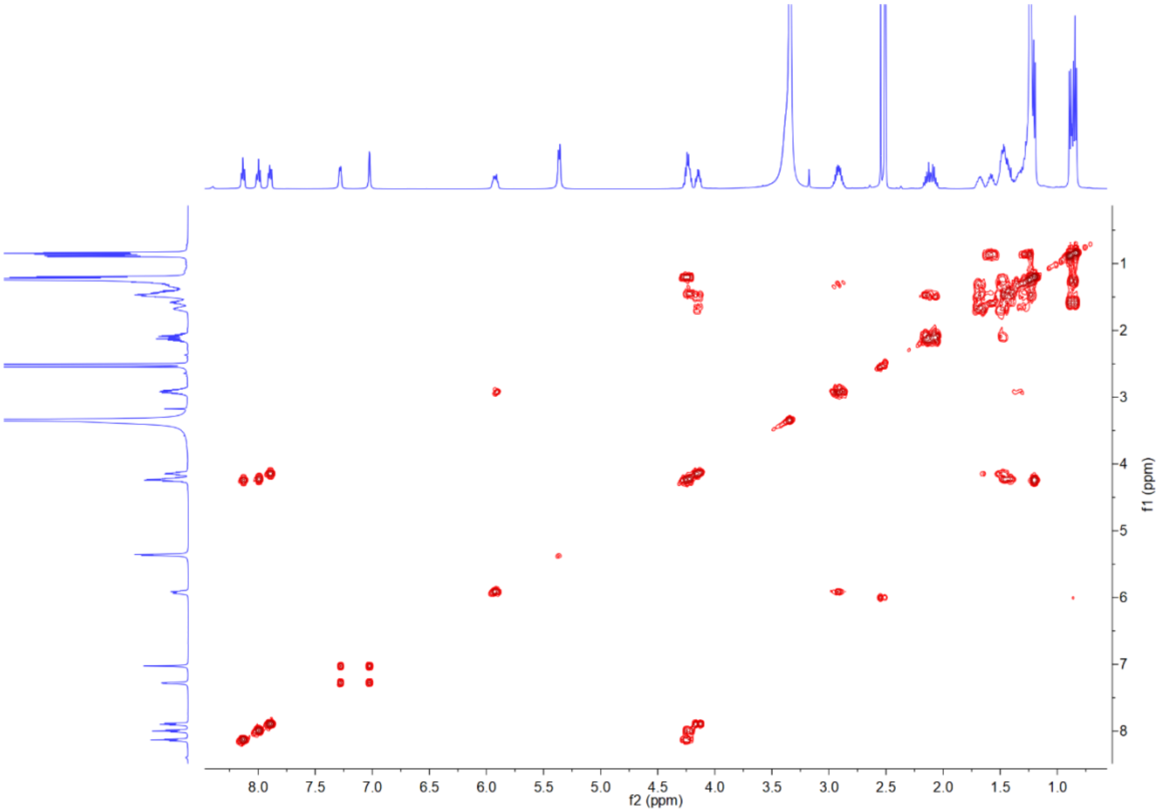


**Supplementary Note Fig. 32 |** $^1H$-$^1H$ COSY spectrum of lipocitide A (**17**) in DMSO-$d_6$.

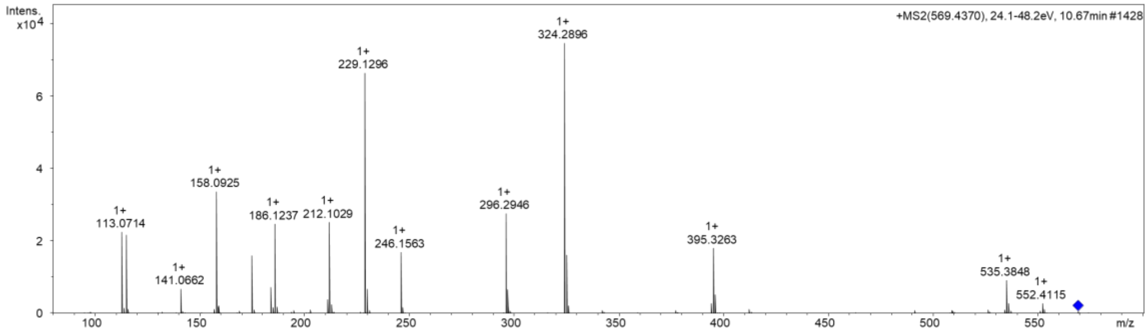


**Supplementary Note Fig. 33 |** HR-ESI-MS of lipocitide A (**17**).

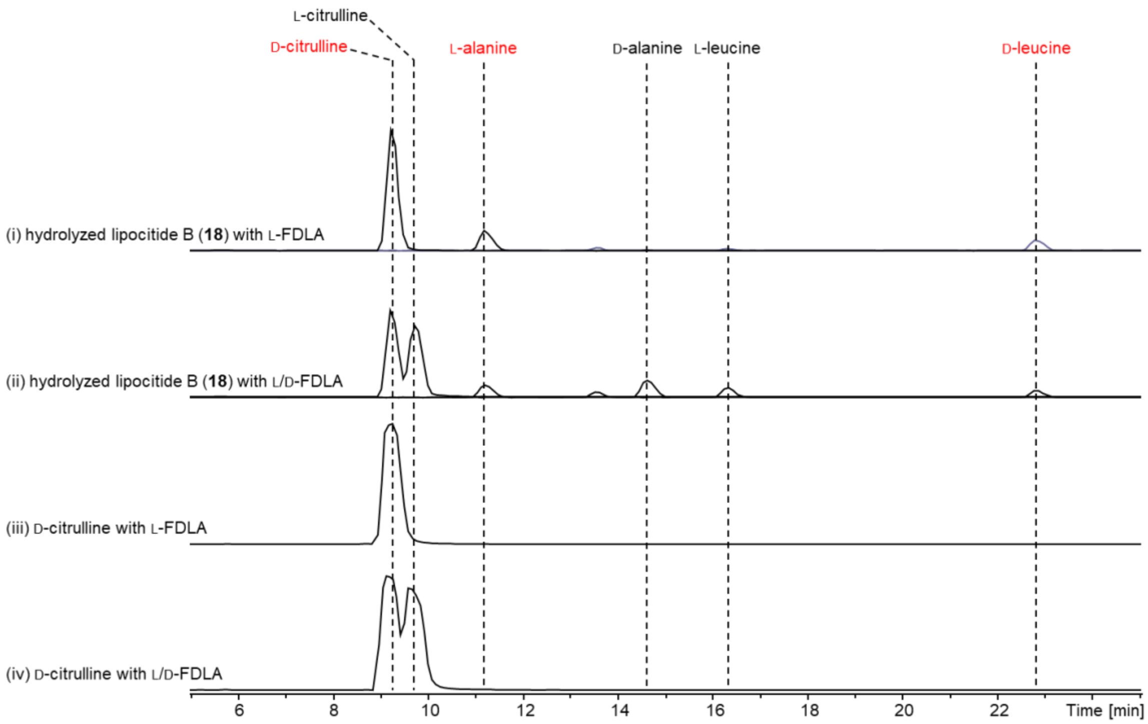


**Supplementary Note Fig. 34 |** Absolute configuration determination of lipocitide B (**18**) by the advanced Marfey's method. HPLC-MS analysis of lipocitide B (**18**) that was hydrolyzed by HCl and subsequently derivatized with L-FDLA and L/D-FDLA. Shown are EICs of FDLA derivatized citrulline, alanine, and leucine. The configuration of amino acids is determined by the elution order. L-FDLA derivatized L-amino acids are eluted prior to D-amino acids [3], except citrulline as illustrated in traces (iii) and (iv). L-FDLA derivatized (iii) D-citrulline has a shorter retention time than (iv) L-citrulline. The determined configuration of an amino acid residue is highlighted in red.

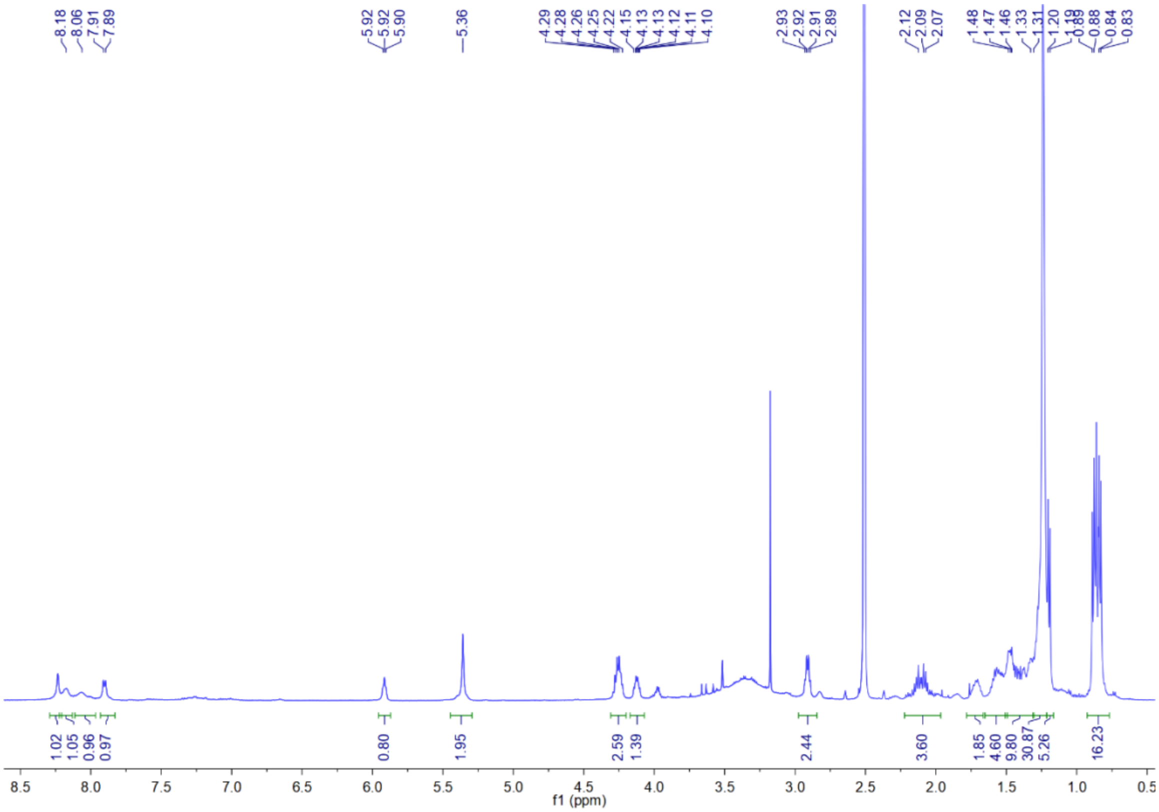

**Supplementary Note Fig. 35 |** $^{1}H$ NMR spectrum of lipocitide B (**18**) in DMSO-$d_6$.

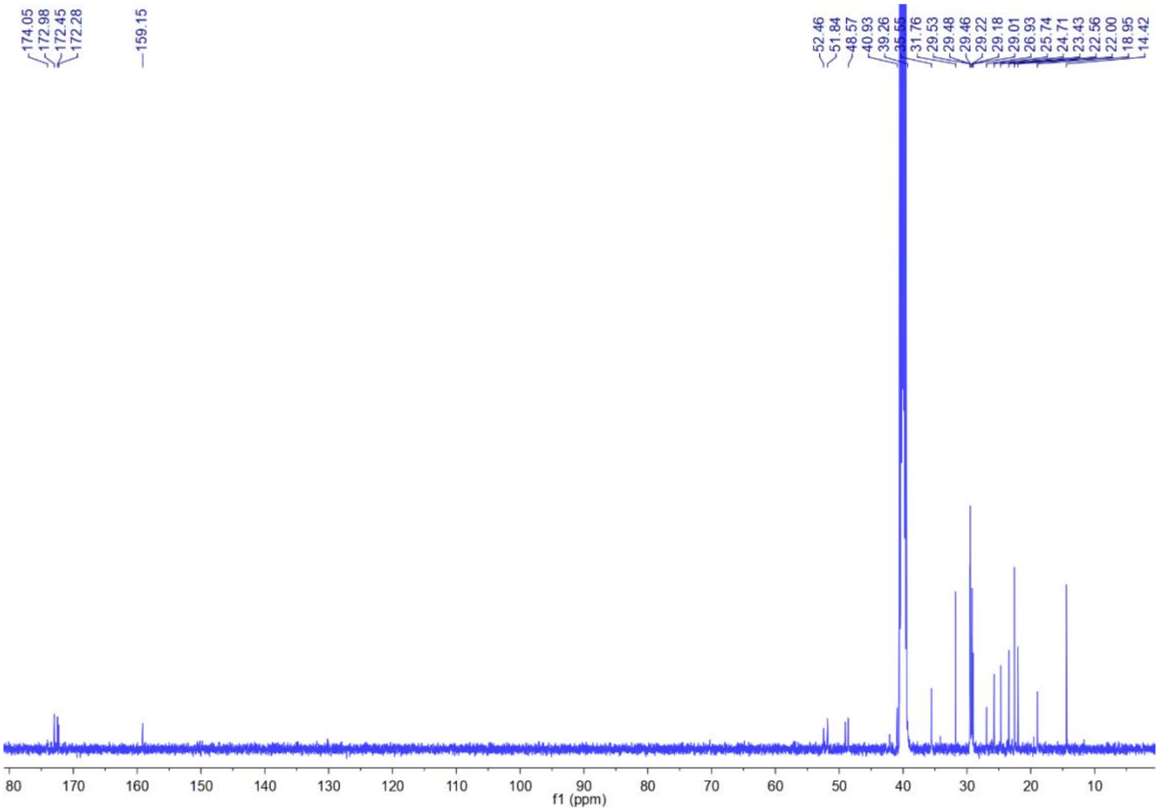

**Supplementary Note Fig. 36 |** $^{13}C$ NMR spectrum of lipocitide B (**18**) in DMSO-$d_6$.

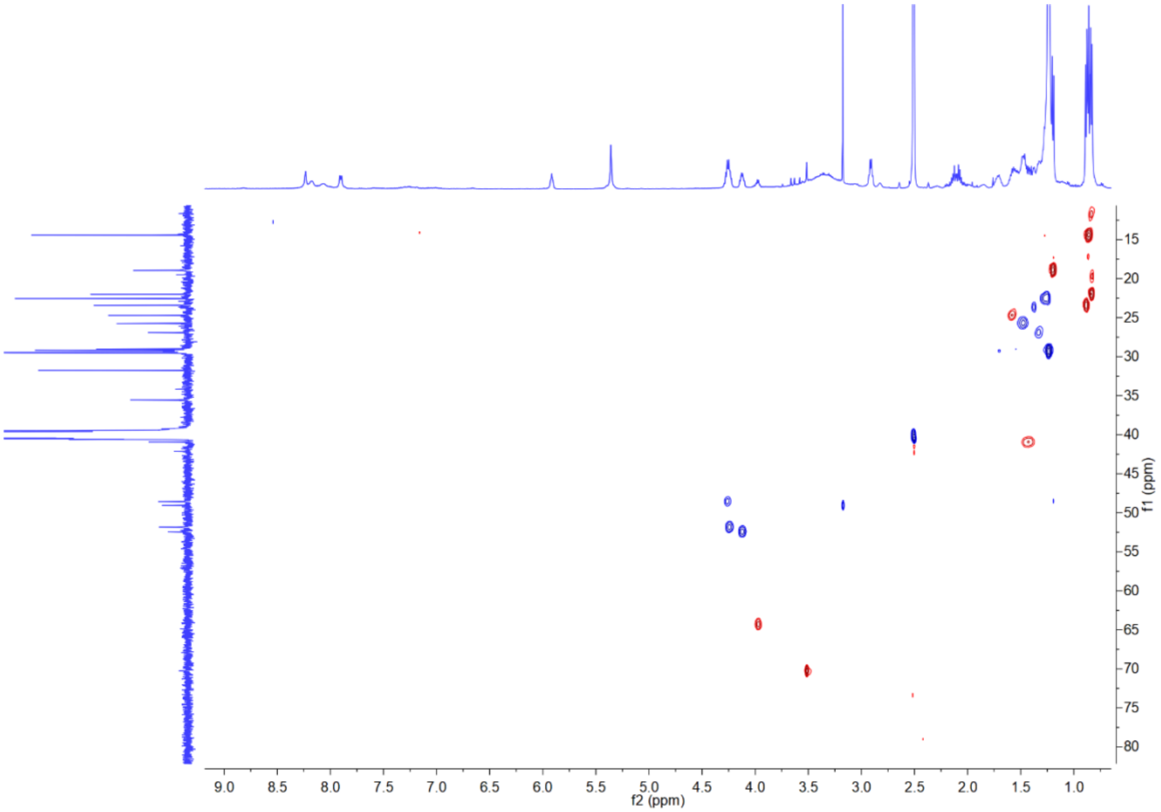


**Supplementary Note Fig. 37 |** HSQC spectrum of lipocitide B (**18**) in DMSO-$d_6$.

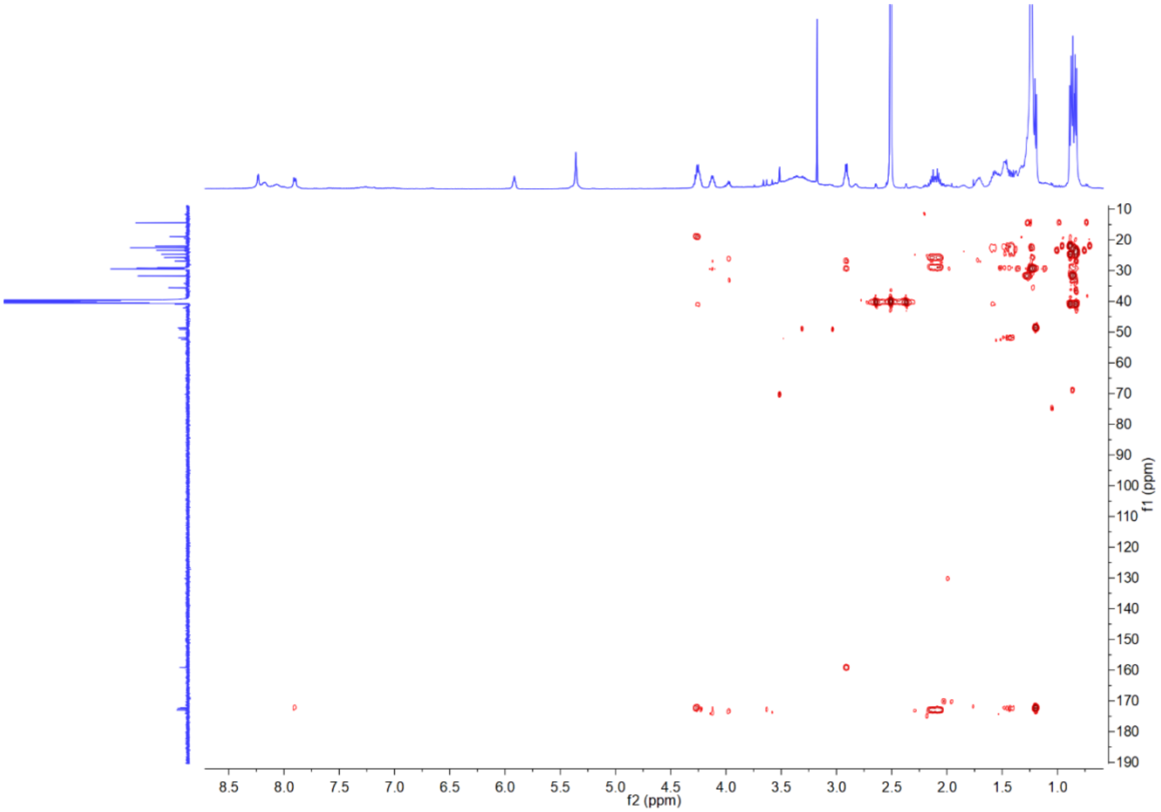


**Supplementary Note Fig. 38 |** HMBC spectrum of lipocitide B (**18**) in DMSO-$d_6$.

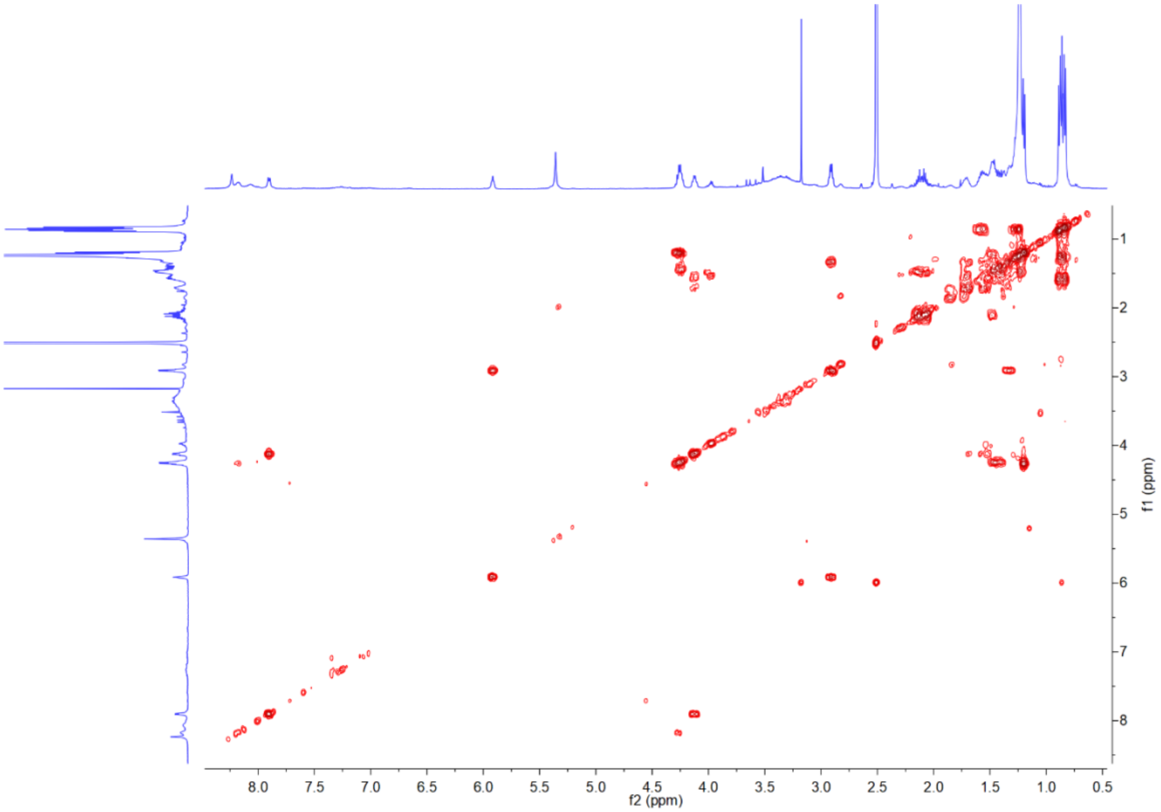


**Supplementary Note Fig. 39 |** $^1H$-$^1H$ COSY spectrum of lipocitide B (**18**) in DMSO-$d_6$.

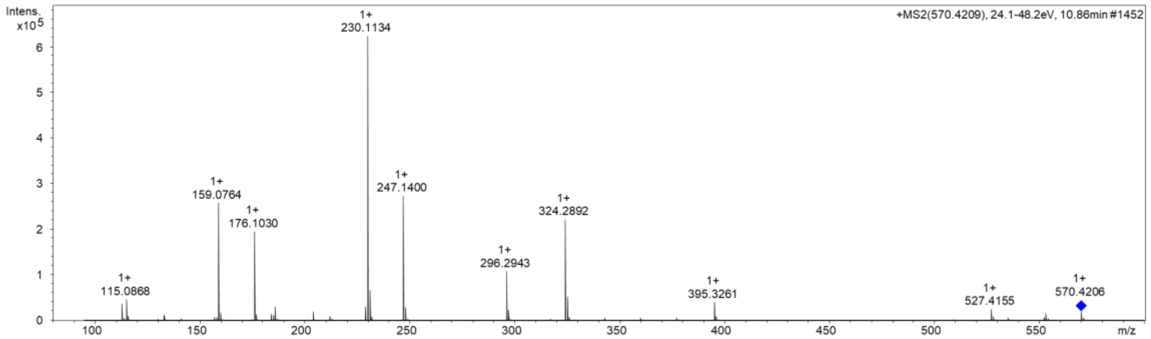


**Supplementary Note Fig. 40 |** HR-ESI-MS of lipocitide B (**18**).

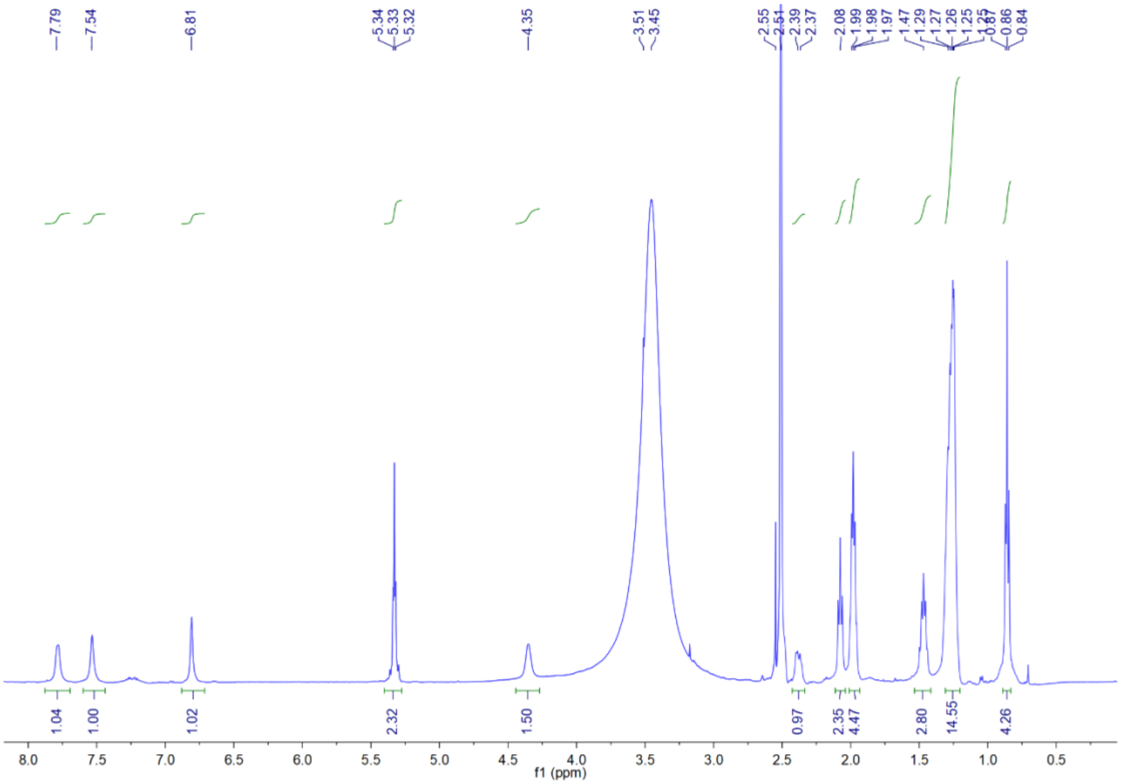


**Supplementary Note Fig. 41 |** $^{1}$H NMR spectrum of N-(ω-7-myristoyl)-D-asparagine (**19**) in DMSO-$d_6$.

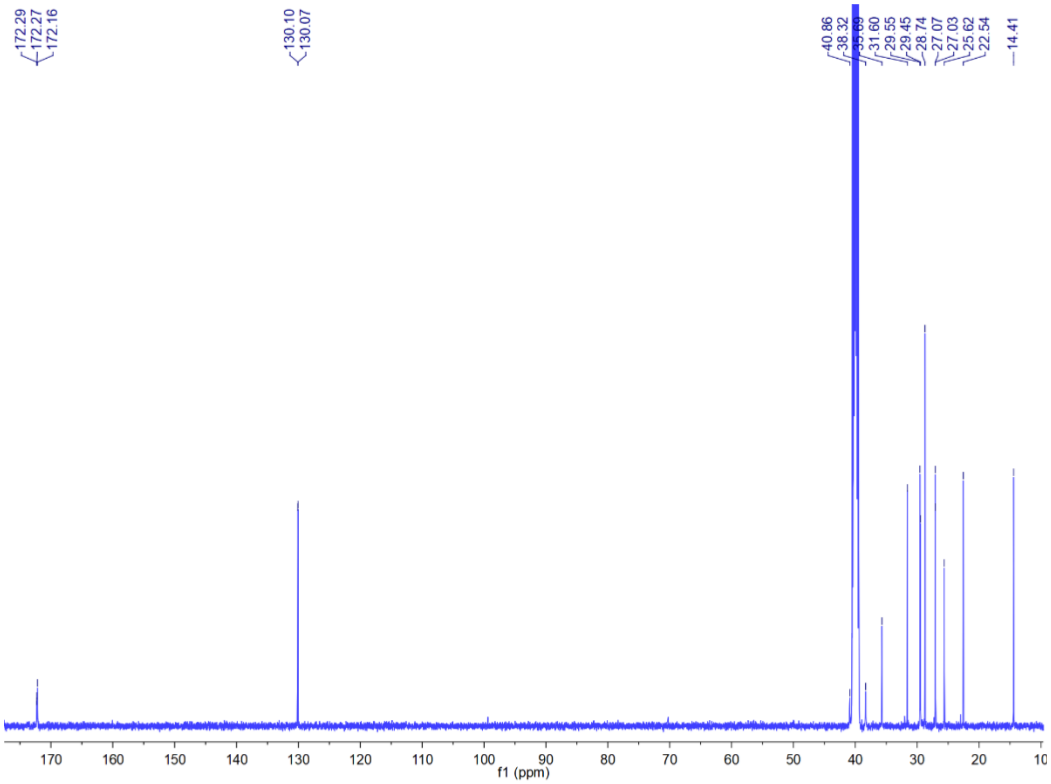


**Supplementary Note Fig. 42 |** $^{13}$C NMR spectrum of N-(ω-7-myristoyl)-D-asparagine (**19**) in DMSO-$d_6$.

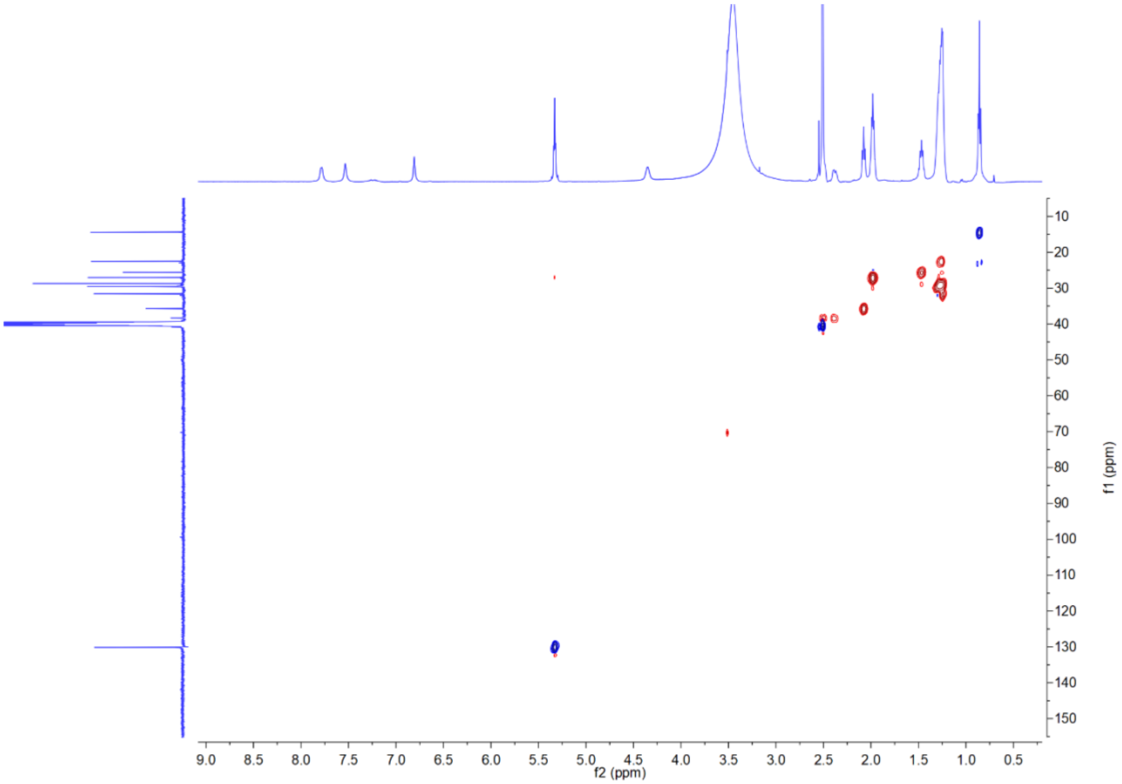


**Supplementary Note Fig. 43 |** HSQC spectrum of N-(ω-7-myristoyl)-D-asparagine (**19**) in DMSO-$d_6$.

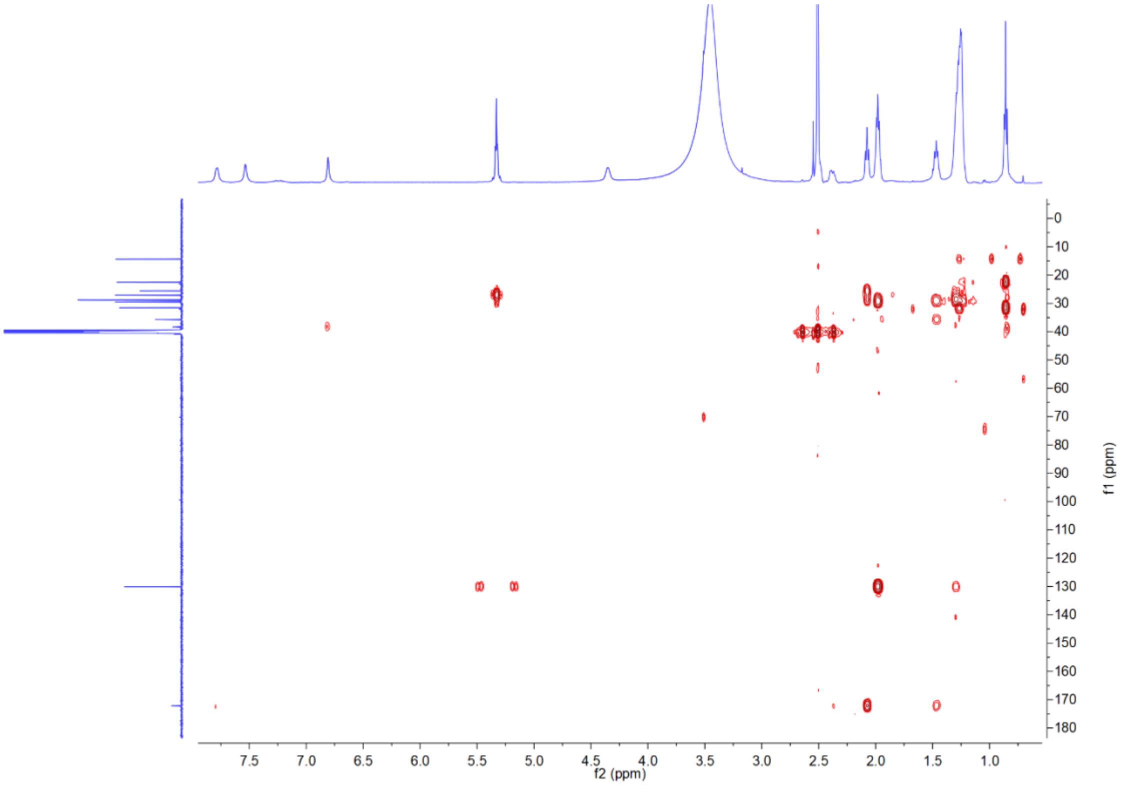


**Supplementary Note Fig. 44 |** HMBC spectrum of N-(ω-7-myristoyl)-D-asparagine (**19**) in DMSO-$d_6$.

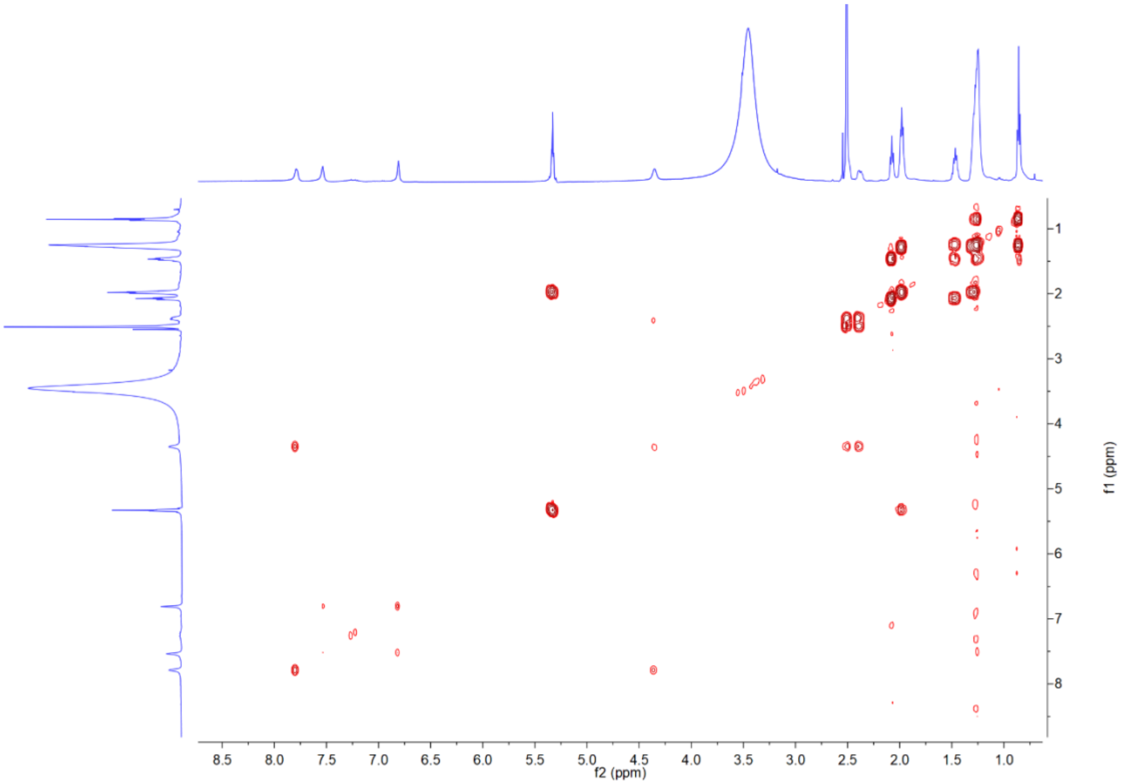


**Supplementary Note Fig. 45 |** $^1$H-$^1$H COSY spectrum of N-(ω-7-myristoyl)-D-asparagine (**19**) in DMSO-$d_6$.

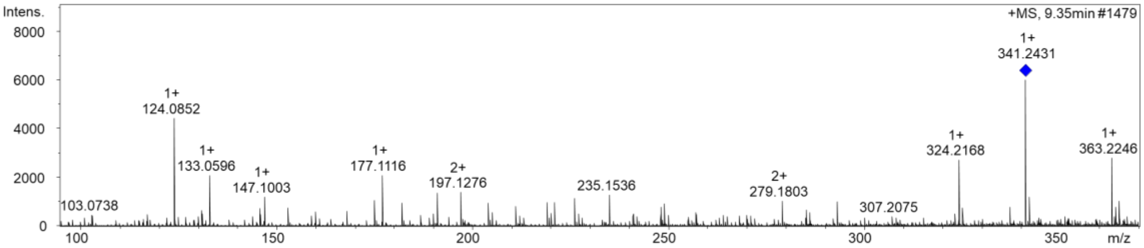


**Supplementary Note Fig. 46 |** HR-ESI-MS of N-(ω-7-myristoyl)-D-asparagine (**19**).

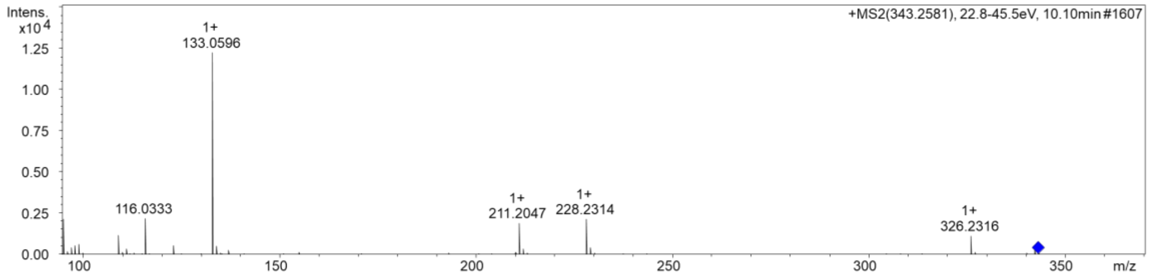


**Supplementary Note Fig. 47 |** HR-ESI-MS of N-myristoyl-D-asparagine (**20**).

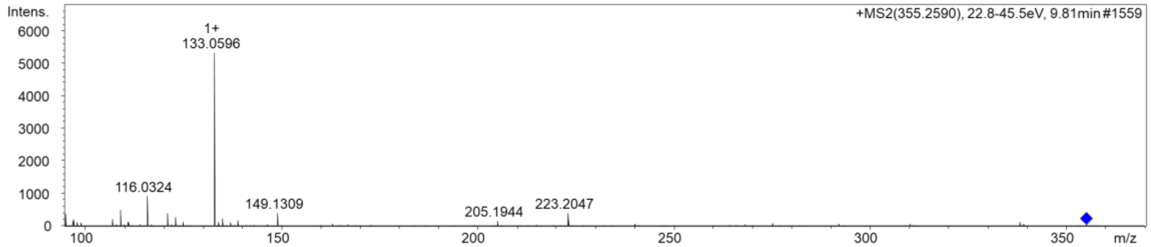


**Supplementary Note Fig. 48 |** HR-ESI-MS of N-(13-methyl-ω-7-myristoyl)-D-asparagine (**21**).

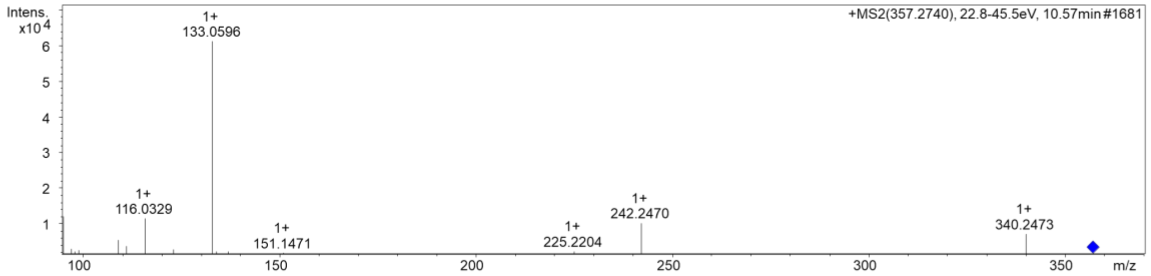


**Supplementary Note Fig. 49 |** HR-ESI-MS of N-(13-methylmyristoyl)-D-asparagine (**22**).

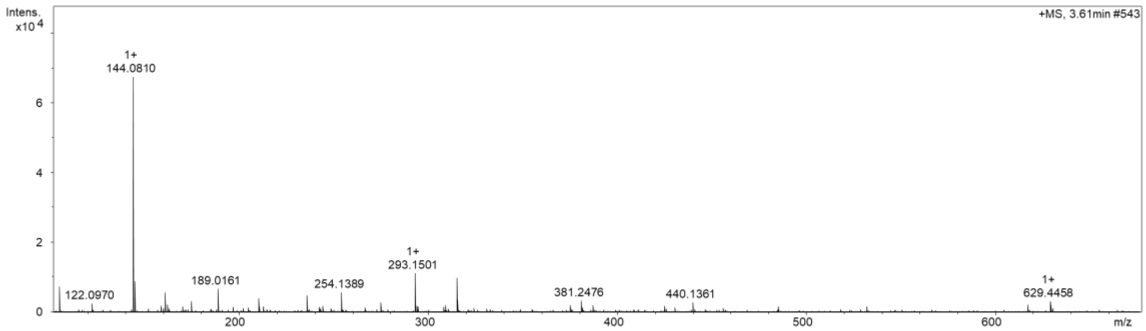


**Supplementary Note Fig. 50 |** HR-ESI-MS of rhabdobranin (**23**).

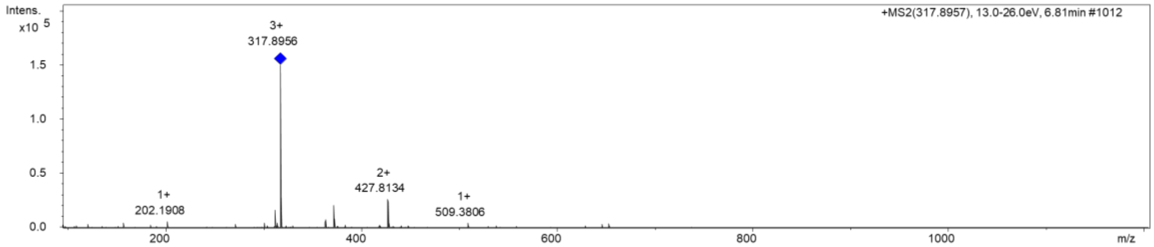


**Supplementary Note Fig. 51 |** HR-ESI-MS of pre-rhabdobranin A (**24**).

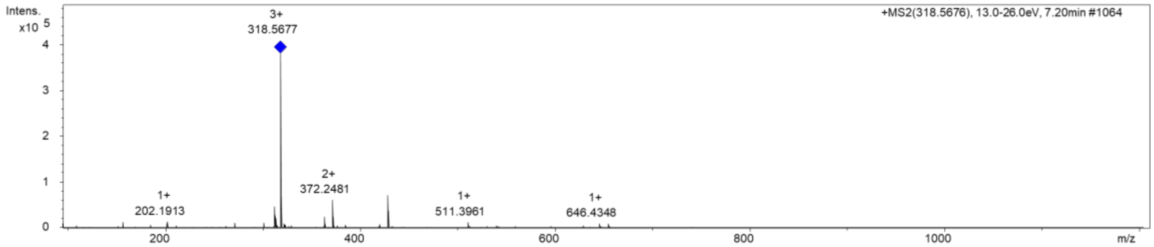


**Supplementary Note Fig. 52 |** HR-ESI-MS of pre-rhabdobranin B (**25**).

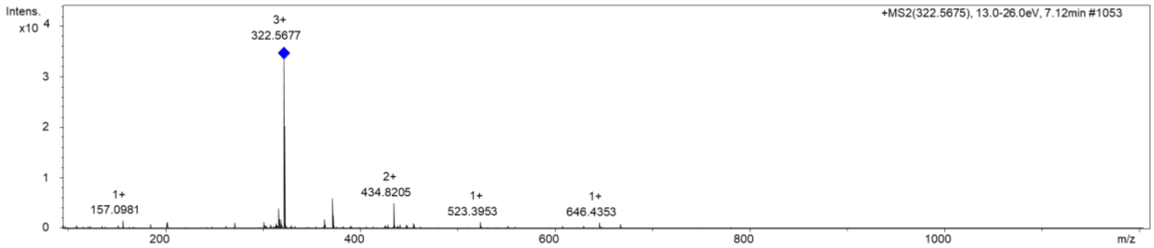


**Supplementary Note Fig. 53 |** HR-ESI-MS of pre-rhabdobranin C (**26**).

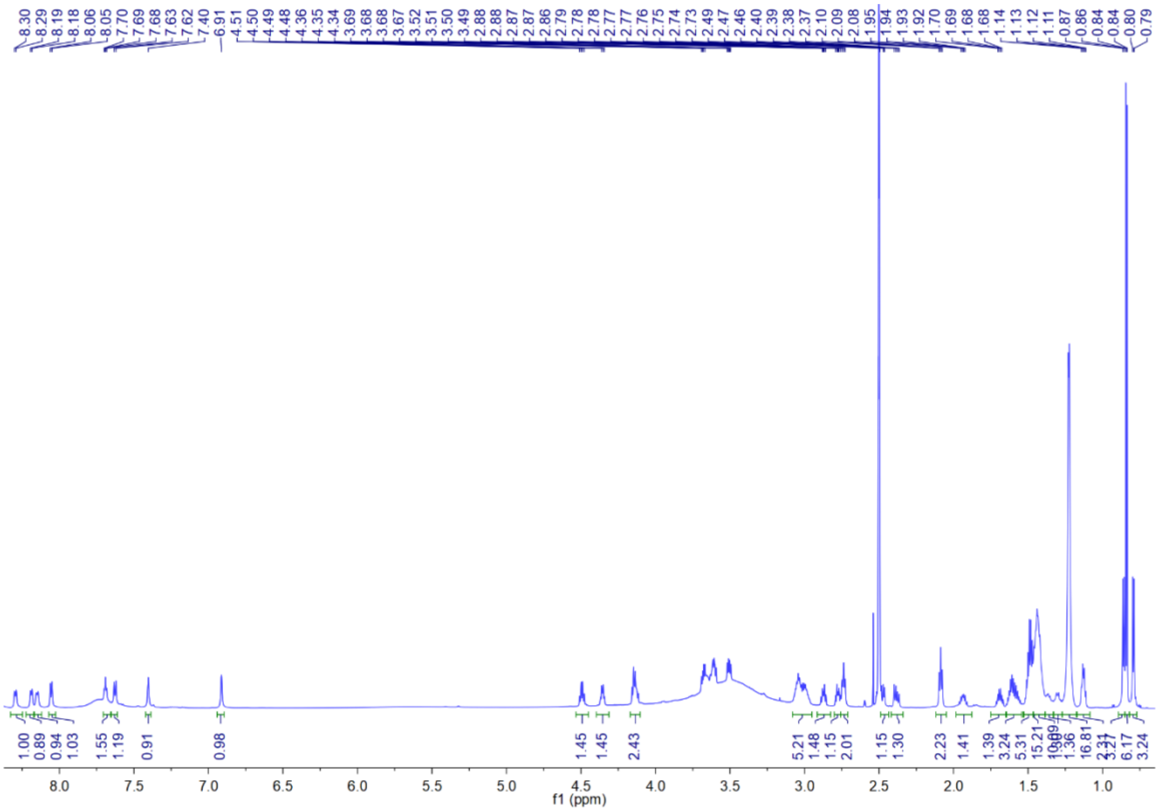

**Supplementary Note Fig. 54 |** $^{1}H$ NMR spectrum of pre-rhabdobranin D (**27**) in DMSO-$d_6$.

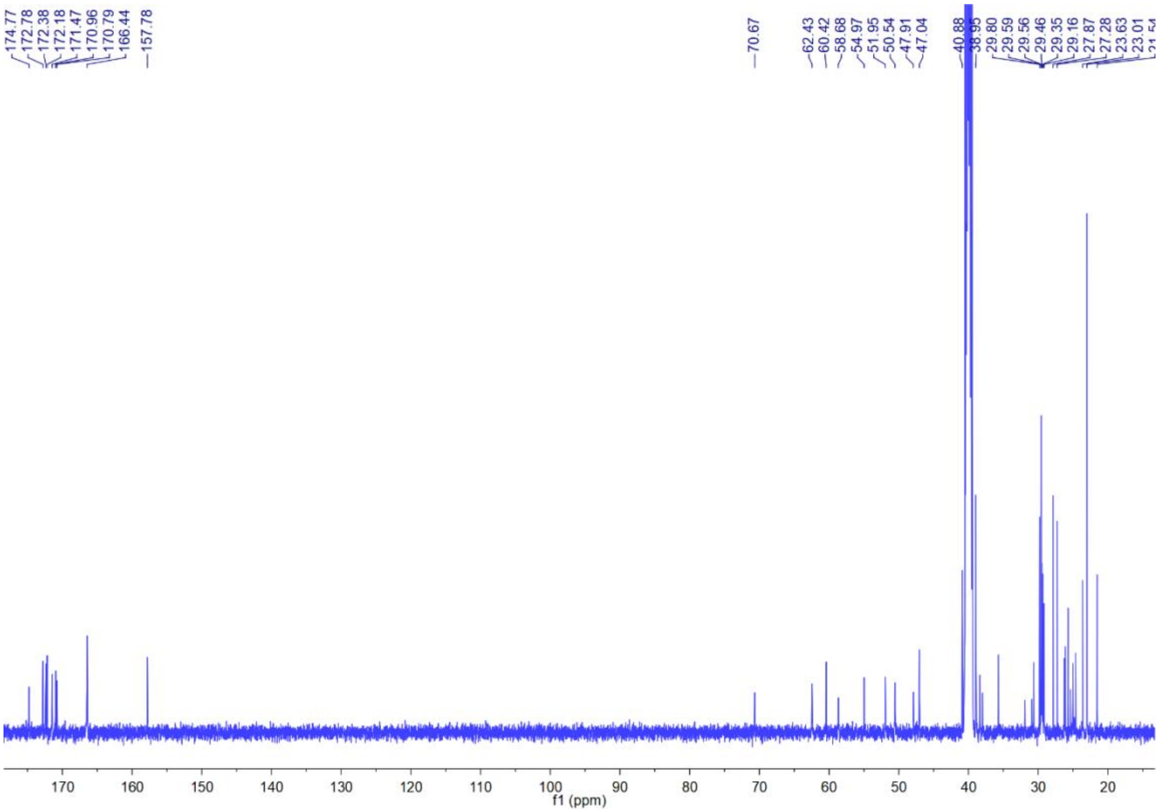

**Supplementary Note Fig. 55 |** $^{13}C$ NMR spectrum of pre-rhabdobranin D (**27**) in DMSO-$d_6$.

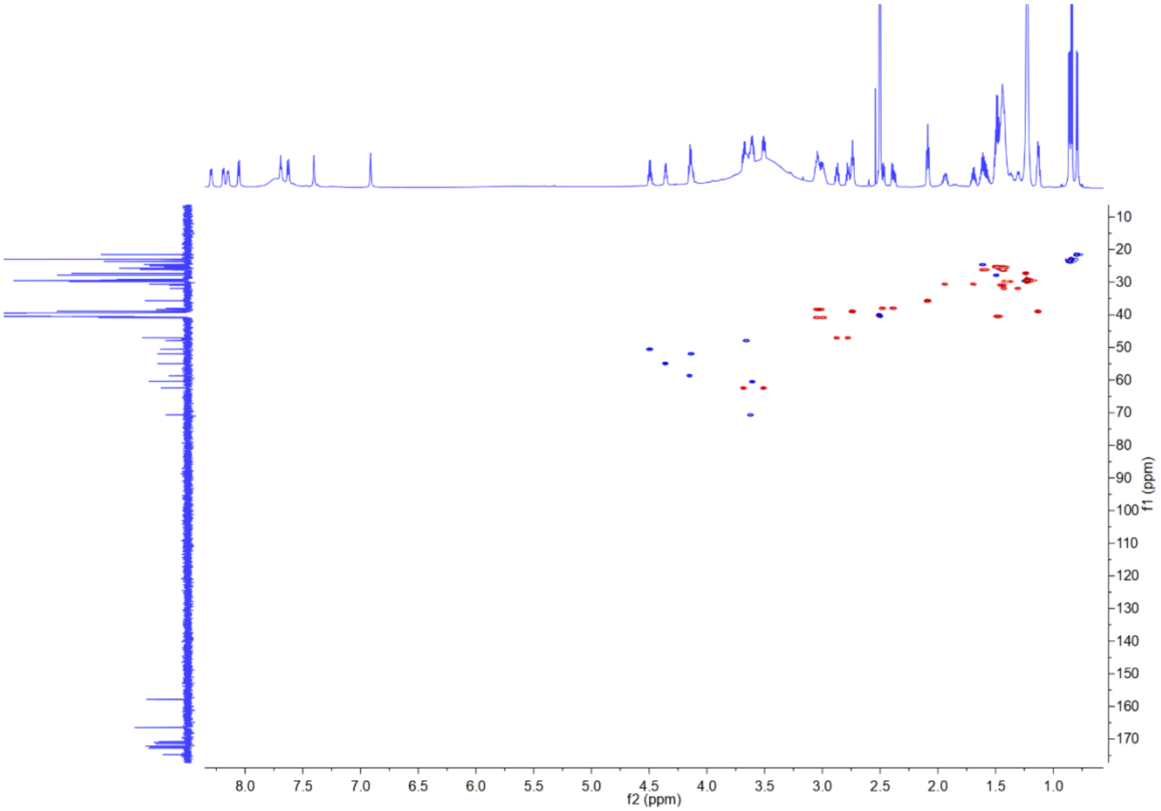


**Supplementary Note Fig. 56 |** HSQC spectrum of pre-rhabdobranin D (**27**) in DMSO-$d_6$.

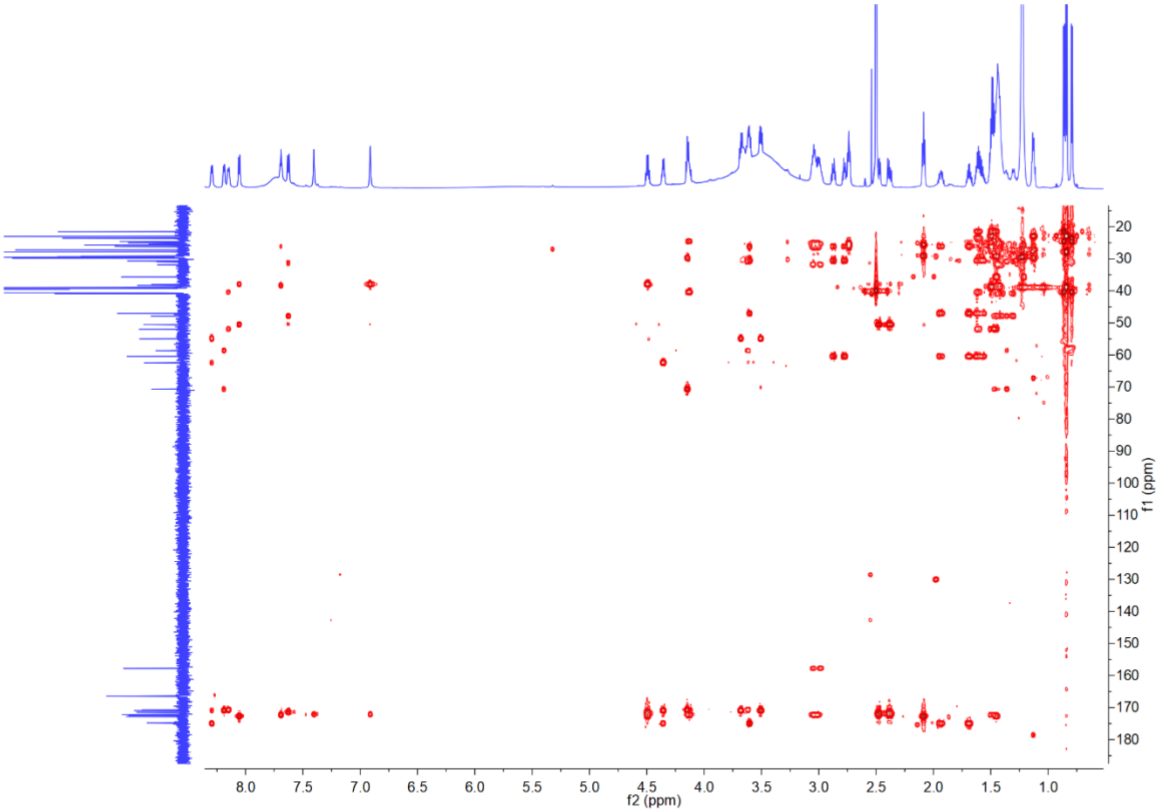


**Supplementary Note Fig. 57 |** HMBC spectrum of pre-rhabdobranin D (**27**) in DMSO-$d_6$.

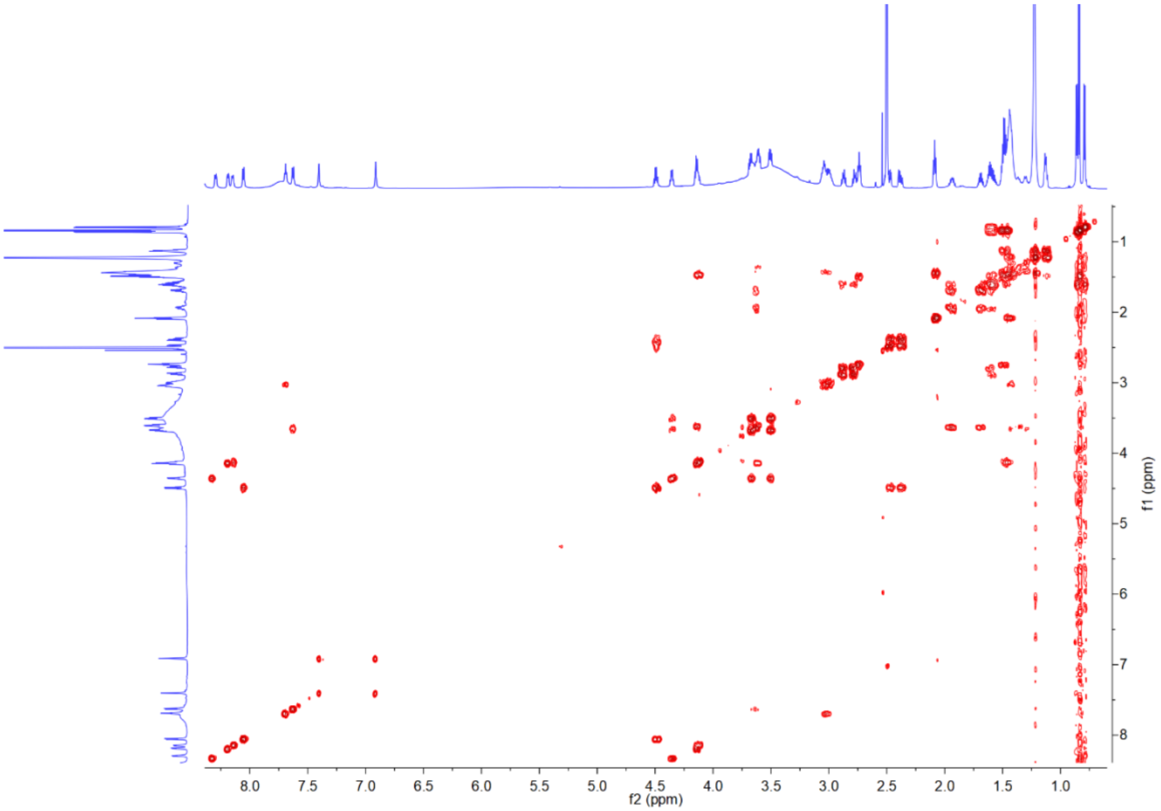


**Supplementary Note Fig. 58 |** $^{1}H$-$^{1}H$ COSY spectrum of pre-rhabdobranin D (**27**) in DMSO-$d_6$.

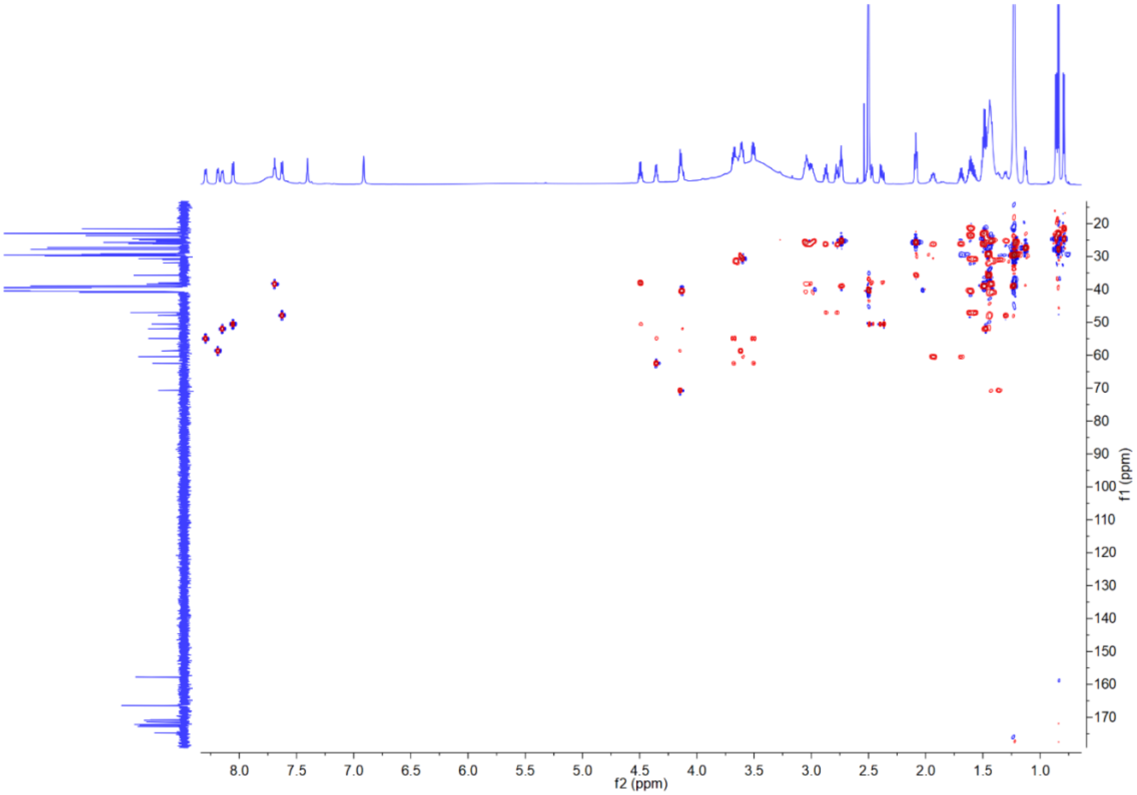


**Supplementary Note Fig. 59 |** HMQC-COSY spectrum of pre-rhabdobranin D (**27**) in DMSO-$d_6$.

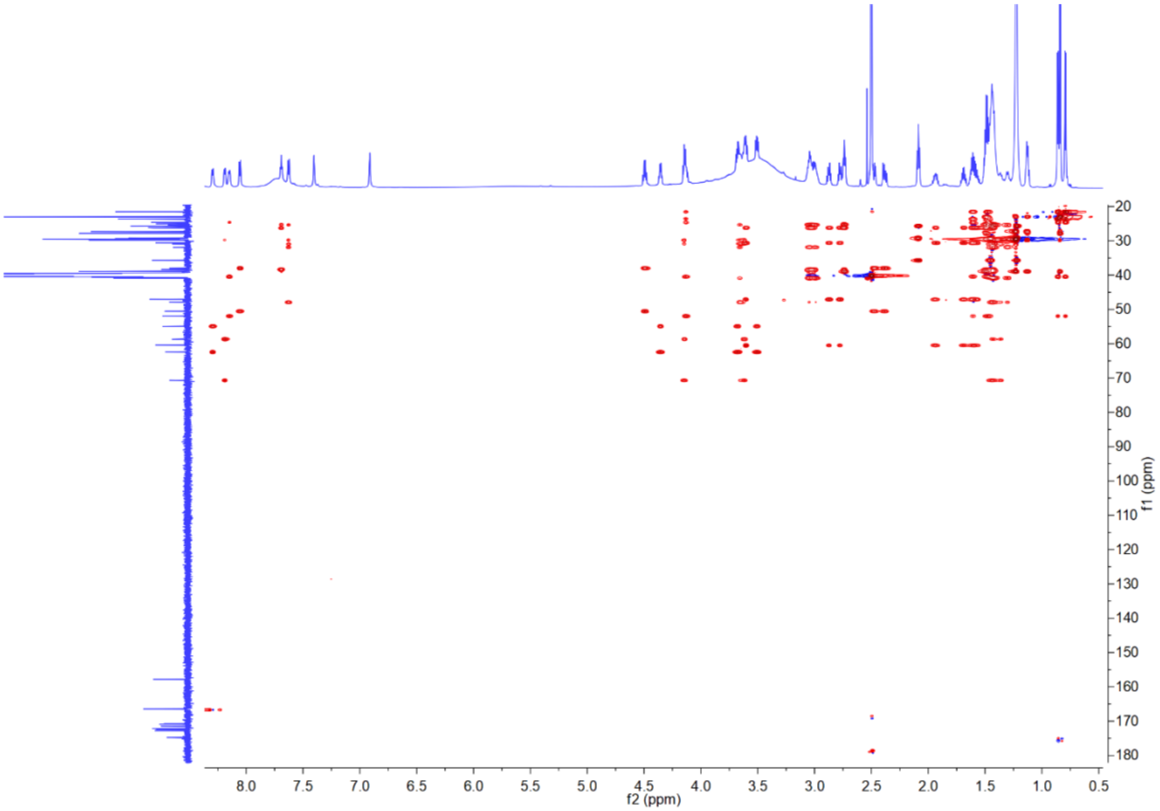


**Supplementary Note Fig. 60 |** HSQC-TOCSY spectrum of pre-rhabdobranin D (**27**) in DMSO-*d*$_6$.

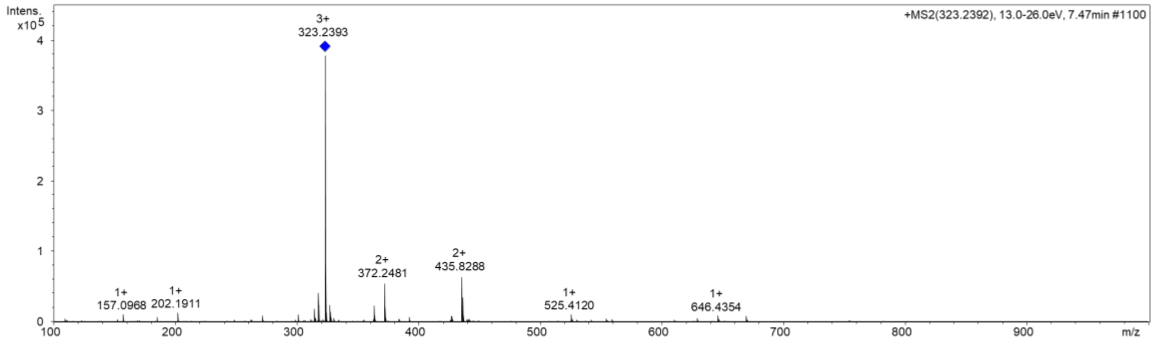


**Supplementary Note Fig. 61 |** HR-ESI-MS of pre-rhabdobranin D (**27**).

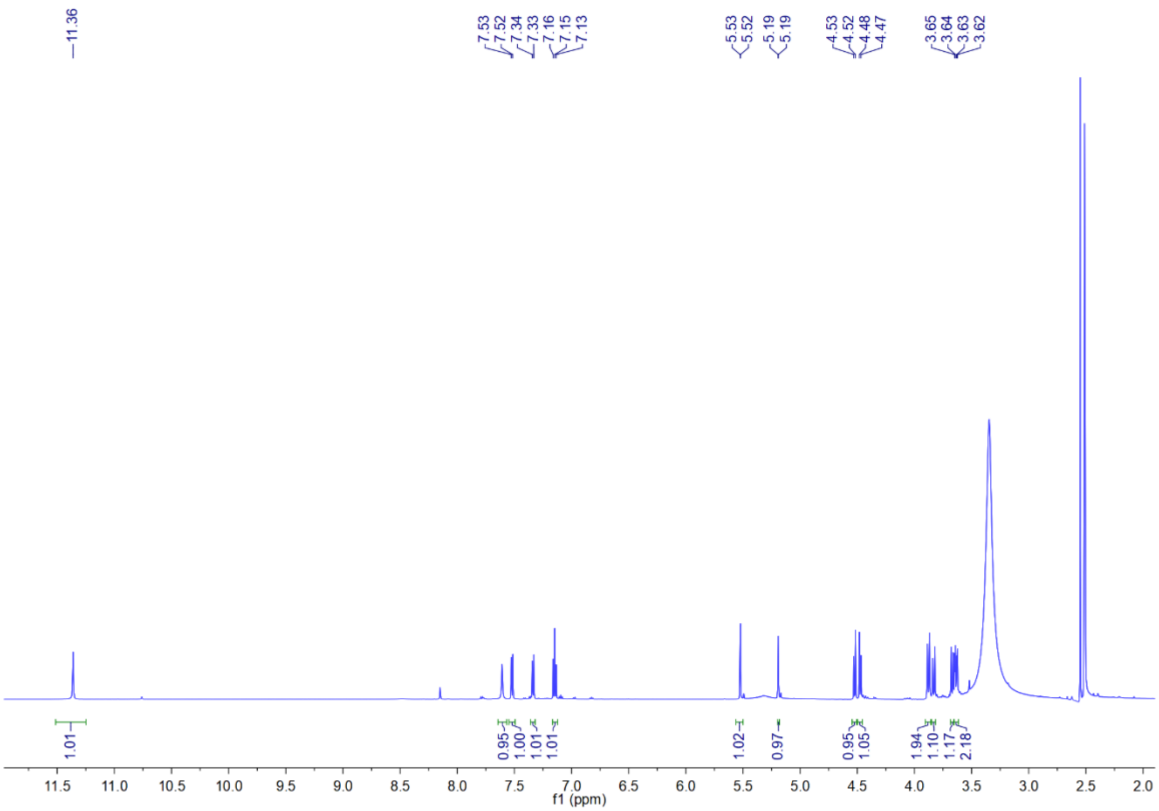


**Supplementary Note Fig. 62 |** $^{1}H$ NMR spectrum of benzobactin A (**28**) in DMSO-$d_6$.

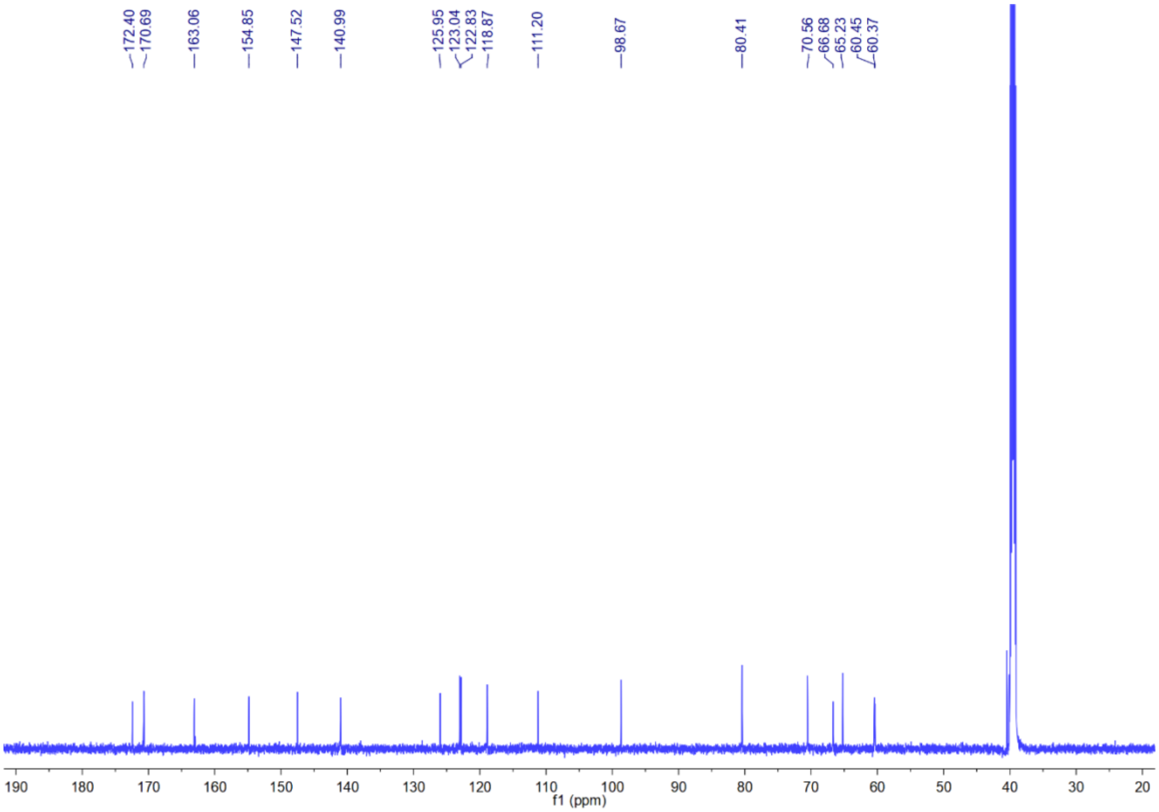


**Supplementary Note Fig. 63 |** $^{13}C$ NMR spectrum of benzobactin A (**28**) in DMSO-$d_6$.

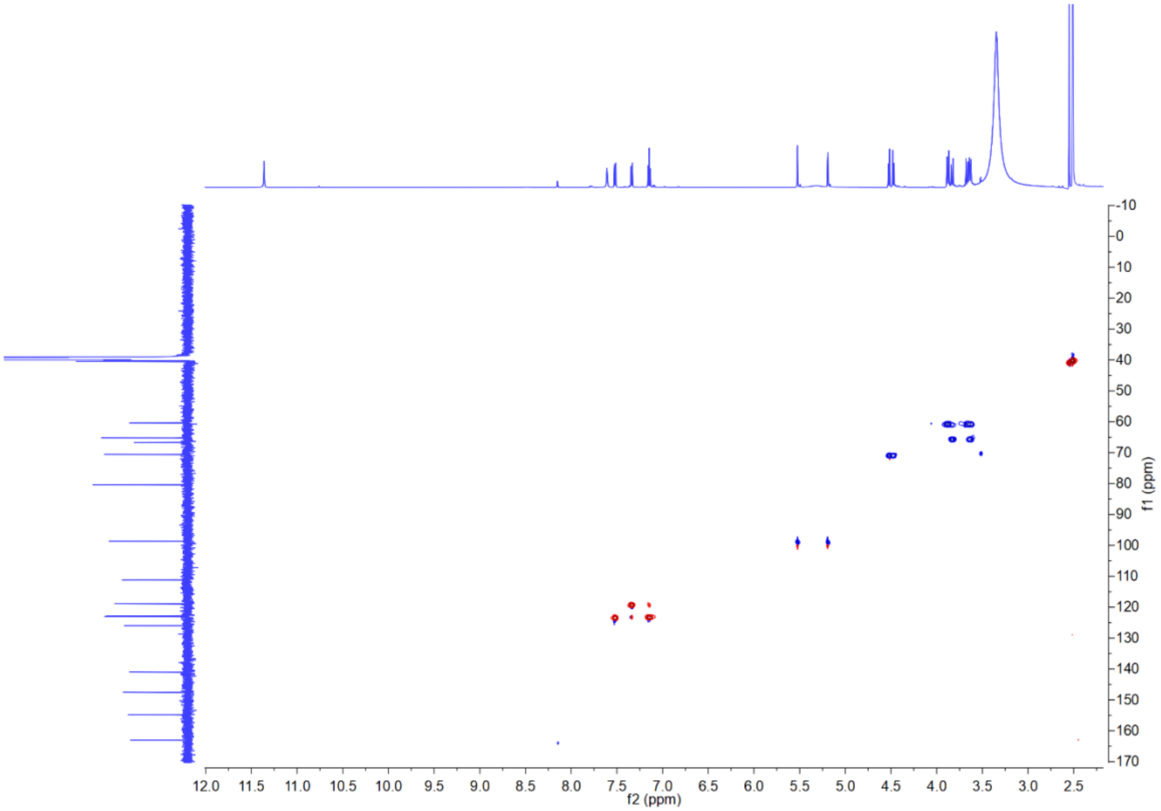


**Supplementary Note Fig. 64 |** HSQC spectrum of benzobactin A (**28**) in DMSO-*d*6.

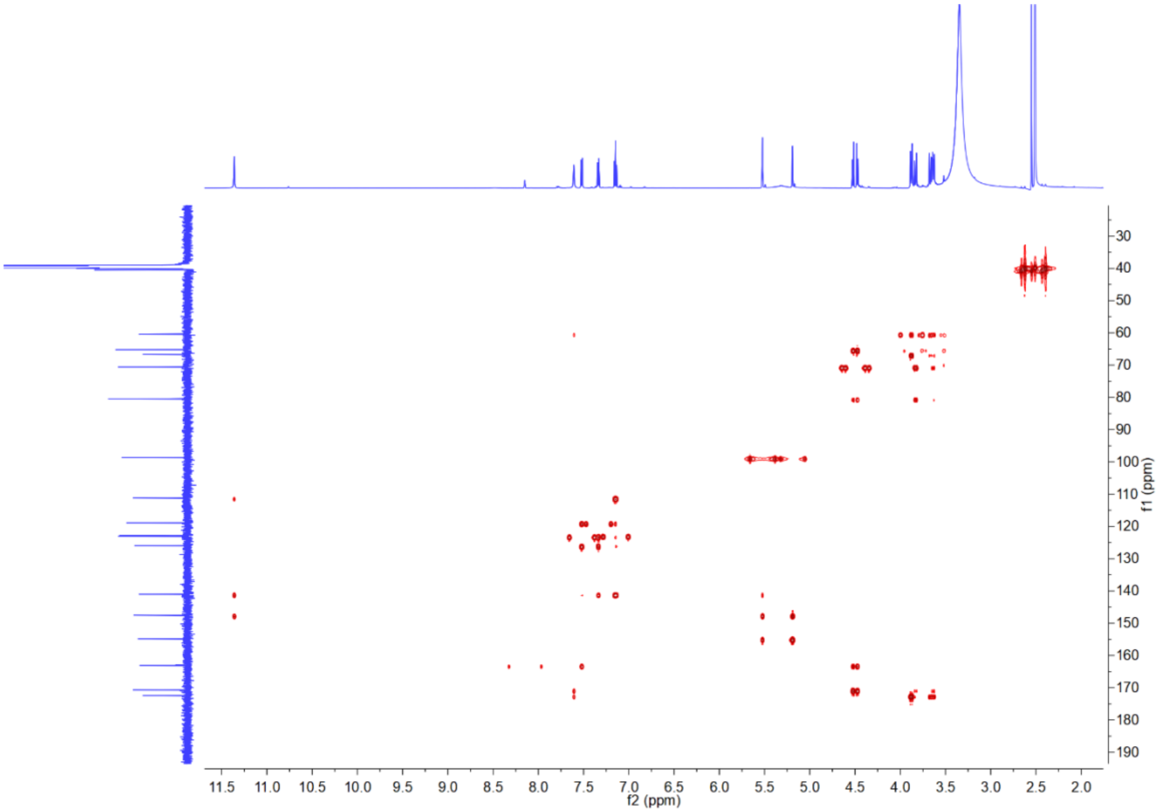


**Supplementary Note Fig. 65 |** HMBC spectrum of benzobactin A (**28**) in DMSO-*d*6.

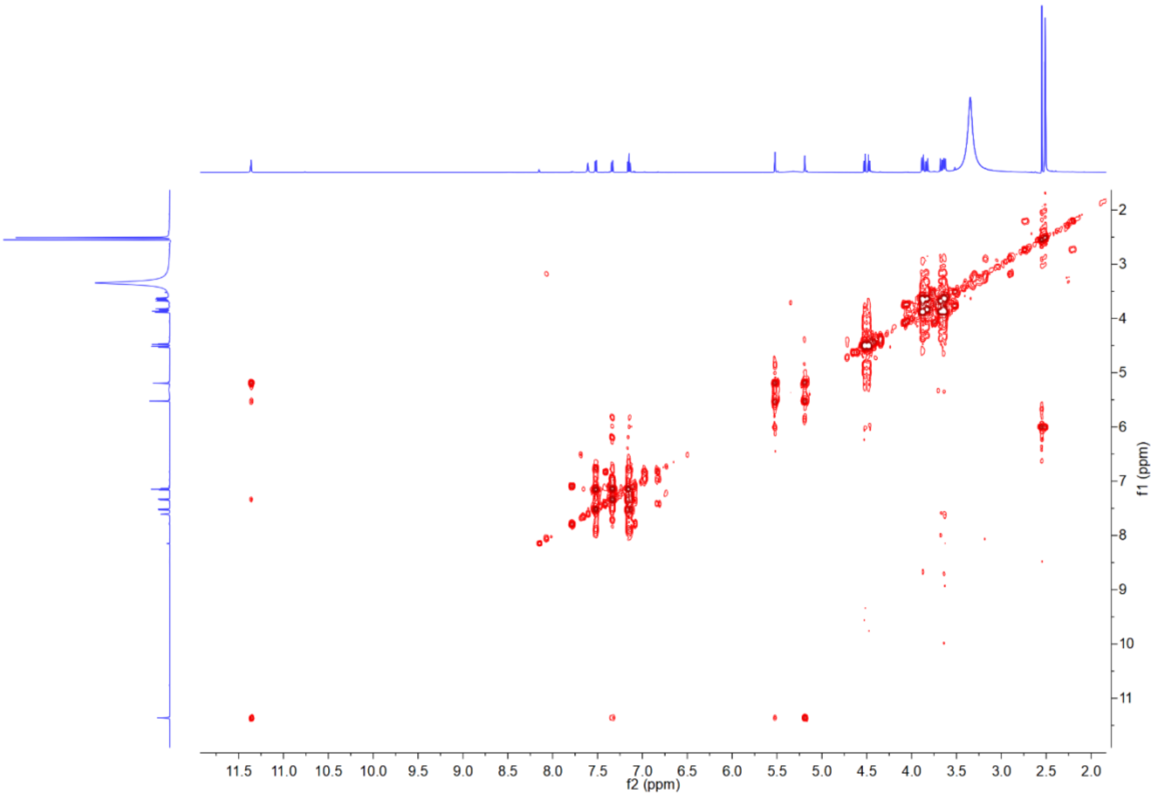


**Supplementary Note Fig. 66 |** $^{1}$H-$^{1}$H COSY spectrum of benzobactin A (**28**) in DMSO-$d_6$.

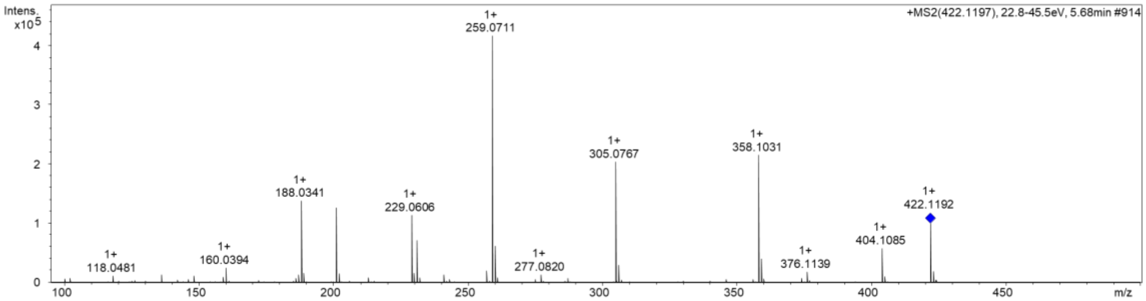


**Supplementary Note Fig. 67 |** HR-ESI-MS of benzobactin A (**28**).

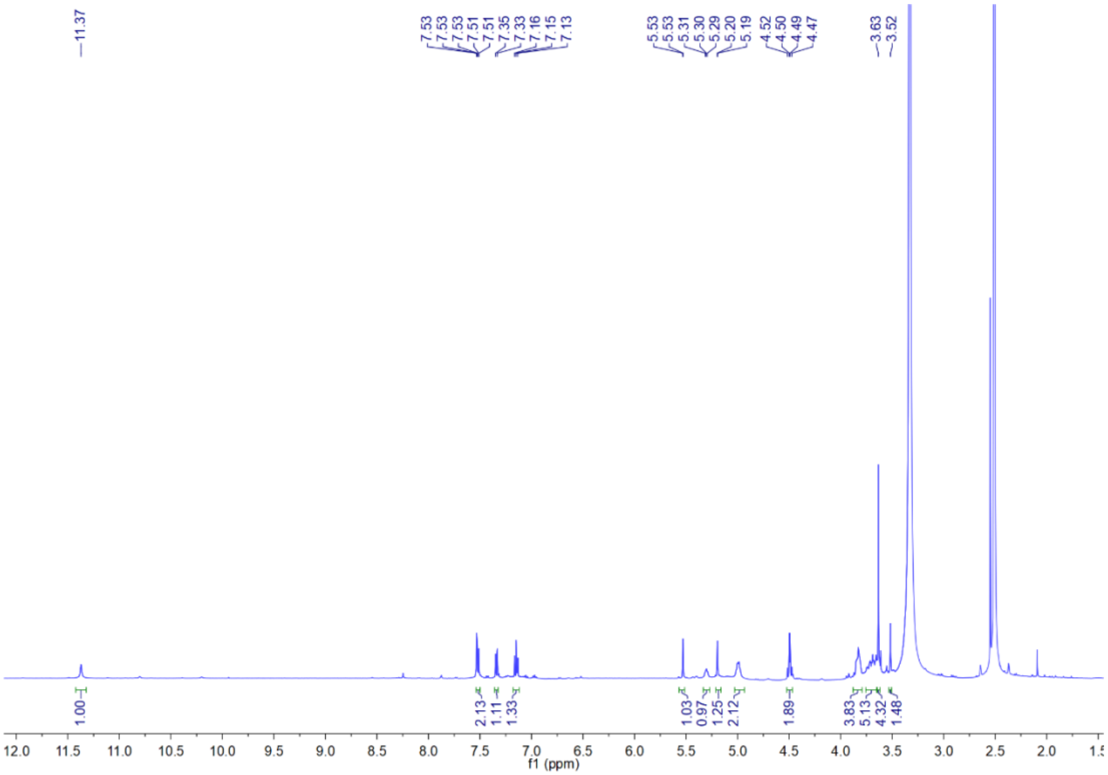

**Supplementary Note Fig. 68 |** $^{1}$H NMR spectrum of benzobactin A methyl ester (**29**) in DMSO-*d*$_6$.

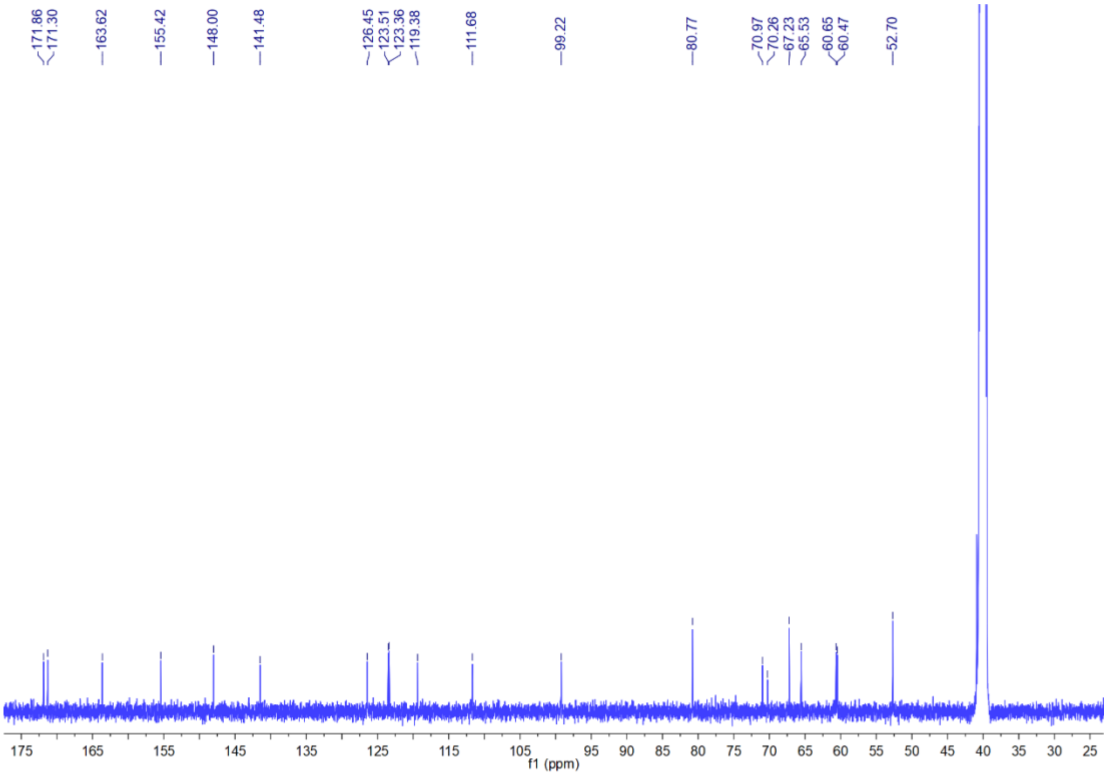

**Supplementary Note Fig. 69 |** $^{13}$C NMR spectrum of benzobactin A methyl ester (**29**) in DMSO-*d*$_6$.

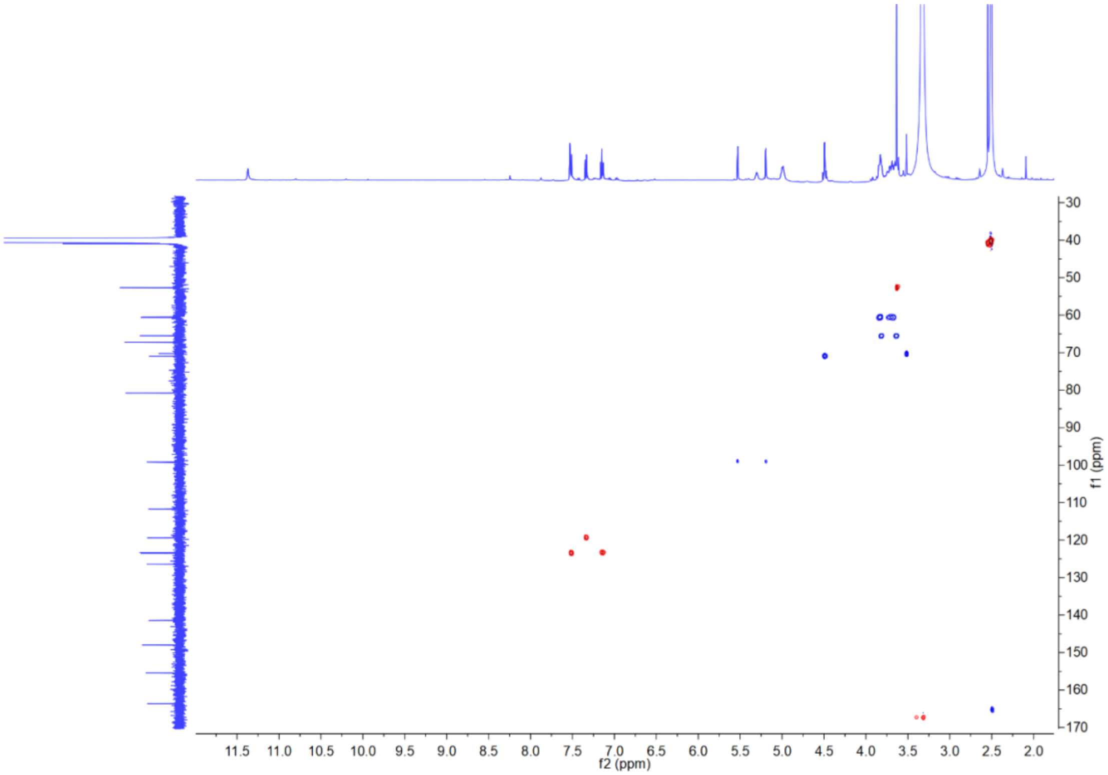

**Supplementary Note Fig. 70 |** HSQC spectrum of benzobactin A methyl ester (**29**) in DMSO-$d_6$.

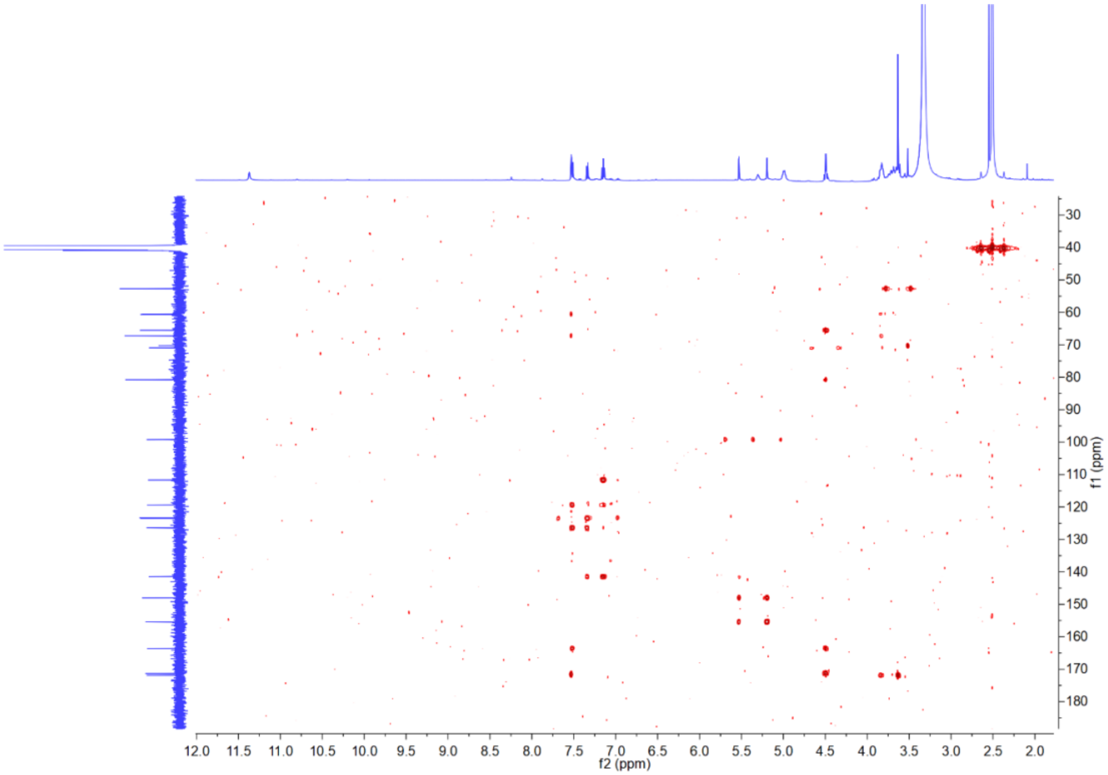

**Supplementary Note Fig. 71 |** HMBC spectrum of benzobactin A methyl ester (**29**) in DMSO-$d_6$.

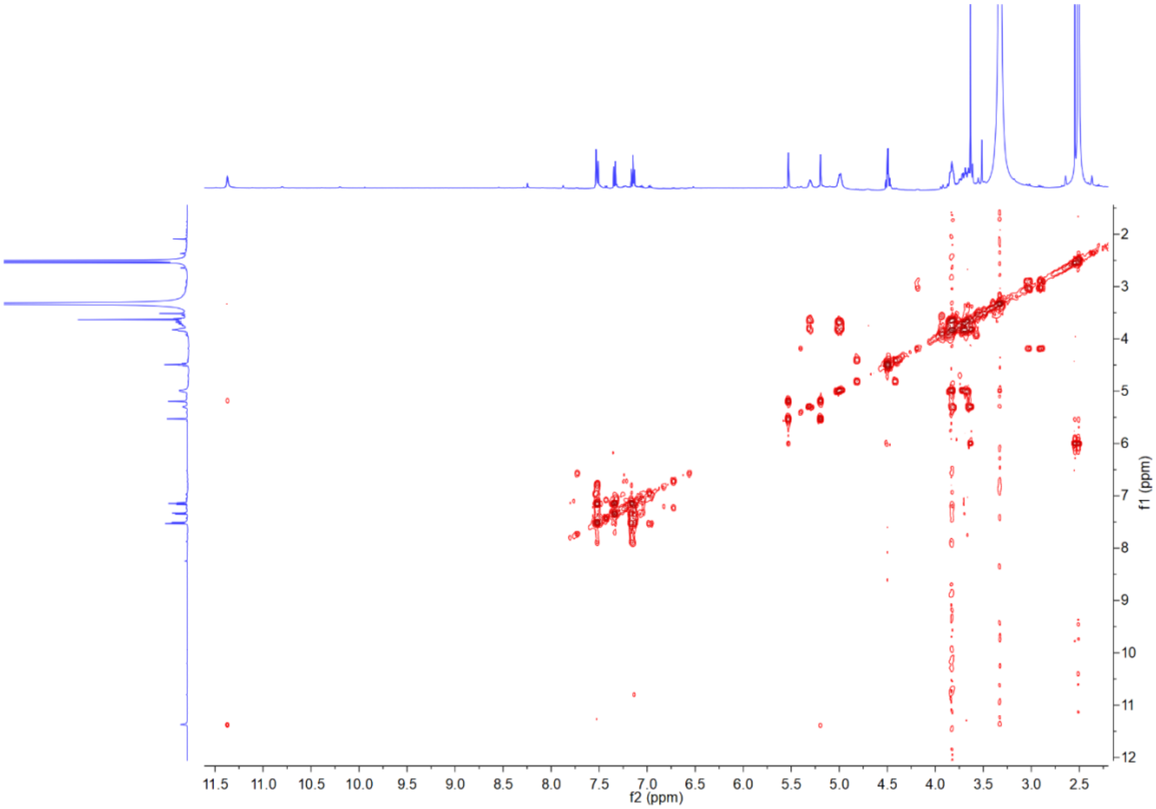

**Supplementary Note Fig. 72 |** $^{1}H$-$^{1}H$ COSY spectrum of benzobactin A methyl ester (**29**) in DMSO-$d_6$.

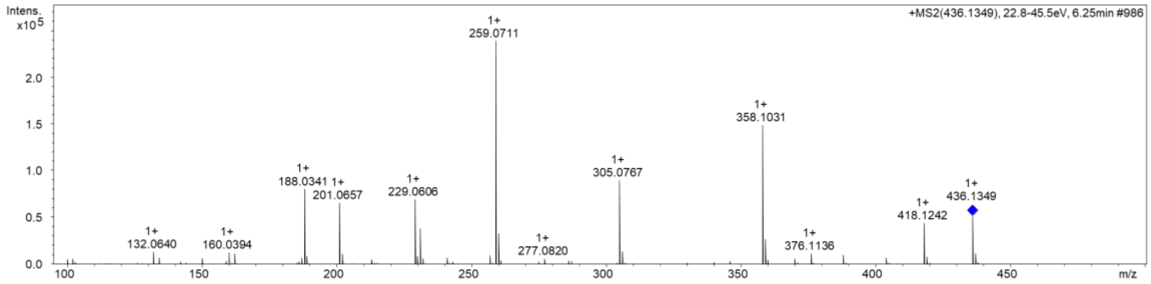


**Supplementary Note Fig. 73 |** HR-ESI-MS of benzobactin A methyl ester (**29**).
